# Density-dependent habitat selection alters drivers of population distribution in northern Yellowstone elk

**DOI:** 10.1101/2022.07.12.499670

**Authors:** Brian J. Smith, Daniel R. MacNulty, Daniel R. Stahler, Douglas W. Smith, Tal Avgar

**Affiliations:** Department of Wildland Resources and Ecology Center, Utah State University, _Logan_, UT 84322, USA; Yellowstone Center for Resources, National Park Service, Yellowstone National Park, Wyoming 82190, USA; Biodiversity Pathways Ltd., P.O.B 63 Mill Bay, BC, V0R 2P0, Canada

**Keywords:** cougar, density dependence, food-safety tradeoff, habitat selection, ideal free distribution, predation risk, predator-prey interactions, RSF, spatial distribution, wolf

## Abstract

Although it is well established that density dependence drives changes in organismal abundance over time, relatively little is known about how density dependence affects variation in abundance over space. We tested the hypothesis that spatial tradeoffs between food and safety can change the drivers of population distribution, caused by opposing patterns of density-dependent habitat selection (DDHS) that are predicted by the multidimensional ideal free distribution. We addressed this using winter aerial survey data of northern Yellowstone elk (*Cervus canadensis*) spanning four decades. Supporting our hypothesis, we found positive DDHS for food (herbaceous biomass) and negative DDHS for safety (openness and roughness), such that the primary driver of habitat selection switched from food to safety as elk density decreased from 9.3 to 2.0 elk/km^2^. Our results demonstrate how population density can drive landscape-level shifts in population distribution, confounding habitat selection inference and prediction, and potentially affecting community-level interactions.

## Introduction

Density dependence is a pervasive ecological process, and incorporating it into models of population abundance is critical for understanding population dynamics and informing management (Abadi *et al*. 2012; Guthery & Shaw 2013). Whereas much is known about how density dependence affects variation in abundance over time, relatively little is known about how it affects abundance over space. This is a particular problem for the study of habitat selection, where density-dependent habitat selection (DDHS) is a foundational assumption (Rosenzweig 1981) that is rarely tested and often ignored (Avgar et al 2020). This gap is significant because unmeasured density-dependent variation in habitat selection may limit the accuracy of empirical models for inferring drivers of fitness (e.g., food vs. safety) and predicting spatial distribution and abundance (Boyce & McDonald 1999; Matthiopoulos *et al*. 2015, 2019).

The expectation of DDHS arises from optimal foraging theory (MacArthur & Pianka 1966) via the ideal free distribution (IFD; Fretwell & Lucas 1969). The IFD postulates that a population’s density influences the fitness benefits that individuals receive from a habitat (Morris 1987), and that individuals use habitats in a way that equalizes fitness across occupied habitats (Fretwell & Lucas 1969; Křivan *et al*. 2008). According to IFD, as population density increases, individuals occupy progressively lower-quality habitats (“spillover”), resulting in “negative DDHS” – the strength of selection for high-quality habitat decreases with density (Rosenzweig 1991; Morris 2003). Alternatively, fitness in certain habitats can increase with population density (Stephens & Sutherland 1999), leading to an increase in selection strength for these habitats – “positive DDHS” (Morris 2002). Positive DDHS manifests as habitat switching, where individuals leave habitats associated with high fitness at a low density, shifting into habitats associated with high fitness at a high density (Greene & Stamps 2001).

Fitness is never determined by a single environmental driver; thus, habitat selection reflects a balancing act along multiple dimensions and scales, each with its own context-dependent relative contribution to overall fitness (multidimensional IFD sensu Avgar *et al*. 2020). Population density is one of these contexts, and whether fitness in a habitat decreases or increases with density varies across different habitat dimensions. For example, fitness typically decreases with density due to reduced per-capita food acquisition (Le Bourlot *et al*. 2014), whereas it typically increases with density due to increased safety (i.e., reduced per-capita predation risk; reviewed by Lehtonen & Jaatinen 2016). If food and safety are spatially independent, we expect negative DDHS along both dimensions: at low density, individuals select habitats with more food and safety, but as density increases, individuals spill into habitats with less food or safety. Conversely, if food and safety are negatively correlated in space, individuals select habitats with more food *or* safety. Thus, we expect negative DDHS along the safety dimension and positive DDHS along the food dimension; as density increases, safety in numbers increases, and simultaneously, increasing intraspecific competition makes food more limiting to fitness, hence a stronger driver of habitat selection. (Avgar *et al*. 2020).

Despite the theoretical and practical importance of DDHS, empirical understanding of DDHS and its effects on inference and prediction from habitat selection models is underdeveloped (Avgar *et al*. 2020). This is especially true in free-living systems (McLoughlin *et al*. 2010), which involve complex trade-offs (e.g., food for safety) that are often missing in experimental systems (Lima & Dill 1990), habitat selection at multiple scales (Johnson 1980), and multiple interacting species. To fill this gap, we tested the hypothesis that tradeoffs between food and safety generate positive DDHS for food resources and negative DDHS for safe habitats, such that the drivers of habitat selection switch from food to safety as density decreases. We did so by constructing and applying a population-level habitat selection model to winter aerial-survey data of northern Yellowstone elk collected over 16 years spanning 4 decades. Our findings demonstrate how ignoring DDHS can confound both ecological inference and prediction. We provide novel evidence of multidimensional IFD and of DDHS as an important driver of habitat selection and spatial distribution in a free-living system.

## Methods

### Study area

Our study occurred in the winter range of the northern Yellowstone elk population (Houston 1982; Lemke & Mack 1998). We expanded the area previously used to define the winter range (e.g., Tallian *et al*. 2017) to include adjacent areas where elk were also occasionally counted (Fig. S1). We believe the modified polygon better captures the full extent used by this population. Our study area encompassed 1900 km^2^, compared with the 1520 km^2^ previous authors have cited. Approximately two-thirds of the winter range falls within the boundaries of Yellowstone National Park (YNP), with the remaining one-third to the north in the state of Montana. Elevations range between 1,500 and 3,000 m, and the area experiences long, cold winters (Houston 1982). Northern Yellowstone elk migrate from high-elevation summer ranges, often in the interior of YNP, to the lower elevation winter range, where they are found from December – April (Houston 1982; White *et al*. 2010).

Wolves (*Canis lupus*) and cougars (*Puma concolor*) are the two main predators of northern Yellowstone elk during winter (Kohl *et al*. 2019). Elk comprised 96% of the wolf diet in winter from 1995 – 2009 (Metz *et al*. 2012) and 75% of the cougar diet from 1998 – 2005 (Ruth *et al*. 2019). Alternative prey for wolves and cougars in the system include bison (*Bison bison*), deer (*Odocoileus hemionus*, *O. virginianus*), moose (*Alces alces*), pronghorn (*Antilocapra americana*), and bighorn sheep (*Ovis canadensis*) (Metz *et al*. 2012; Ruth *et al*. 2019).

### Data collection

A timeseries of winter counts of northern Yellowstone elk extends to the 1920s, and the population has fluctuated with various management practices and climatic conditions throughout it (MacNulty *et al*. 2020). Since 1988, these data have included georeferenced locations of elk groups in many years, allowing us to estimate habitat selection. Elk were counted via aerial, fixed-wing surveys designed to provide a full population census. Surveys took place mainly between 08:00 and 12:00 over 1 – 4 days between late December and March of each winter (Appendix S1). In some years, the count did not occur or georeferenced group data were only available within YNP. We used a state-space model (see Appendix S1 in Tallian *et al*. 2017) to interpolate total abundance in years with incomplete data so that we could predict using our fitted habitat selection model. We used the partial georeferenced data for validation (see *Model evaluation*). Counts reached an all-time high in 1994 near 20,000 elk and a low of less than 4,000 in 2013 (MacNulty *et al*. 2020). Thus, this timeseries of georeferenced counts provided information about elk distribution across a wide range of population densities, ideal for measuring DDHS.

During the study period (1988 – 2020), densities of wolves and cougars also varied in the study area. Wolves were reintroduced to YNP during 1995 – 1997 (Bangs & Fritts 1996), and their population increased to a maximum in 2003 and declined thereafter (Smith *et al*. 2020). Cougar densities generally increased across the study period (Ruth *et al*. 2019; Marcus *et al*. 2022). Consequently, the risk of predation by each of these predators has varied substantially across the timeseries of elk distribution analyzed here.

### Model structure

We modeled elk counts (*n_i,t_*) within a pixel (*i*) in a year (*t*) using a Bayesian generalized linear mixed model (GLMM). We used 1 km^2^ pixels to average over elk daily movements (Appendix S1). We did not have data to directly account for imperfect detection, but we found our results were insensitive to it (Appendix S2). We used a negative-binomial likelihood to accommodate overdispersion in our count data, a distribution commonly used to model animal group sizes (Ma *et al*. 2011). Factors other than habitat, e.g., sex-specific social interactions, are at least partially responsible for group size distributions (Gerard *et al*. 2002), which results in overdispersion that should be modeled to avoid overfitting. Elk group sizes in our dataset range from 1 to over 1,000 individuals, as seen with elk in other areas (Proffitt *et al*. 2012; Brennan *et al*. 2015).

We modeled the expected count (*λ_i,t_* = *E*[*n_i,t_*]) in each pixel and year as a function of 22 covariates (indexed by *k*; Table S1), a time-varying offset (*α_t_*), a temporal random effect (*η_i,t_*), and a spatial random effect (*s_i_*, eqn. 1), all of which we describe in detail in Appendix S3. We used the natural logarithm (hereafter, log) as the link function.

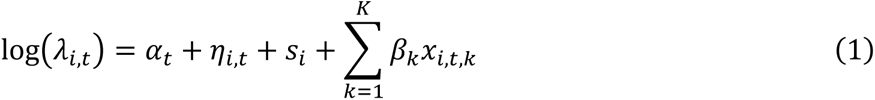

We estimated the size parameter of the negative binomial distribution as a single free parameter (*r*) such that: 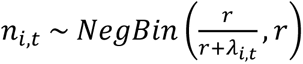.

Because we set the area of each pixel to 1.0 km^2^, we can interpret *λ_i,t_* as expected density in elk/km^2^. Note that “expected density” refers to the expected value of our response variable. Contrast this with “(log) average range-wide density,” which is a predictor variable, hereafter `log(Dens)`.

### Habitat variables

We treated habitat variables as fixed effects representing food, safety, or other conditions. The slope of each variable measures habitat selection (see Appendix S4). We rasterized all count data and variables on a 1-km grid, retaining only pixels falling within the study area (N = 1,978). We describe processing for all variables in Appendix S5. Variables describing conditions – snow-water equivalent (SWE), elevation, and cosine and sine of aspect (northing and easting respectively; Table S1) – were included to control for important known drivers of elk density but were not the target of inference.

Because northern Yellowstone elk are primarily grazers (Houston 1982), we measured food using total herbaceous biomass from the Rangeland Analysis Platform (RAP) annual biomass (v2.0) layer (Robinson *et al*. 2019; Jones *et al*. 2021). This layer combines information about total growth from 16-day NDVI data with plant functional type estimates to calculate growth of grasses and forbs (the annual layer sums all 16-day layers). For each winter, we used the layer corresponding to the preceding growing season to measure potential forage availability for elk that winter. We log-transformed this covariate to reduce the influence of very high values from agriculture outside YNP. We refer to this as the food variable.

We measured predation risk using the risky places approach such that habitat covariates indexed predation risk (Moll *et al*. 2017). Previous research has shown that risk to elk from wolves and cougars varies with tree canopy openness and terrain roughness. The wolf habitat domain is characterized by high openness and low roughness, whereas the cougar habitat domain is characterized by lower openness and high roughness (Kohl *et al*. 2019). On a fine temporal (5 h) and spatial scale (30 m), Kohl et al. (2019) showed that individual elk manage risk from wolves and cougars by moving into each predator’s habitat domain at the time when that predator is least active. At the coarser scale of our analysis, we expected elk to select 1-km^2^ pixels with a mixture of intermediate levels of openness and roughness that facilitate efficient switching between wolf and cougar habitat domains across the diel cycle. Thus, we expected elk density to be greatest at intermediate values of these variables, which indicate high heterogeneity in the pixel (Appendix S1; Fig. S4). To test this, we included linear and quadratic terms for openness and roughness to parameterize a parabola that quantified how elk select for safety at the 1-km^2^ scale. Hereafter, we refer to these as the safety variables. To check the raw data for a negative correlation between food and safety, we created a composite safety variable by taking the product of the openness and roughness rasters. We used Spearman’s correlation to check this assumption. All other food-safety comparisons were model based (see *Quantifying DDHS*).

To support our assertion that the safety variables reflect how elk perceive risk, we included interactions between wolf and cougar densities and each of the safety variables. If the safety variables were good metrics of safe and risky places, we expected predator densities to alter the strength of selection for the safety variable or shift the parabola vertex (most preferred openness/roughness). We predicted increasing wolf density would decrease selection strength or push the vertex away from open and smooth (i.e., away from the wolf and into the cougar habitat domain). Conversely, we predicted increasing cougar density would increase selection strength or push the vertex towards open and smooth habitat. Whereas aerial surveys occurred mostly during daylight hours before noon when wolves were more active than cougars (Kohl et al. 2019), we did not expect an effect of survey timing due to our coarse spatial scale (Appendix S1).

To measure DDHS, we included an interaction between the food/safety variables and `log(Dens)`. These interactions allowed for flexibility in the patterns of DDHS, but due to the linear and quadratic terms involved in the formulation, interpretation of the effects is most easily accomplished graphically (see *Quantifying DDHS*). Note that we assume that DDHS occurs as a function of the current density, not a time-lagged density. Whereas time lags are important for density dependence to operate on population growth rates (Turchin 1990), we assume that the mechanisms for DDHS (competition for resources, safety in numbers) operate more instantaneously than their fitness consequences.

### Model fitting

We created all quadratic and interaction terms and separately scaled and centered them before fitting. We did not transform the log-density offset or the sine and cosine of aspect, but we z-transformed all other variables to facilitate model fitting. We performed all data preparation and analyses in R (v. 4.1.1), and we fitted the GLMM via MCMC using R package NIMBLE v. 0.11.1 (de Valpine *et al*. 2017, 2021). We used Laplace priors with mean 0 on all regression parameters, referred to as the “Bayesian lasso,” to prevent overfitting (Hooten & Hobbs 2015). We ran the model for 100,000 iterations across three chains, discarded the first 20,000 as burn-in (including adaptation), and thinned by 20 to obtain 4,000 posterior samples/chain for inference. All analysis code and data are available on GitHub and published through Zenodo (doi:10.5281/zenodo.6687904).

### Model evaluation

We used the Gelman-Rubin statistic to evaluate MCMC convergence (Gelman & Rubin 1992). We then used out-of-sample data to validate our model. Counts outside of YNP in the Montana portion of the study area were unavailable in 1994, 2002, or 2004, so we withheld the YNP data for these years from model fitting and used them to validate our model. Additionally, we withheld data from 2020, which were available for the entire study area, as an additional year of validation data.

To perform validation, we used our fitted model to predict expected elk densities for each year, including all fixed and random effects. We compared expected elk densities under the model to observed densities from the count by calculating ordinary residuals (*n_i,t_* − *λ_i,t_*), Pearson residuals 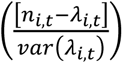, and Spearman’s correlation coefficient between predicted and observed densities. We compared ordinary residuals and Spearman’s correlation between the training and testing years. We used Pearson residuals to check for residual spatial autocorrelation by fitting non-parametric spline correlograms to these residuals (Bjørnstad & Falck 2001).

To estimate model goodness-of-fit, we calculated a likelihood-based pseudo-R^2^ for each posterior sample from the MCMC, yielding a distribution of pseudo-R^2^. We calculated our pseudo-R^2^ using the method of Nagelkerke (1991), which compares the likelihood of the data under the fitted model to a null model. For our null model, we fitted a model where expected elk density was a function of only the time-varying offset (*α_t_*), and the only other parameter estimated by the model was the size parameter of the negative binomial distribution, *r*.

### Quantifying DDHS

We quantified DDHS by measuring relative selection strength (RSS), the ratio of expected densities in two habitats (Avgar *et al*. 2017; Fieberg *et al*. 2021). RSS(*x*_1_, *x*_2_) is how many times more elk we expect at the habitat in the numerator (*x*_1_) compared to the habitat in the denominator (*x*_2_). The effect sizes of the habitat variable–density interactions indicate the strength of the DDHS, but because (1) this is on the scale of the link function, and (2) multiple variables are involved in the safety interaction, it is easier to visualize DDHS via RSS. Credible intervals around our RSS predictions account for the uncertainty in parameter estimates and their covariance, which clarifies inference on the quantity of interest (DDHS). Figure 1 provides a hypothetical example of how RSS indicates DDHS. Our model estimates expected density using linear (food; Fig. 1A) or quadratic (safety; Fig. 1B) formulations. A plot of RSS as a function of density reveals the pattern of DDHS: positive slopes show positive DDHS (Fig. 1C), negative slopes show negative DDHS (Fig. 1D), and a horizontal line shows no DDHS. We calculated RSS for the food and safety variables by comparing habitats that differ by one standard deviation (SD) in the focal variable (with all other variables held at their mean), which makes the magnitude of RSS comparable between variables. For safety, we altered openness and roughness by 0.5 SD each (1 SD total). The habitat we chose for the numerator (*x*_1_) always had the higher expected elk density. For the food variable, *x*_1_ = 661 kg/ha and *x*_2_ = 376 kg/ha (Fig. 1A). For the safety variables, we chose values that did not span the vertex (Fig. 1B). Under our hypothesis of a trade-off between food and safety, we predicted positive DDHS for food and negative DDHS for safety.

**Figure 1.**
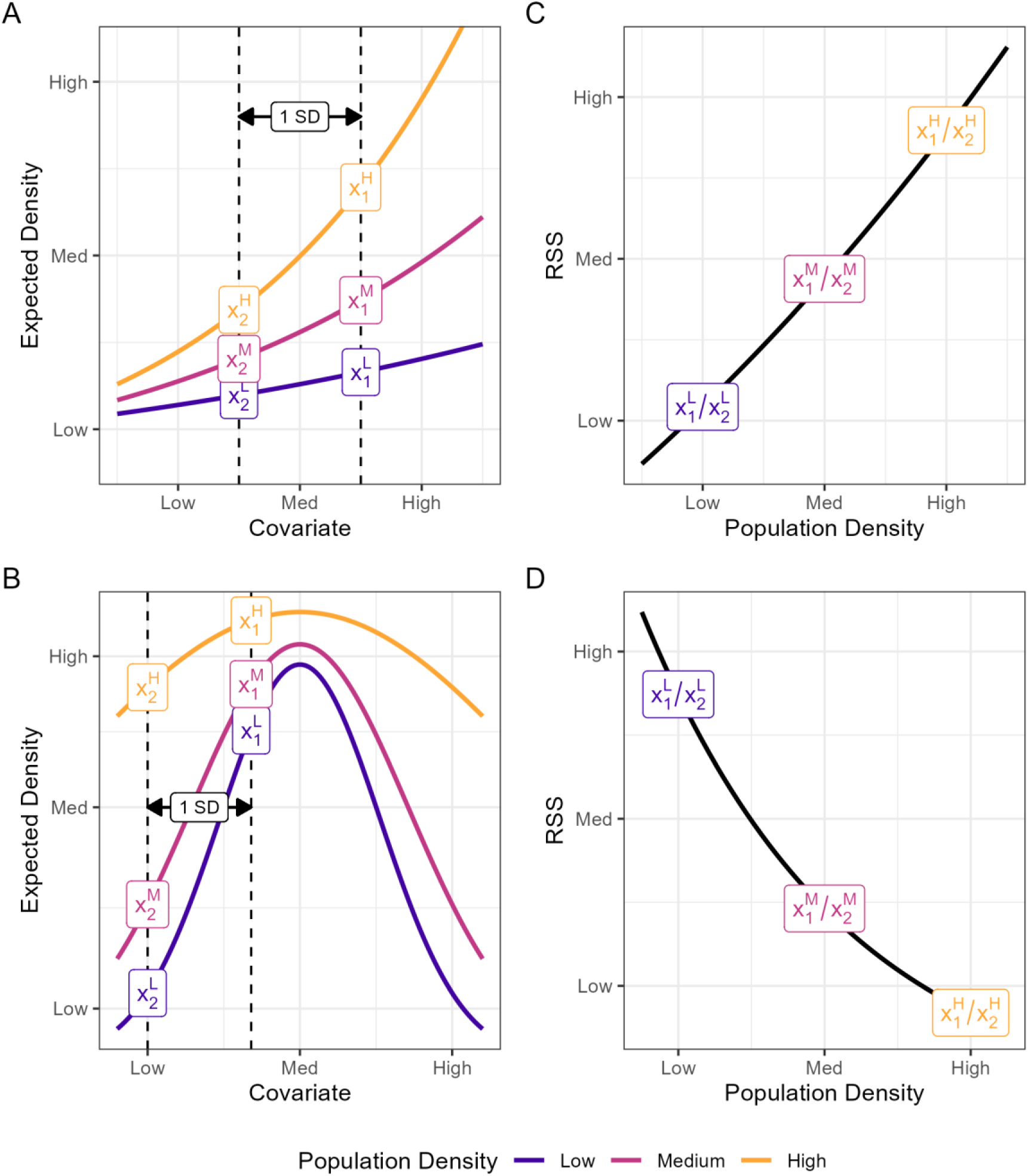
Conceptual figure depicting density-dependent habitat selection (DDHS). Left column (A and B) shows how expected density could change with a habitat covariate and average population density (color). Right column (C and D) recasts the patterns in A and B in terms of relative selection strength (RSS), the ratio of expected densities in different habitats. We calculated RSS(x1, x2) as the ratio of expected densities for a 1-SD change in the covariate (dashed vertical lines). For our purposes, the numerator (x1) is always larger than the denominator (x2). “H”, “M”, and “L” in superscript refer to high, medium, and low population density. Calculating RSS across a range of average population densities (e.g., x1H/x2H, x1M/x2M, x1L/x2L) yields the RSS curve, which more clearly demonstrates DDHS. In (A), expected density is modeled with just a linear term for the covariate, and expected density increases monotonically with an increase in the covariate (positive habitat selection). In this example, RSS (slope of each line) increases with population density; this is positive DDHS (C). Alternatively, if RSS decreased with population density, this would be negative DDHS (not shown). In (B), expected density is modeled with inear and quadratic terms such that expected density peaks at an intermediate value. A narrow parabola at low density indicates stronger selection, whereas a wider parabola at high density indicates weaker selection. This example demonstrates negative DDHS but note that positive DDHS is also possible. We calculated RSS as the ratio of expected densities when the covariate is near the vertex of the parabola (x1) vs. when the habitat covariate is lower (x2). Calculating RSS across a range of population densities yields the RSS curve (D), which in his case demonstrates negative DDHS. In summary, whether a habitat covariate is modeled with solely a linear term or also includes a quadratic term, the slope of the RSS curve plotted against population density shows the pattern of DDHS.

To understand the relative drivers of habitat selection, we compared the magnitude of RSS between our habitat variables. Together with our predictions of positive DDHS for food and negative DDHS for safety, we predicted that habitat selection was driven by safety at low densities and food at high densities; i.e., we predicted that the two RSS curves with opposite slopes would cross.

To understand the impact of density on predicted elk distribution, we created a map of the study area showing the change in RSS from high (9.3 elk/km^2^) to low (2.0 elk/km^2^) density which approximated the observed decrease over time. We fixed all spatial covariates and predator densities to their value in 2008, then we calculated RSS where *x*_1_ was each observed pixel and *x*_2_ was a habitat with mean values for all covariates. We took the log of RSS so that relative to mean conditions, positive values represent selection, zero represents no preference, and negative values represent avoidance. We repeated this at low and high elk density. For each pixel, we subtracted the high-density log-RSS from the low-density log-RSS, which we term Δ log-RSS. Positive values indicate that selection for the pixel increased as elk density decreased, whereas negative values indicate that selection decreased with elk density.

## Results

### Model overview & assessment of key assumptions

Our model indicated that elk selected for southwest-facing, low-elevation slopes with minimal snow cover. This is indicated by negative coefficients for `cos(Asp)` [aspect northing], `sin(Asp)` [aspect easting], `Elevation` and `SWE` (Fig. 2, Fig. S6).

**Figure 2.**
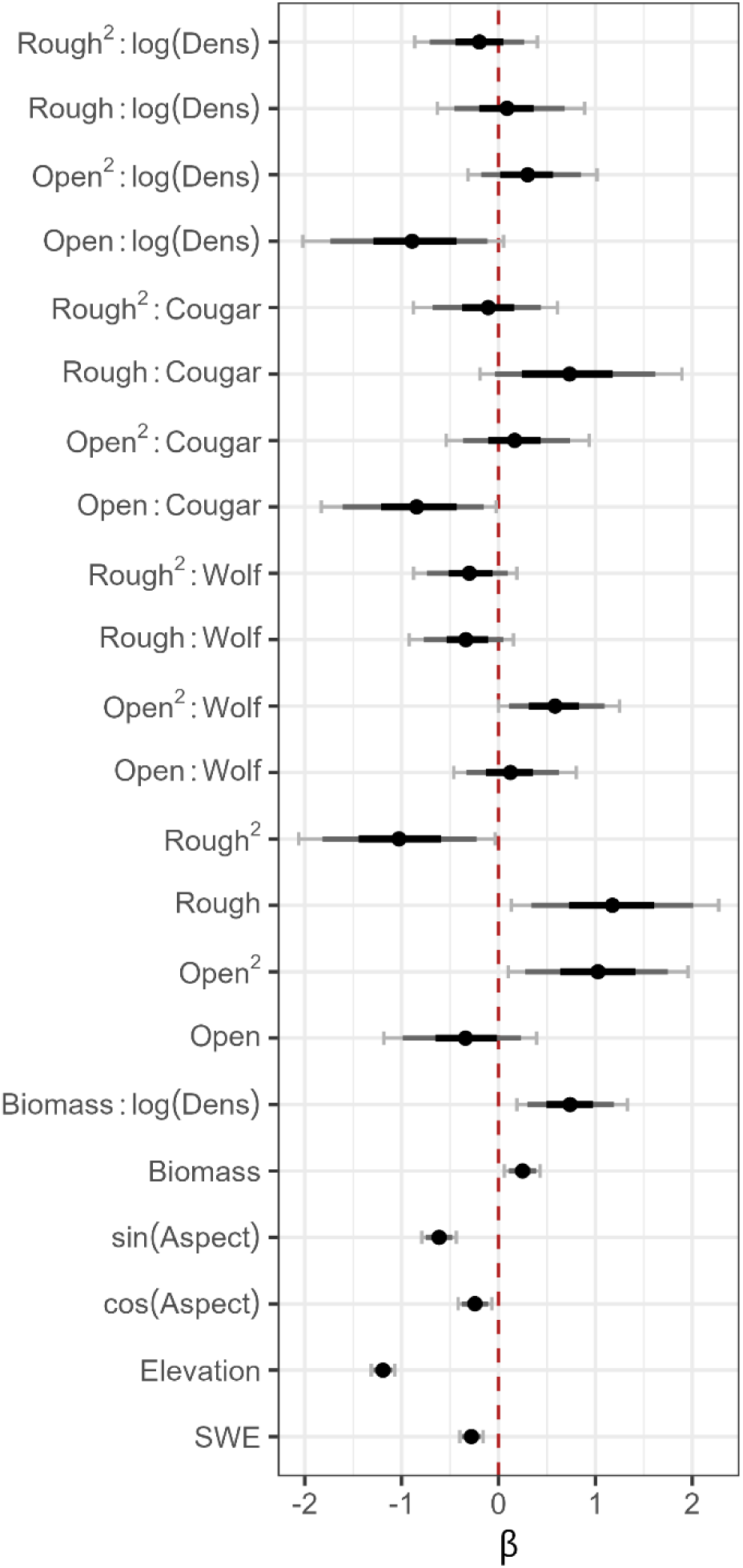
Fitted model coefficients. Points are posterior means and bars are credible intervals (black bars: 50% credible intervals; dark gray bars: 80% credible intervals; light gray bars with end caps: 90% credible intervals). Red dashed line indicates 0 (no effect).

Expected elk density increased with herbaceous biomass (Fig. S7A), consistent with our assertion that herbaceous biomass reflects elk forage availability. Contrary to our prediction that elk would most prefer an intermediate openness, expected elk density was greatest at 100% openness, and the relationship between openness and elk density was largely monotonic for the observed range of openness (Fig. S7B). Expected elk density was greatest for a roughness of 23.5 m (intermediate, as expected) with all other covariates held at their mean (Fig. S7C). Spearman’s correlation between biomass and the product of openness and roughness was -0.18, supporting our assumption that food and safety are negatively correlated.

Elk altered selection for safety variables with increasing predator densities in a manner that indicated openness and roughness were valid indices of spatial variation in predation risk. As wolf density increased, RSS for openness decreased (Fig. 3A), whereas it increased as cougar density increased (Fig. 3B), consistent with our predictions. Wolf density increased RSS for roughness, but the effect of cougars was negligible (Fig. S8). Wolf density shifted the vertex of the parabola in the expected direction (from wolf to cougar habitat domains): at low wolf density (0 wolves/100 km^2^), the vertex was 22.3 m (90% credible interval [CI]: 20.1 – 25.0), and at high wolf density (10 wolves/100 km^2^), the vertex was 26.1 m (90% CI: 22.1 – 32.1; Fig. 3C). Cougar density did not shift the vertex (Fig. 3D).

**Figure 3.**
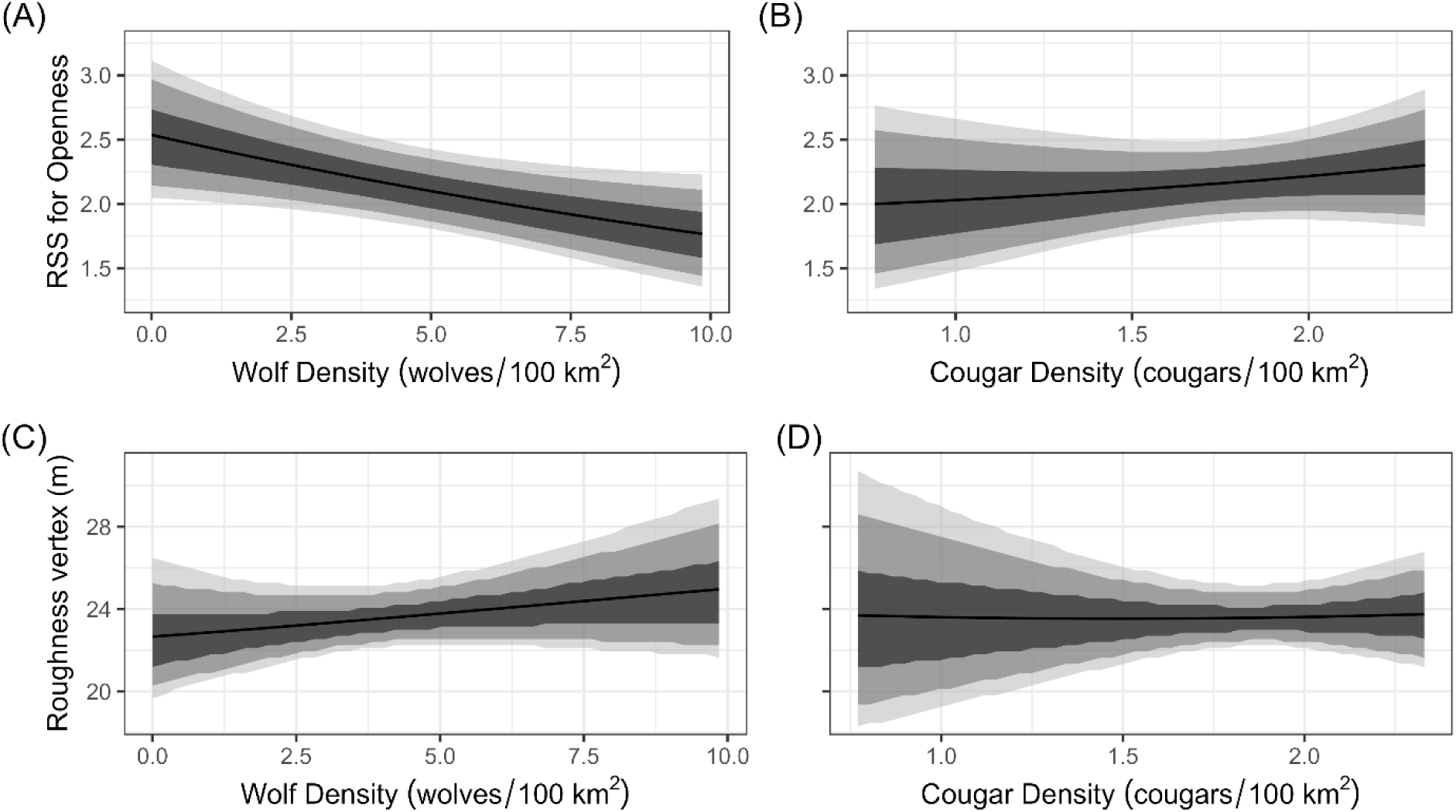
Changing elk selection for risk variables with predator density. Openness and roughness describe important dimensions of the wolf and cougar habitat domains, i.e., those habitats where they are most likely to kill elk. The wolf habitat domain is open and smooth, whereas the cougar habitat domain is rough and more forested. Relative selection strength (RSS) for openness (top row) is the ratio of expected density when openness is 100% vs. when openness is 82%, a 1 SD change in openness. (A) RSS for openness decreases with wolf density (less selection for the wolf habitat domain), whereas (B) it increases with cougar density (less selection for the cougar habitat domain). The roughness vertex (bottom row) is the value of roughness that is most strongly selected by elk, i.e., the vertex of the parabola describing selection. (C) The roughness vertex shifts to rougher habitat as wolf density increases (away from the wolf habitat domain); however, (D) elk did not shift the roughness vertex in response to cougar density. In all panels, predictions were made using samples from the entire posterior distribution. Solid black lines are mean effects and shaded gray envelopes are 50%, 80%, and 90% credible intervals.

### Density-dependent habitat selection

Elk exhibited positive DDHS with respect to food, demonstrated by the positive `Biomass:log(Dens)` coefficient (Fig. 2) and the positive slope of the RSS curve (Fig. 4). Elk exhibited negative DDHS for openness, shown by the negative `Open:log(Dens)` coefficient (Fig. 2) and the negative slope of the RSS curve (Fig. 4). Since the Open^2^:log(Dens) coefficient was estimated near 0 (Fig. 2), the linear coefficient drove the pattern (Fig. 4). These patterns are consistent with a trade-off between food and safety, expected under the multidimensional IFD (Avgar *et al*. 2020), supporting our main hypothesis. By contrast, elk exhibited no DDHS with respect to roughness. The 50% CIs for the `Rough:log(Dens)` and the Rough^2^:log(Dens) terms overlapped 0 (Fig. 2). Although the mean trend in RSS for roughness with increasing elk density was slightly positive, high uncertainty indicated this effect was negligible (Fig. S9).

**Figure 4.**
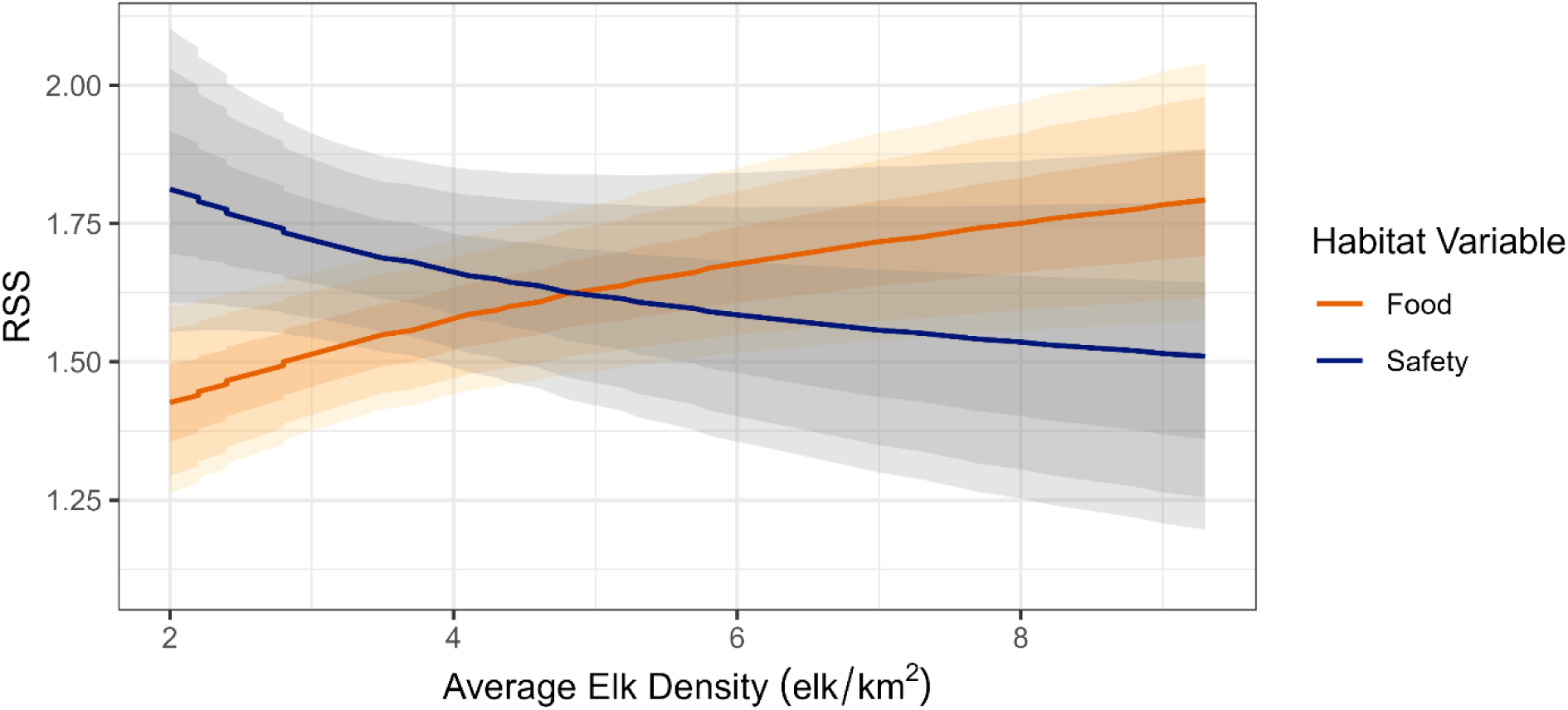
Density-dependent habitat selection for food and safety. At low density (x-axis), relative selection strength (RSS) for a one standard deviation (1 SD) change in safety (0.5 SD of openness and 0.5 SD of roughness) is greater than the RSS for a 1 SD change in food (herbaceous biomass). At high density, this relationship flips, with a greater RSS for food than for safety. Changing average range-wide elk density alters the main driver of elk habitat selection from safety at low density to food at high density. Solid lines are posterior mean estimates and shaded envelopes are 50%, 80%, and 90% credible intervals.

The stronger driver of elk habitat selection was safety at low elk density and food at high elk density (Fig. 4), matching our prediction. At low elk density (2.0 elk/km^2^), the RSS for a one-SD change in food was 1.43 (90% CI = 1.26 – 1.60) and the RSS for a one-SD change in safety was 1.81 (90% CI = 1.55 – 2.10). At high elk density (9.3 elk/km^2^), the RSS for a one-SD change in food was 1.79 (90% CI = 1.57 – 2.04) and the RSS for a one-SD change in safety was 1.51 (90% CI = 1.20 – 1.88).

Posterior mean Δ log-RSS varied from -1.6 to 3.0 with a mean of 0.0 across all pixels, indicating that changing selection resulted in habitat switching across space – the selection strength for some habitats decreased while the selection for other habitats increased, resulting in a redistribution of the population. Figure 5 illustrates the spatial pattern in Δ log-RSS.

**Figure 5.**
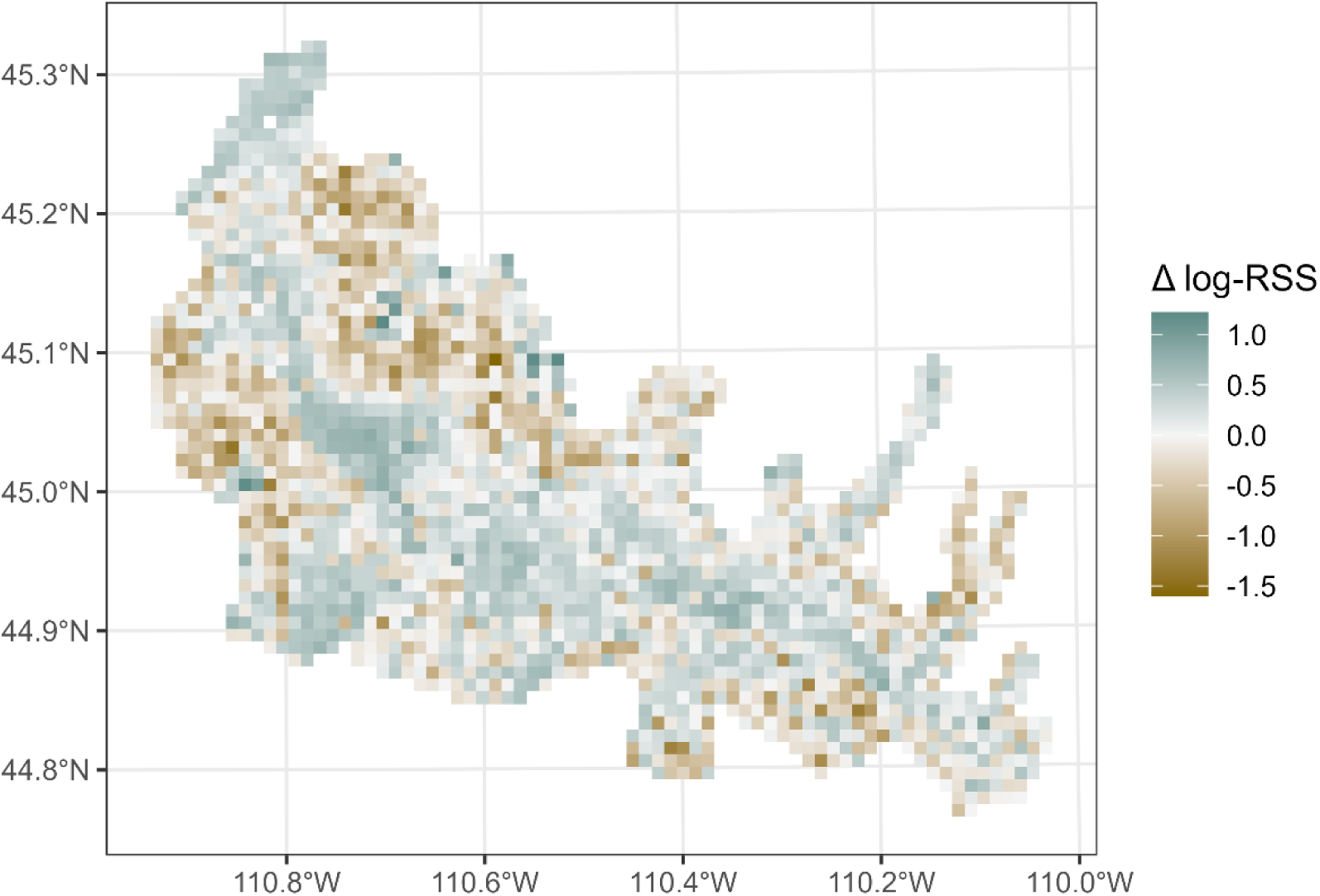
Change in log-RSS from high elk density to low elk density. Relative selection strength (RSS) is the ratio of expected density in each pixel of the landscape to the expected density in a habitat with all habitat variables held at their mean. The natural logarithm of RSS (log-RSS) is a measure of habitat selection, with positive values indicating preference versus the mean conditions, zero indicating no preference versus the mean conditions, and negative values indicating avoidance versus the mean conditions. Here, we plot Δ log-RSS, the difference between log-RSS when average range-wide elk density is high (9.3 elk/km2) versus when average range-wide elk density is low (2.0 elk/km2). Positive values (blue-green pixels) indicate that selection for the pixel increased as elk density decreased (the observed pattern over time), whereas negative values (brown pixels) indicate that selection for the pixel decreased as elk density decreased. All habitat variables and predator densities are held at their 2008 levels for demonstration.

### Model evaluation

The Gelman-Rubin statistics indicated MCMC convergence for all top-level parameters (Appendix S6). Ordinary residuals and Spearman’s correlations were similar between in-sample and out-of-sample predictions (Fig. S10). Residuals had mean near 0, indicating good model accuracy. Mean pseudo-R^2^ for our model was 0.065 (90% CI = 0.064 – 0.066), indicating low precision (to be expected under the negative binomial distribution). We found little to no residual spatial autocorrelation in all years (Fig. S11).

## Discussion

Density dependence is a fundamental concept in ecology, and its importance for a species’ abundance over time is well understood (Dennis & Taper 1994; Berryman & Turchin 2001; Brook & Bradshaw 2006). Although theory establishes that density-dependence should also act on abundance in space (Rosenzweig 1991; Morris 2003), comparatively little empirical work has demonstrated the role of density dependence in shaping population distribution. In this study, we provide rare empirical evidence that density alters drivers of habitat selection and population distribution in a free-living system, consistent with theoretical expectations under the multidimensional IFD (Avgar *et al*. 2020). This is a broadly important conceptual advance because it links population density to the community-level interactions (i.e., consumer-resource, predator-prey) that determine an organism’s distribution in space (Rosenzweig & Abramsky 1997). Observations of this system at a constant density would lead to erroneous inference on the drivers of habitat selection, biased predictions of population distribution, and misunderstanding of community interactions under unobserved densities.

Interactions between predator densities and safety variables indicate that the latter were valid indices of predation risk. RSS for openness decreased with wolf density (Fig. 3A) and increased with cougar density (Fig. 3B). The effect of predator densities on RSS for roughness was weak; rather, increasing wolf density pushed the most preferred roughness into rougher habitat (Fig. 3C). We found no effect of cougar densities on selection for roughness (Fig. 3D), and the predator density effects were greater for wolves than for cougars. Nevertheless, elk avoided densely forested, rough habitats across all elk and predator densities (Fig. S7 B and C), indicating that risk from cougars was an important driver of elk distribution. Our finding that elk selected for maximum openness (Fig. S7B) and that DDHS for safety was primarily driven by openness (Fig. S9B) further suggests that elk distribution was more strongly influenced by cougars than wolves. These results independently corroborate the findings of Kohl et al. (2019) that multiple predators (wolves and cougars) influence elk space use.

We found support for our hypothesis that the food-safety tradeoff in our system leads to a switch from safety driving distribution at low density to food driving distribution at high density (Fig. 4). We found evidence for positive DDHS for food and negative DDHS for safety, which we would expect when there is a tradeoff between food and predation risk. Indeed, biomass and the product of openness and roughness were negatively correlated (Spearman’s r = -0.18). While this food-safety tradeoff is expected to be ubiquitous in nature (Brown & Kotler 2004), it is difficult to measure, and our approach provides evidence of it. Were there no tradeoff, elk at low density could occupy habitats that provided both food and safety, and we would expect spillover from those habitats as density increases (i.e., negative DDHS). The negative correlation between food and safety means that elk cannot satisfy both of these requirements in the same place, thus giving rise to the tradeoff and habitat switching as density increases (i.e., positive DDHS; Greene & Stamps 2001; Avgar *et al*. 2020). The switch in the relative importance of food and safety leads to some habitats on the landscape becoming less selected while others become more selected; the result is a redistribution of the population (Fig. 5). This switch is consistent with theoretical and empirical work on behavioral ecology in predator-prey systems. Organisms should change their risk assessment based on conspecific density (Peacor 2003), and antipredator behaviors such as vigilance and habitat selection should adjust accordingly (Mooring *et al*. 2004). This is also consistent with the predation-sensitive food hypothesis, whereby both food and predation limit prey populations (Sinclair & Arcese 1995); as competition increases and food becomes limiting, prey will increase selection for food and decrease selection for safety. Our work demonstrates how these behaviors manifest in space.

Current empirical understanding of DDHS in free-living vertebrate systems is mostly limited to small mammals (e.g., Rosenzweig & Abramsky 1985; Morris 1989; Morris *et al*. 2000) or domesticated mammals on islands (e.g., Mobæk *et al*. 2009; van Beest *et al*. 2014b). Previous studies of DDHS in free-living systems, though rare, have also focused on elk (van Beest *et al*. 2014a; Merrill *et al*. 2020) or congeneric red deer (*Cervus elaphus*; McLoughlin *et al*. 2006; Pérez-Barbería *et al*. 2013); see Supplemental Discussion for a comparison with our work (Appendix S7). Often, space-use data are collected on short time scales during which population densities are similar (McLoughlin *et al*. 2010), and the resulting habitat selection models are snapshots of true, DDHS patterns (Boyce & McDonald 1999; Boyce *et al*. 2016; Northrup *et al*. 2022). Our results show that inference or predictions from these snapshots may be unreliable under changing densities.

Space use of organisms can be measured from two perspectives: a population-level (Eulerian) perspective which measures changes in population density at various places over time or an individual-level (Lagrangian) perspective which tracks the locations of individuals over time (Turchin 1998). Analyses from the Eulerian perspective lend themselves more readily to projection across space at large scales, whereas analyses from the Lagrangian perspective can uncover more detailed mechanism. Individual behavior scales up to population-level distributions (Mueller & Fagan 2008), so results from the two perspectives can, in principle, be reconciled. In practice, few studies have compared the Eulerian and Lagrangian approaches to studying population-level space use (but see Phillips *et al*. 2019; Bassing *et al*. 2022). In our case, we find both similarities and discrepancies between our work (Eulerian) and previous studies (Lagrangian) of elk habitat selection in northern Yellowstone (Appendix S7). Without the Lagrangian perspective, we could not identify whether DDHS occurred because (1) individuals changed their habitat selection traits (behavioral plasticity), or (2) individuals that exhibited certain habitat selection traits had differential survival and/or reproduction (demographic sorting), although previous research has suggested it is a demographic sorting effect (White *et al*. 2012). Which perspective provides the best understanding of DDHS is still an open question. Most empirical work on DDHS has made use of isodar theory (Morris 1988, 2011), which is constructed from the Eulerian perspective and naturally integrates fitness. An important knowledge gap concerns how to connect DDHS to fitness from the Lagrangian perspective, which can reveal more detailed mechanisms of how population regulation plays out in space.

We believe the capacity of DDHS to qualitatively alter community-level interactions is underappreciated and understudied (see Rosenzweig & Abramsky 1997), especially in predator-prey systems where positive DDHS may lead to landscape-scale habitat switching. For example, the relative importance of consumptive vs. non-consumptive effects (Peacor *et al*. 2013; Sheriff *et al*. 2020) may critically depend on DDHS (habitat selection is an important antipredator trait; Trussell *et al*. 2006). If prey only respond to risk by altering their habitat-selection traits at low density (when competition is less costly), the importance of non-consumptive effects may be overstated. Similarly, DDHS may impact competition between prey species. Habitat selection is an important mechanism for reducing competition between species, which may determine the level of competition (Rosenzweig 1981), especially when prey species face differential risk. For example, spatial overlap between bison and elk in northern Yellowstone is potentially greater than expected at high elk density if neither species responds strongly to predators.

In conclusion, consistent with theoretical expectations, density dependence alters habitat selection and distribution in free-living systems. Incorporating DDHS into models of distribution is crucial whenever density is variable; ignoring its effects may lead to severely compromised inferential and predictive performance. Our findings underscore that the effects of the food-safety tradeoff on prey distribution are dynamic, and that inference and prediction in these systems depends on prey density.

## Acknowledgements

Our work was partially supported by NSF grant NSF-DEB 1245373. Financial support for BJS was provided by the USU Office of Research. We thank the Northern Yellowstone Cooperative Wildlife Working Group for collecting the count data, with special thanks to C. Travis Wyman for help understanding the counts. Clark Rushing, Kezia Manlove, and Erica Stuber provided valuable comments on earlier versions of the analysis. Simona Picardi, Danielle Berger, Courtney Check, and Ronan Hart provided helpful comments on earlier drafts of the manuscript.

## Supporting Information

### Appendix S1 Aerial Survey Timing and Scale

We analyzed aerial survey data – designed to census the population – collected from 1988 to 2020. Elk were counted by 3 – 4 fixed-wing aircraft flying non-overlapping areas of the winter range (Fig. S1). Surveys took place over one to four days between late December and March of each year (Fig. S2A). Winter surveys were assigned to the year of that January; e.g., the survey for the winter of 1991 occurred in January 1991 and the survey for 1992 occurred in December 1991. The count did not occur in 1996 or 1997. The count was done via helicopter in 2019 and was not included due to differing protocol. Counts for 1989, 1991, 2006, and 2014 are considered unreliable as a census of the population and so were not included. For three years (1994, 2002, and 2004), georeferenced group data are only available inside of Yellowstone National Park (data from the State of Montana were not georeferenced). We excluded these data from model fitting and used them for model validation (see main text). A full census was available for 2020, but we chose to withhold it as an additional validation year with complete georeferencing.

Since survey timing varied from year to year, we checked that survey timing did not introduce a bias in our model. We plotted the model’s ordinary residuals as a function of survey date, and visually checked for any pattern by fitting a very flexible GAM smoothing line (Fig. S3). The fitted line showed no obvious pattern. We tested this further with an ANOVA, where we treated each of the 15 unique mean survey dates as categorical variables and the Pearson residuals as the response. We computed p-values for pairwise differences using Tukey’s Honest Significant Difference method to check for differences between Pearson residuals for any pair of dates. All p-values were large (>0.23; Table S2), which indicated no meaningful differences between residuals for any pair of dates.

Surveys recorded groups between 07:00 and 15:00, but 92% of observations were between 08:00 and 12:00 (Fig. S2B). Kohl *et al*. (2019) found that wolf movements peaked at 10:00 (in the middle of these survey times), whereas cougar activity peaked at 03:00 and 04:00 for male and female cougars, respectively (outside all survey times). Thus, surveys occurred at a time of day when wolves were the more active predator. To account for this and other potential temporal effects, we chose a coarse spatial scale to average over daily movements. Our 1 km x 1 km raster cells were chosen to be coarse enough to accomplish this. We verified that elk daily winter movements rarely exceeded this area by comparing the area of a raster cell (1 km^2^) to the area used daily by GPS-collared elk during winter (data not included in this manuscript). We found that on 93% of winter elk-days, a 100% MCP around all of an elk’s locations had an area < 1 km^2^ (B. Smith, pers. comm.). This was based on 44,444 elk-days from 169 unique individuals tracked between 2001 – 2022.

Furthermore, this coarse spatial grain is large enough to capture (potentially multiple) patches of trees within a matrix of openness. For example, a raster cell with 50% openness typically contains a mix of open and forested patches – not a homogenous patch of intermediate forest cover. This patchy habitat allows elk to shuttle between patches across the diel cycle, whereas a raster cell with 0% or 100% openness hinders this diel shuttling behavior (Fig. S4). Fitting a parabola to openness therefore measures how much heterogeneity in openness is selected for by elk.

We have designed our modeling approach with respect to spatial and temporal scale, while considering variation in survey dates and times, and, thus, we believe our model is a valid representation of the winter distribution of northern Yellowstone elk that is insensitive to survey date and time of day.

### Appendix S2 Sensitivity to Imperfect Detection

Previous research evaluating aerial counts of elk, including work in northern Yellowstone, has found that elk group size and vegetation cover (i.e., tree canopy openness) affect detection of elk groups (Samuel *et al*. 1987; Singer & Garton 1994). Because openness is an important covariate in our habitat selection model, we were concerned that imperfect detection as a function of openness could bias our results. We did not have data to explicitly estimate imperfect detection for each of the survey years we analyzed (e.g., mark-resight data, double observers, or multiple surveys). Instead, to evaluate any potential bias, we conducted a sensitivity analysis wherein we excluded pixels outside a certain openness threshold from the analysis and refitted the model. We compared coefficients and predicted DDHS patterns between the original model and the reduced dataset models.

We used three openness cutoffs: <30%, <50%, and >99.9%. We discarded all pixels that had openness less than/greater than the cutoff in *any* year and refitted the model. The first two cutoffs (<30% and <50%) discarded the pixels we considered least reliable, i.e., due to their low openness, we expected detectability to be low. The third cutoff (>99.9%) discarded the pixels we considered most reliable; we did this to determine if the remaining pixels were reliable enough to replicate the pattern on their own. We ran each MCMC procedure with 2 chains and 50,000 iterations per chain, discarded the first 22,000 iterations as burn-in and adaptation, and then thinned by 7 to 4,000 iterations per chain. All code for this analysis can be found with the main analysis on GitHub and archived through Zenodo (doi:10.5281/zenodo.6687905).

The resulting dataset for the <30% cutoff was 95% of the original data, the resulting dataset for the 50% cutoff was 70% of the original data, and the resulting dataset for the >99.9% cutoff was 73% of the original data. We found all of our resulting patterns were qualitatively similar between the models: we found positive DDHS for biomass, negative DDHS for safety, and a change in their relative importance (Fig. S5). Note that for the <99.9% cutoff, the means do not cross in the plotted domain, but the credible intervals overlap, and the pattern remains similar (Fig. S5D). Thus, we concluded imperfect detection did not meaningfully impact our inference.

### Appendix S3 Model Structure and Random Effects

As detailed in the main text, we modeled the expected count (*λ_i,t_*) in each pixel in each year as a function of 22 covariates (indexed by *k*; Table S1), a time-varying offset (*α_t_*), a temporal random effect (*η_i,t_*), and a spatial random effect (*s_i_*, eqn. 1). We described the fixed covariates in the main text, and we described the offset and random effects in detail here.

#### Offset

To account for changing total abundance across years, we included a time-varying offset, *α_t_*.The offset was the log of average range-wide density of elk across the study area in each year, i.e., the natural logarithm of the total count (*N_t_*) divided by the number of pixels in the study area (Ω), 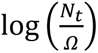. If none of the covariates had any effect (i.e., for all *k*, *β_k_* = 0), then the expected density in each pixel would simplify to 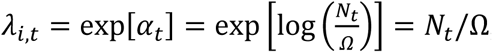, which is simply the average range-wide density for that year. In that way, the offset acts like a time-varying intercept. Note that when we refer to “average range-wide density” or “log average range-wide density” throughout, we are referring to a predictor variable, such as this offset described here or the interaction terms described below. Contrast this with “expected density,” which is the expectation of our response variable, *λ_i,t_*.

#### Temporal Random Effect

We used the temporal random effect (*η_i,t_*) to account for temporal fluctuations in the effect of a raster pixel being outside YNP compared to inside YNP. The two main motivations for this were (1) an observed shift in the elk distribution from primarily inside YNP in the early years of our study to primarily outside the northern boundary of YNP in later years (Tallian et a. 2017; MacNulty et al. 2020), and (2) a late hunt managed by the state of Montana that occurred in January of each year prior to 2010 but ceased afterward (MacNulty et al. 2020). We estimated the effect in each year (*t*) as coming from a normal distribution with mean *μ_η_* and variance 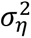. We multiplied the effect by a binary covariate indicating whether each pixel was outside of YNP, with outside = 1 and inside = 0.

The temporal random effect (*η_i,t_*) had a mean of -1.74 (90% credible interval = -2.05 – -1.42), and a variance of 0.56 (90% credible interval = 0.24 – 1.09). The posterior mean effect for each year ranged - 2.77 – -0.39, and the trend was positive over time (Fig. S12). In this context, negative values indicate lower elk densities outside YNP than would be expected based on the fixed effects in the model and the positive trend indicates increasing densities outside YNP over time.

#### Spatial Random Effect

We used the spatial random effect (*s_i_* ∈ ***s***) to account for residual spatial autocorrelation in elk density, i.e., consistent spatial pattern in elk density not captured by the other terms in the model. We modeled it as a Gaussian process (a.k.a., kriging), i.e., the vector ***s*** is a realization of a multivariate normal distribution with mean 0 and variance-covariance matrix, **Σ** (***s*** ∼ *N*(**0, Σ**)). Each element of the variance-covariance matrix was calculated as a decaying exponential function of the distance between the two pixels, ‖*x_i_* − *x_j_* ‖, and two free parameters, *σ* and *ρ*, estimated by the model: 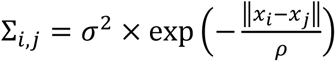. The parameter *σ* controlled the variance within a pixel (i.e., the distance between a pixel and itself is 0, thus Σ_*i,i*_ = *σ*^2^). The parameter *ρ* controlled the rate of decay with distance, with smaller *ρ* indicating faster decay. To improve model fitting, we scaled distances between pixels to range from 0 – 1 by dividing by the maximum distance.

The estimated spatial random effect, which was constant across time, was near 0 inside YNP and positive outside YNP (Fig. S13). The posterior mean of each *s_i_* ranged from -0.03 to 1.61, but the mean of each mean *s_i_* inside YNP was 0.01, whereas outside YNP it was 0.22. The decay parameter of the spatial covariance function (*ρ*) had posterior mean = 14.08 (90% credible interval = 6.67 – 19.49) and the standard deviation parameter (*σ*) had posterior mean = 0.63 (90% credible interval = 0.43 – 0.79). Thus, the covariance between two pixels separated by 1 km was 0.41 (90% credible interval = 0.19 – 0.63) and the covariance between two pixels separated by 100 km was 0.38 (90% credible interval = 0.16 – 0.59), indicating overall low spatial autocorrelation (Fig. S14). Together, the two random effects resulted in some spatially autocorrelated variation in elk densities outside YNP, with densities inside YNP almost entirely captured by the fixed effects (Fig. S15).

### Appendix S4 Equivalence to Habitat Selection Functions

Our approach to modeling habitat selection at the population level was parallel to more familiar approaches taken with individual telemetry locations, habitat-selection functions (HSFs, a.k.a., RSFs). In most HSFs, the observed locations (coded as 1) and background locations (coded as 0) are analyzed with logistic regression, the parameters of which are exactly equivalent to the parameters of an inhomogeneous Poisson point process (IPP) model (reviewed in Fieberg *et al*. 2021). The IPP model describes the *density* of points as a function of habitat variables; however, in telemetry-based HSFs, the resulting predictions are only *proportional* to density (Avgar *et al*. 2017). An equivalent model can be fit by discretizing space and using Poisson regression (Manly *et al*. 2002; Hooten *et al*. 2017). As we describe below, we fit our HSF using the latter approach – we modeled elk counts as a function of habitat covariates in discrete raster pixels (note that we used the negative binomial distribution as an overdispersed Poisson distribution). The resulting coefficients (the βs described in the main text for the covariates listed in Table S1) are interpreted in exactly the same way as if they had come from a telemetry HSF – but in our population-level study, we interpret the model predictions as spatiotemporally explicit elk densities.

### Appendix S5 Covariate Processing

We began with a master template raster, with a projected coordinate reference system (EPSG: 26912; UTM zone 12, NAD83) – the extent determined by the northern Yellowstone elk winter range polygon (Fig. S1) – and 1 km x 1 km resolution. We rasterized all count data and covariates (see below) on this template, using bilinear interpolation when resampling the covariate rasters. We projected rasters from their native resolution to our master template raster using the function `projectRaster()` from the `raster` package (Hijmans 2022). After all covariate processing, we retained only pixels falling within the northern winter range boundary for analysis, a total of 1,978 1-km^2^ pixels.

#### Conditions

We included conditions to account for known drivers of elk density that were not the target of inference. These were snow-water equivalent (SWE), elevation, cosine of aspect (aspect northing), and sine of aspect (aspect easting).

We obtained SWE data from Daymet v 4.0 (Thornton *et al*. 2020).

We obtained a digital elevation model (DEM) from the National Map’s 3DEP program (U.S. Geological Survey 2020) The DEM was projected to a 30 x 30 m resolution in a UTM projection (zone 12N, NAD83). We extracted elevation from the DEM directly and processed the DEM to calculate aspect (and roughness, see *Safety*) rasters in R using the `terrain()` function from the `raster` package (Hijmans 2022). We calculated aspect northing (cosine of aspect) and aspect easting (sine of aspect) from the aspect raster. Following all calculations at a 30-m resolution, we reprojected the raster to our master template raster.

#### Food

We measured food using total herbaceous biomass from the annual biomass (v2.0) layer from the Rangeland Analysis Platform (RAP; Jones *et al*. 2021). For each winter, we used the layer corresponding to the previous growing season to measure potential elk forage available from the previous growing season (e.g., for winter 1988, we used total annual biomass from 1987). We downloaded total biomass layers from RAP for 1987 – 2019, which were natively in geographic coordinates (longitude and latitude; WGS84) at approximately 30 m resolution (0.0002694946 degrees) and organized in two layers: annual herbaceous biomass and perennial herbaceous biomass. We summed these two layers to obtain total herbaceous biomass. We reprojected a stack of total herbaceous biomass rasters for each year to our master template. Finally, we log-transformed this covariate to reduce the influence of very high values from some areas of irrigated agriculture north of YNP.

#### Safety

We measured predation risk using the risky places approach, and previous research has shown that risk to elk from both wolves and cougars can be described by using tree canopy openness and terrain roughness (see main text for details).

We measured openness using the cover product from RAP. For each winter, we used the layer corresponding to that year (e.g., for winter 1988, we used cover for 1988). We downloaded the cover rasters from RAP for 1988 – 2020, and we extracted the trees layer, which measures the tree coverage in a pixel in %. As with herbaceous biomass, the cover layers were natively in geographic coordinates (longitude and latitude; WGS84) at approximately 30 m resolution (0.0002694946 degrees). We converted from tree cover to openness by subtracting the values from 100%, and then we divided by 100 to leave the data in the range [0, 1], where 1 is completely open (no tree cover) and 0 is 100% tree cover. We reprojected a stack of openness rasters for each year to our master template.

We measured roughness using the DEM described above (see *Conditions*). We used the `terrain()` function with `opt = “roughness”` to calculate roughness from the DEM. For a focal pixel, roughness was measured as the difference between the maximum and the minimum value of the pixel and its 8 surrounding pixels, thus the units were m (inherited from the DEM). After calculating roughness at the 30-m resolution, we reprojected the raster to our master template raster.

#### Predator densities

To support our assertion that the safety variables reflect how elk perceive risk, we included interactions between wolf and cougar densities and each of the safety variables. Wolf counts were available from National Park Service monitoring, and we used wolf population size for the northern winter range, specifically. We converted wolf counts to densities by dividing by the area of the northern winter range polygon that falls within the boundaries of Yellowstone National Park (wolves/km^2^; Fig. S1) and then multiplied by 100 to get wolves/100 km^2^.

Cougar densities in northern Yellowstone have been estimated from field data during 3 phases across the period of time we analyzed. Phase 1 estimated the density each year from 1987 – 1993, Phase 2 from 1998 – 2004, and Phase 3 from 2014 – 2017. Densities increased from Phase 1 through Phase 2, leveling off by Phase 3. We used linear interpolation across the entire dataset to fill in the gaps between Phase 1 – 2 and between Phase 2 – 3. From the end of Phase 3 until the end of our elk timeseries (2018 – 2020), we used the average cougar density in Phase 3, rather than assuming continued growth. Fig. S16 shows estimated and interpolated cougar densities.

### Appendix S6 Model Evaluation

We evaluated convergence of our MCMC algorithm by using the Gelman-Rubin statistic (Gelman & Rubin 1992), which we calculated using the function `gelman.diag()` from the R package `coda` (Plummer *et al*. 2006). Potential scale reduction factors (PSRFs) near 1 indicate MCMC convergence. PSRF point estimates for all top-level parameters were < 1.02 and the upper bounds of all confidence limits were < 1.06 (Table S3), indicating good convergence.

To estimate model goodness-of-fit, we calculated a likelihood-based pseudo-R^2^ for each posterior sample from the MCMC, yielding a distribution of pseudo-R^2^. We calculated pseudo-R^2^ using the method of Nagelkerke (1991), which compares the likelihood of the data under the fitted model to a null model. For our null model, we fitted a model where expected elk density was a function of only the time-varying offset (*α_t_*), and the only other parameter estimated by the model was the size parameter of the negative binomial distribution, *r*.

Ordinary residuals and Spearman’s correlations were similar between in-sample and out-of-sample predictions (Fig. S10). Residuals had mean near 0, indicating good model accuracy (Fig. S10A), and Spearman’s correlations were similar between in-sample data (mean r = 0.28) and out-of-sample data (mean r = 0.27; Fig. S10B). Mean pseudo-R^2^ for our model was 0.065 (90% CI = 0.064 – 0.066), indicating relatively low precision. We found little to no residual spatial autocorrelation in all years (Fig. S11).

Our model’s low pseudo-R^2^ is not surprising. Factors other than habitat, e.g., social interactions among elk, are at least partially responsible for group size distributions (Gerard *et al*. 2002; Brennan *et al*. 2015). Elk group sizes in our dataset range from 1 to over 1,000 individuals, and this is seen with elk in other study areas (Proffitt *et al*. 2012; Brennan *et al*. 2015). Thus, to capture the large variance in group size distributions, we chose to model elk counts using the negative binomial rather than the Poisson distribution. The free dispersion parameter of the negative binomial can soak up the variation in counts rather than attributing that variation to covariates. This is an important feature of our model: modeling elk counts with the Poisson distribution would be misleading as it would require a stronger effect of habitat to capture the variation in the observed data (i.e., placing more weight on the covariates and potentially overfitting). One result of this is that predicting observed number of elk in a particular pixel shows low precision, i.e., a wide range of counts is plausible for a given expected value. While our model has low precision, it does have good accuracy, shown by out-of-sample validation (Fig. S10). Thus, our model estimates the expected number of elk in a pixel accurately and captures the slopes of our covariates from which we draw inference. Translating this expectation to an actual count was not the goal of our model.

### Appendix S7 Supplemental Discussion

#### Comparison to previous studies

##### Elk and Red Deer

Previous studies of DDHS in free-living systems, though rare, have also focused on elk (van Beest *et al*. 2014; Merrill *et al*. 2020) or congeneric red deer (*Cervus elaphus*; McLoughlin *et al*. 2006; Pérez-Barbería *et al*. 2013). In the absence of predation risk, red deer exhibited negative DDHS and spillover, as expected (McLoughlin *et al*. 2006; Pérez-Barbería *et al*. 2013).

Most similar to our work, Merrill *et al*. (2020) looked at DDHS in elk in Ya Ha Tinda under risk of wolf predation. This system exhibits the negative correlation between food and safety that we would expect would generate results similar to ours; however, the results do not show this. Ya Ha Tinda elk exhibited negative DDHS for forage biomass and positive DDHS for risky places – exactly the opposite of what we found. Additionally, an interaction between forage biomass and wolf risk became more strongly negative with elk density – i.e., elk selected the best forage even less under wolf predation risk, and this response to predators became stronger when elk were at high density (see Fig. 2 in Merrill *et al*. 2020). One possible explanation is that elk can make use of the “human-shield” effect to effectively reduce their risk from wolves in this system. If this effect sufficiently reduces the predation risk, then we would no longer expect positive but instead negative DDHS for food, since elk can use the “shield” to select for the best forage, even at low density. In that case, the correlation between the food and safety variables would result in both being subject to negative DDHS, just as Merrill et al. (2020) reported. Nevertheless, this is another piece of evidence that DDHS occurs in large herbivores.

##### Northern Yellowstone

Several previous studies have focused on individual winter habitat selection by telemetered northern Yellowstone elk. Ours is the first to take a population-level (Eulerian) perspective; nonetheless, there is some broad agreement in our conclusions. Mao et al. (2005) compared habitat selection in two distinct time periods: 1985–1990 (pre-wolf reintroduction) and 2000–2002 (post-wolf reintroduction). While they assumed no DDHS, the population declined >25% (9.3 elk/km^2^ in 1988; 6.7 elk/km^2^ in 2001) between these periods. They found elk selected more open habitats in winter post-wolf reintroduction – at lower density – agreeing with our finding of negative DDHS for openness (Fig. S9B). Kohl et al. (2018, 2019) used step-selection functions (SSFs) to show that elk selected habitat to avoid wolves (2018) or wolves and cougars (2019), but only when those predators were most active. Elk use of risky places at times of day when risk was low may suggest that during the timing of our aerial surveys, elk would indeed prioritize food over safety. These studies used data from 2001 – 2004, during which time elk densities were at moderate levels but decreasing; they did not account for possible DDHS. Elk density in 2001 was 6.7 elk/km^2^ and dropped to 4.3 elk/km^2^ by 2004, densities at which our model predicts food and safety are approximately equally important (Fig. 4).

Other studies of northern Yellowstone elk have qualitative discrepancies with our results. Fortin et al. (2005) concluded that elk used resources less under risk from wolves. Their study occurred at an intermediate density (6.7 elk/km^2^ in 2001), when our model suggests food should be a stronger driver than safety (Fig. 4). This discrepancy could be due to scale and study design; they focused at a fine spatiotemporal scale using SSF. Cusack et al. (2020) looked at spatial overlap with risk variables. They analyzed elk data from 2012 – 2015, when elk densities were consistently near 2.1 elk/km^2^ – the lowest in our timeseries. Our model suggests that safety should be most important during this time, yet Cusack et al. (2020) found very little evidence that elk avoid wolves in time or space. This discrepancy is possibly explained by methodological differences, with their analysis mostly relying on either winter-long spatial overlap or very fine scale encounters by individuals and our analysis relying on an annual snapshot of the population distribution.

### Appendix S8 Supplemental Tables

**Table S1.**
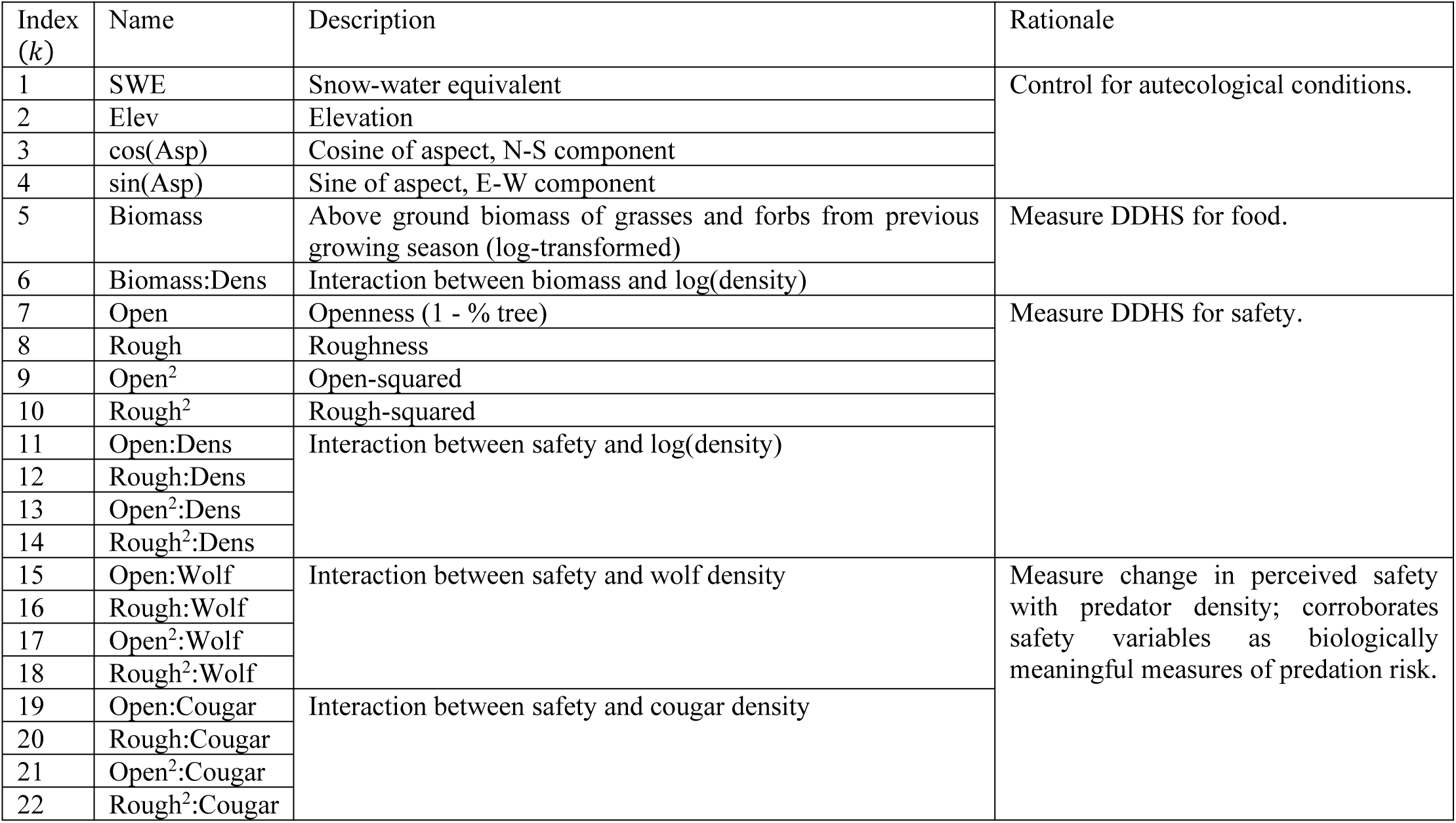
Model covariates. Covariates included to model expected density (*λ_i,t_*). Colons (:) in the “Name” column indicate interactions. “Rationale” describes the reason we included a parameter in the model. The first four variables (k = 1 – 4) measure fixed conditions that are known to affect elk density but were not the main target of inference. Variables including biomass (k = 5 – 6) are the food variables and variables including openness or roughness (k = 7 – 22) are the safety variables. DDHS = density-dependent habitat selection.

**Table S2.**
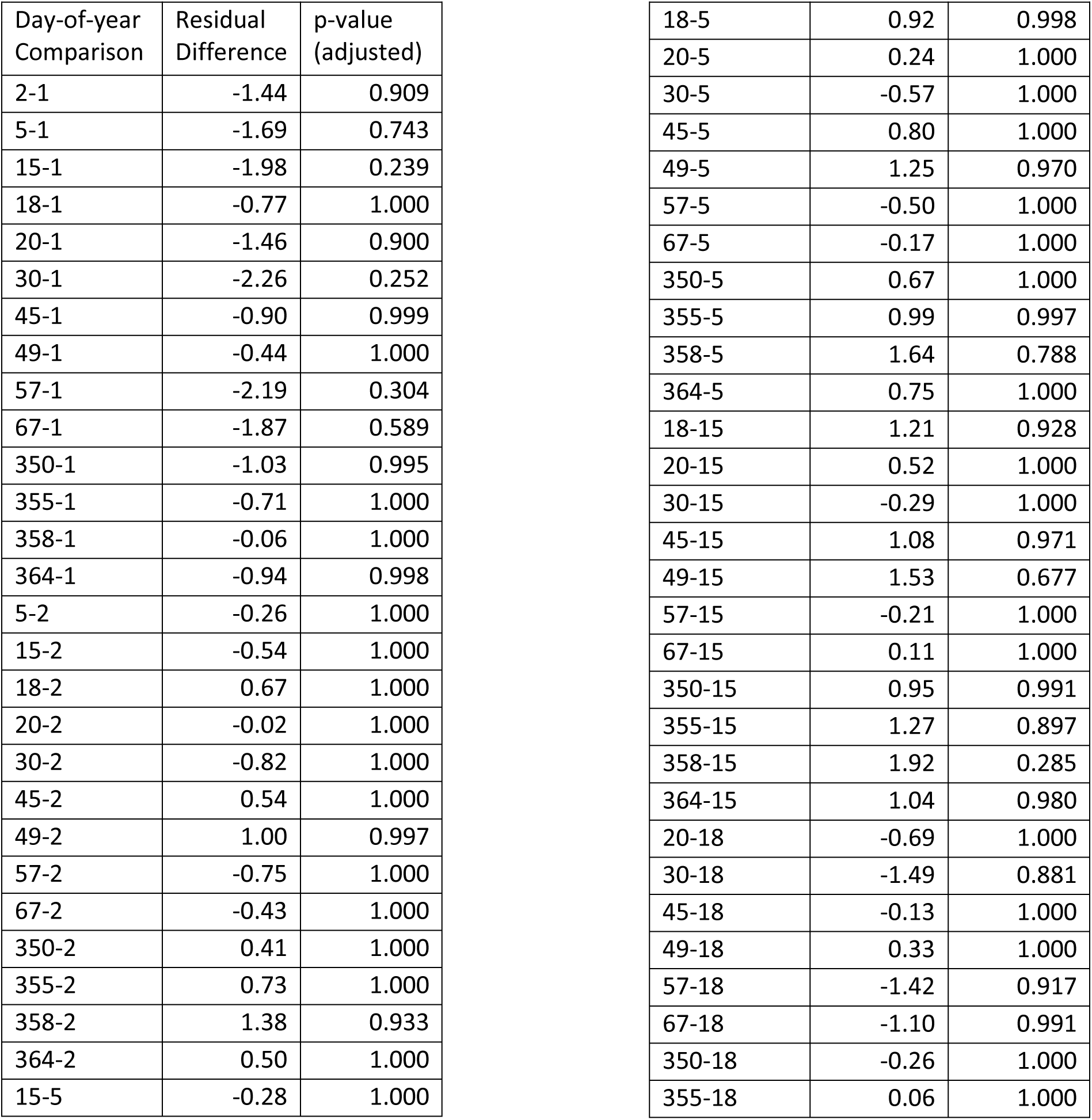
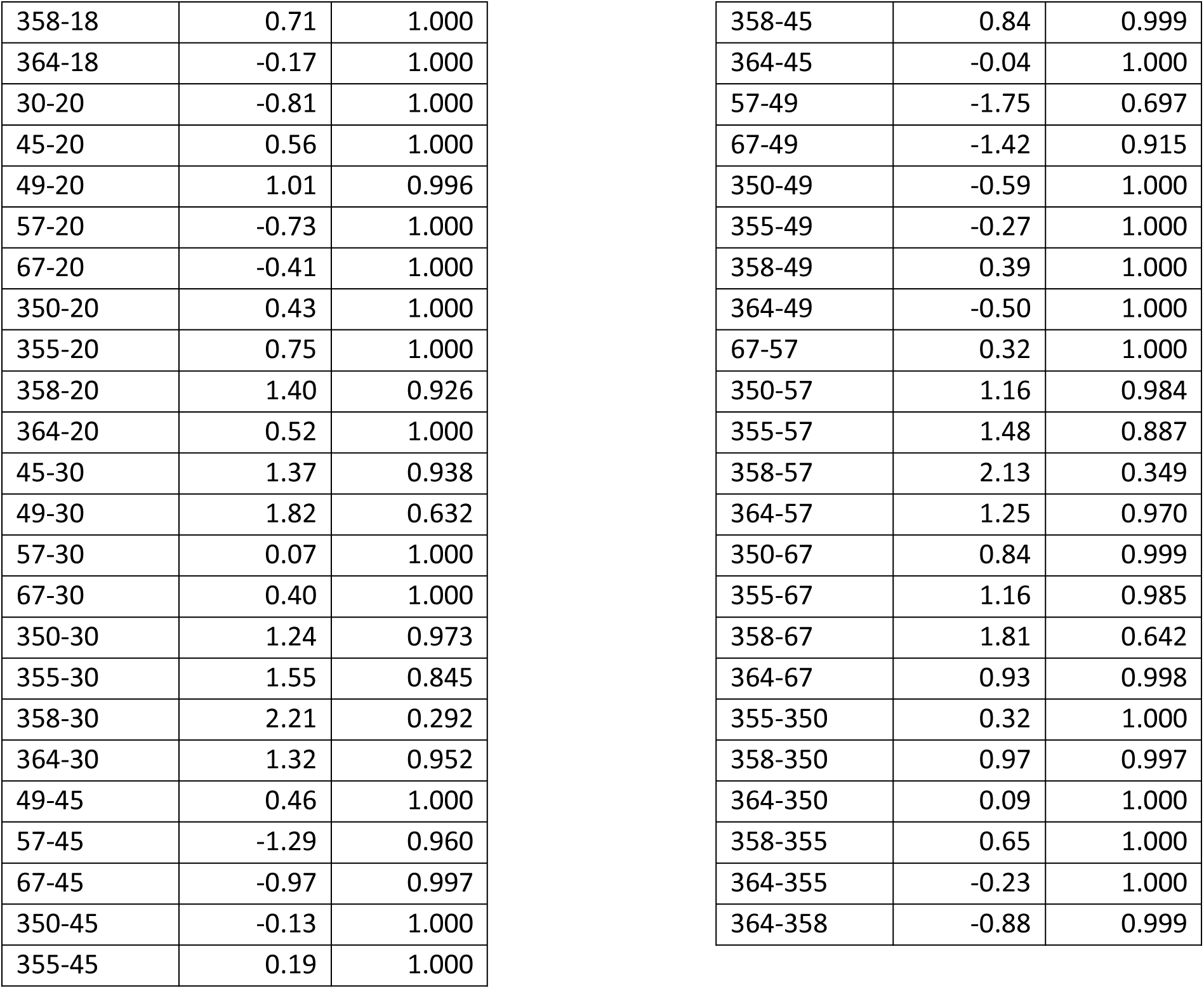
ANOVA for residuals by month. ANOVA for model residuals by month. We examined whether the date of the survey affected model residuals. “Day-of-year Comparison” indicates the two compared dates, and “Residual Difference” indicates the difference in mean residuals between the two dates. The p-value was adjusted using Tukey’s Honest Significant Difference test. All p-values are > 0.23, showing that residuals do not differ between any pairs of dates. These same data are visualized in Fig. S3.

**Table S3.**
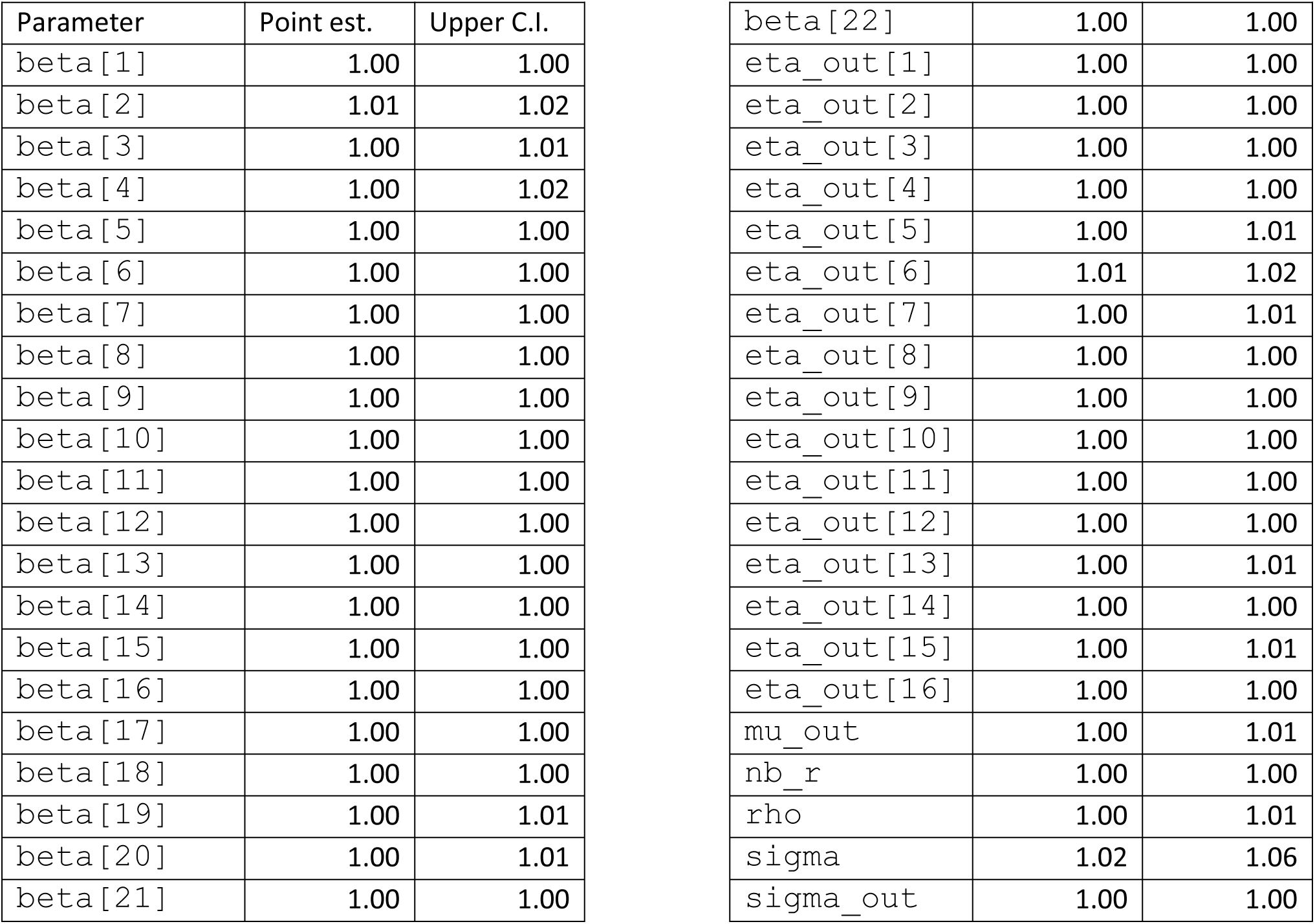
Gelman-Rubin Statistics. Gelman-Rubin statistics for MCMC convergence diagnosis. Values for potential scale reduction factors (PSRFs) close to 1 indicate model convergence. All PSRFs for top-level parameters were estimated to be < 1.02, indicating good convergence. Parameter names match the NIMBLE code, which is available on GitHub and archived on Zenodo (doi:10.5281/zenodo.6687905). The column “Point est.” gives the estimated PSRF, and the column “Upper C.I.” gives the upper bound on the PSRF.

### Appendix S9 Supplemental Figures

**Figure S1.**
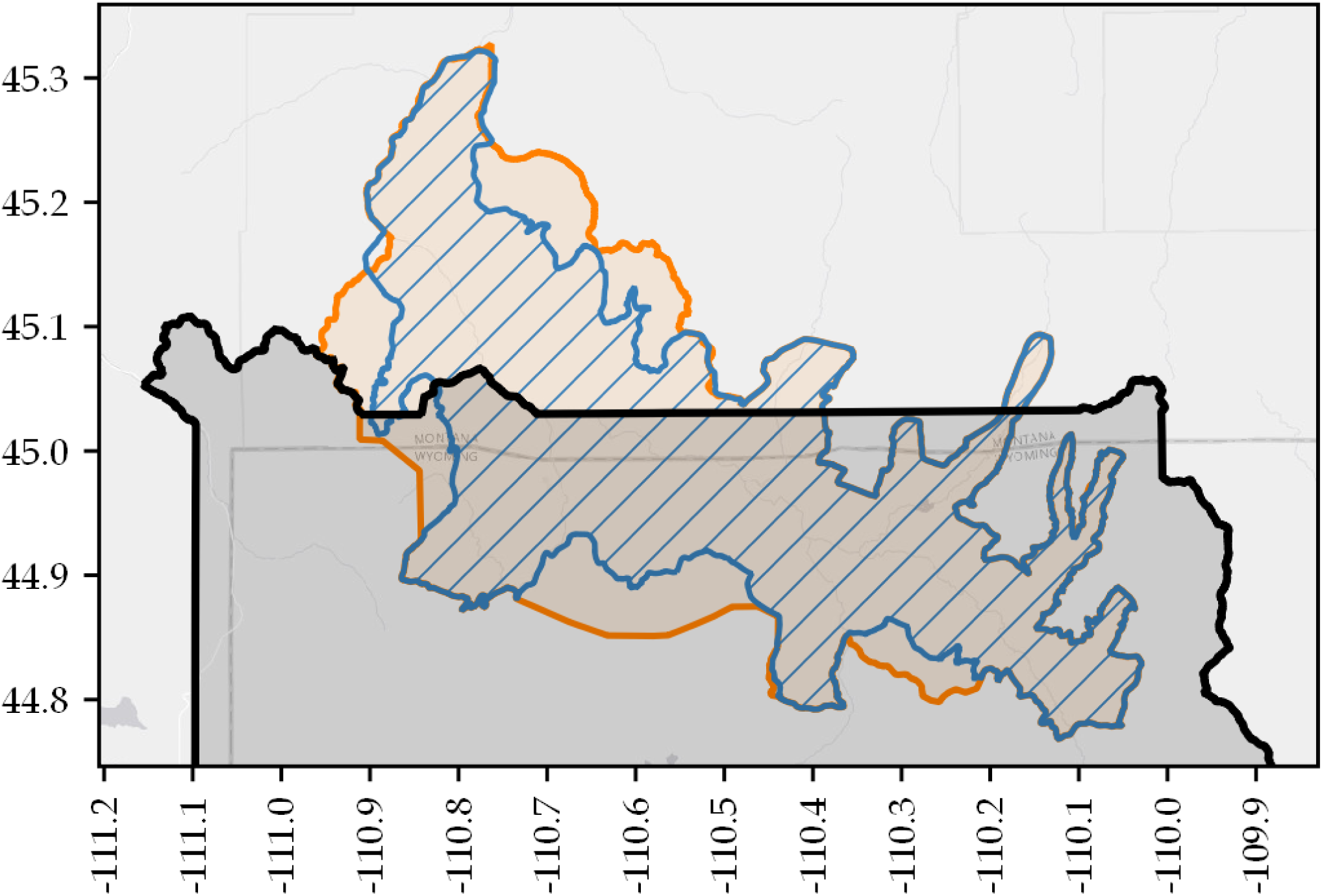
Study area map. Study area showing historic northern Yellowstone elk winter range polygon (blue; 1,520 km^2^), and our adjusted winter range polygon (orange; 1,900 km^2^), and the boundary of Yellowstone National Park (black).

**Figure S2.**
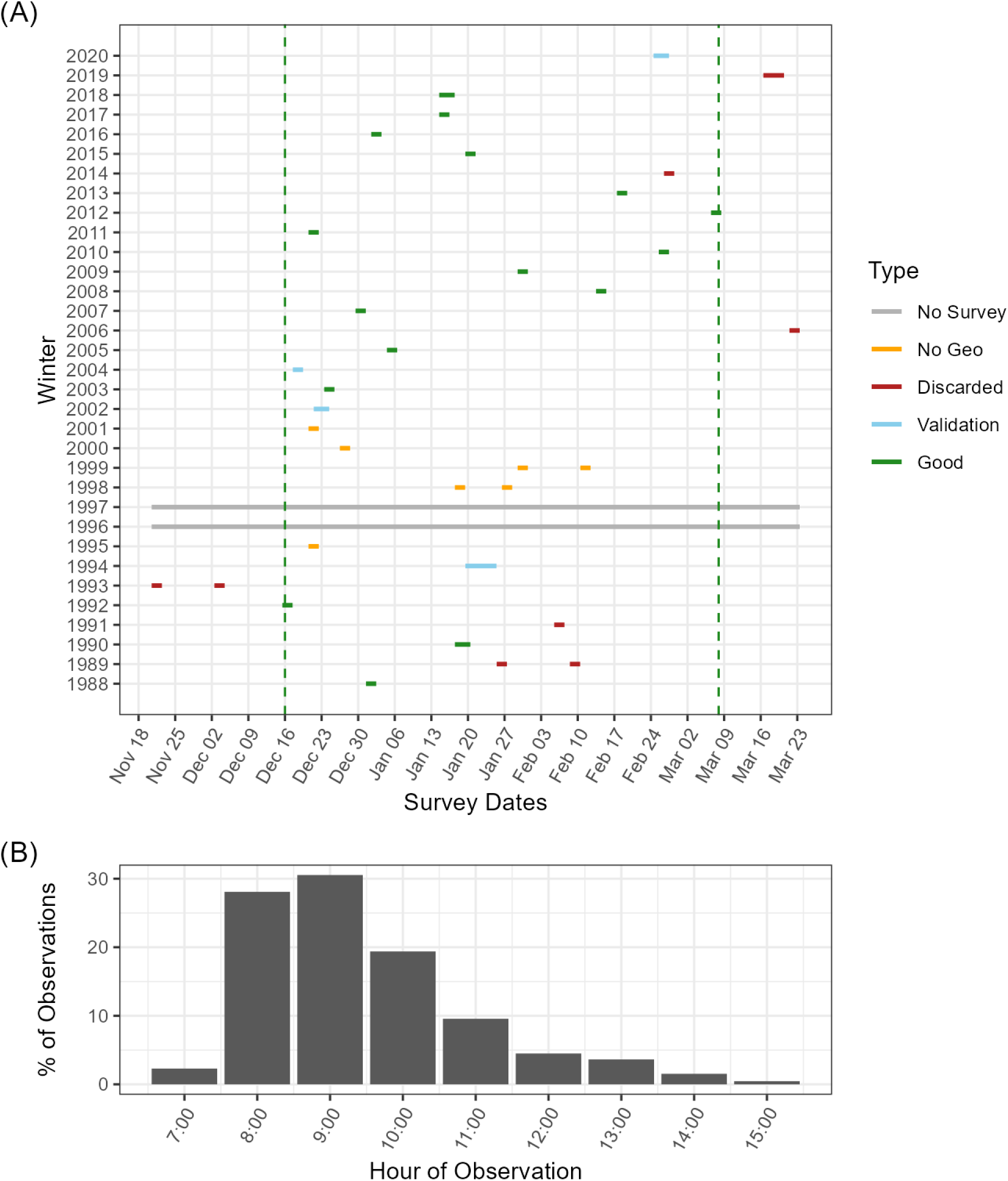
Survey dates and times. We fit our model using data collected from 1988 to 2020. Locations of elk groups (≥ 1 elk) and their sizes were recorded during aerial surveys from fixed-wing aircraft. (A) Surveys used in the model were conducted between December and March in each year. In 1996 and 1997, no survey occurred (gray horizontal lines). Counts for 1989, 1991, 2006, and 2014 are considered unreliable as a census of the population and so were discarded (red lines). The count for 2019 was conducted via helicopter late in the season and was also discarded (red ine). Georeferenced group data are unavailable between 1995 and 2001 and could not be included in the model (orange). In 1994, 2002, and 2004, georeferenced group data are only available inside of Yellowstone National Park (data from the State of Montana were not georeferenced); these data were not included for model fitting but were included for model validation (blue lines). A full census was available for 2020, but we chose to withhold it as an additional validation year with complete georeferencing (blue line). All other years were used for model fitting (green lines). All surveys used in modeling fall between December 16 and March 7 (vertical green dashed lines). (B) Surveys were conducted primarily in the morning hours, and 92% of group observations were recorded between 08:00 and 12:00.

**Figure S3.**
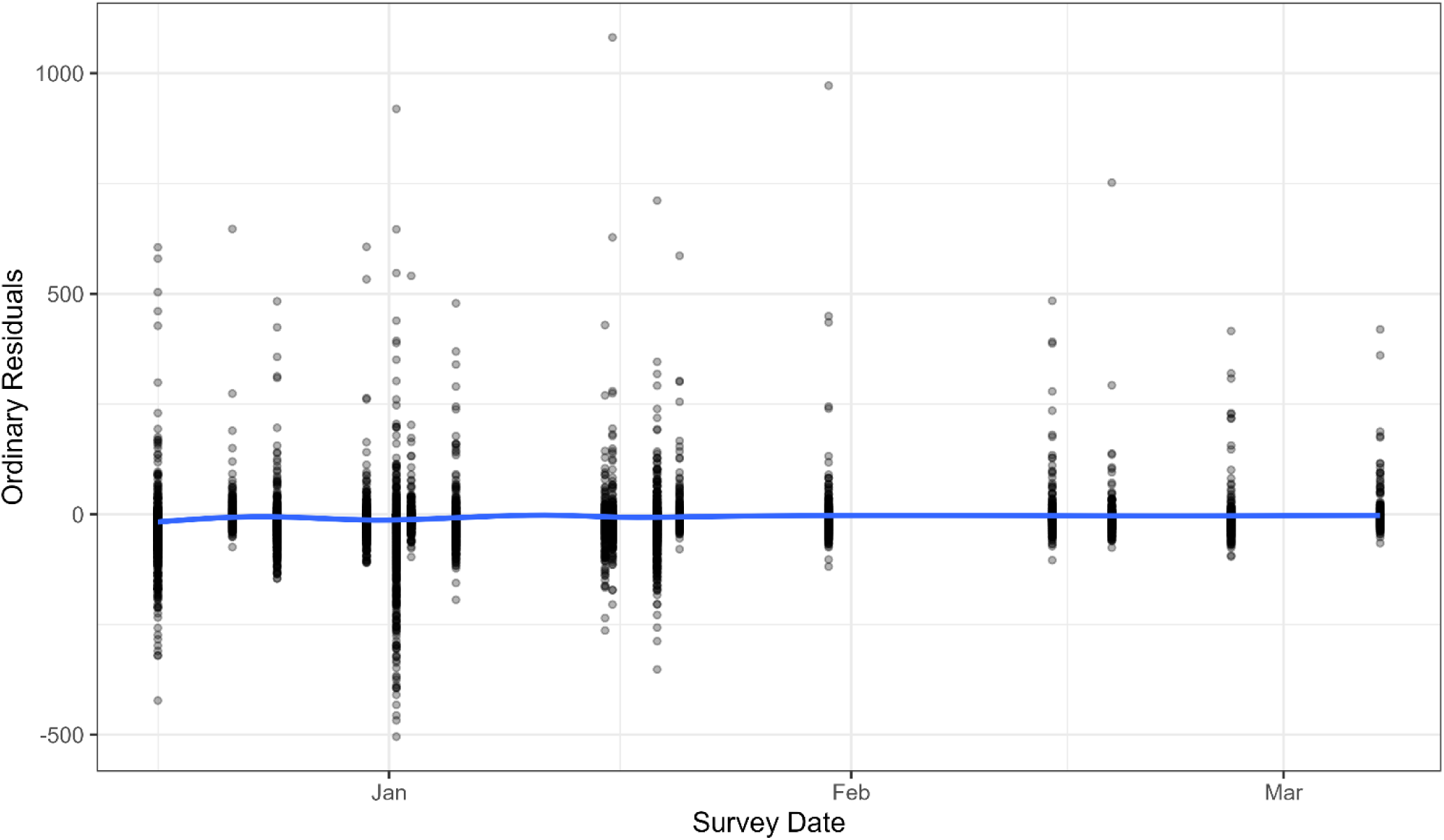
Residuals by survey month. Ordinary model residuals by survey month. We checked that variable survey dates (Fig. S2A) did not introduce a bias in our model. We plotted the model’s ordinary residuals as a function of survey date, and visually checked for any pattern by fitting a very flexible GAM smoothing line, which could have picked up subtle non-linear differences by date (blue line). The fitted line shows no obvious pattern, which was confirmed by an ANOVA and Tukey’s HSD test (Table S2).

**Figure S4.**
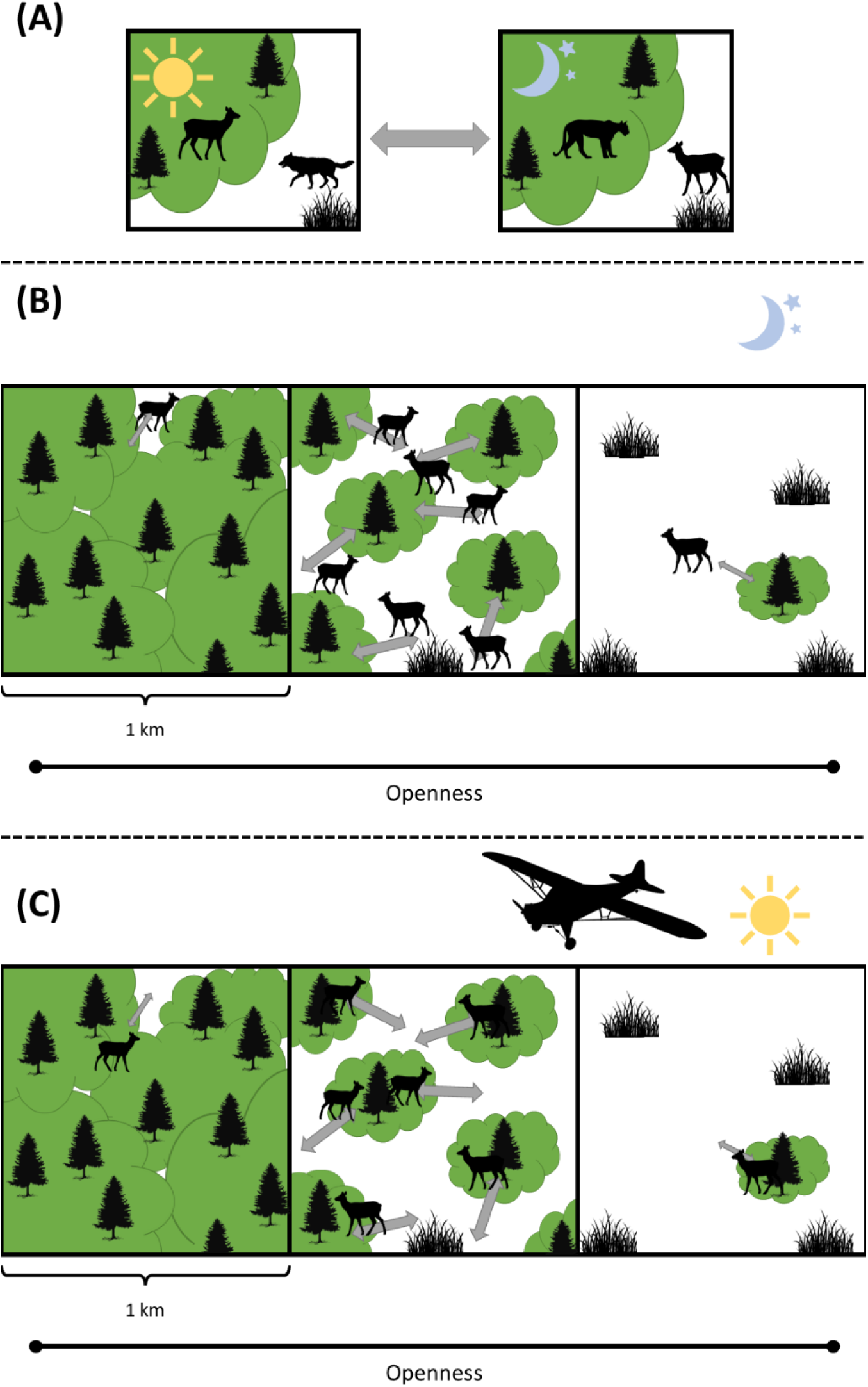
Scale of analysis. Spatial scale of analysis. We rasterized all counts and covariates on the same 1 km x 1 km raster grid. We chose this coarse spatial scale to average over daily elk movements and capture the population-level distribution. Elk are known to move between habitats characterized by openness (depicted here) and roughness (not depicted) at different times of day to manage their risk from both wolves and cougars. In terms of openness, risk from wolves is greatest in habitats with little or no tree cover, whereas risk from cougars is greatest in forested habitats. (A) Elk spend daytime hours, when wolves are more active, in forested habitats (left panel), whereas they spend nighttime hours, when cougars are more active, in open habitats (right panel). (B and C) In one-km pixels across a gradient of varying openness, elk are limited by the heterogeneity of the pixel, with gray bidirectional arrows representing diel shuttling. The gray arrows are identical between (B) and (C), but (B) shows the nighttime position of elk and (C) shows the daytime position of elk. At the 1-km scale, we hypothesized that elk density would be greatest in habitats with intermediate openness (middle panel of B and C), which provide multiple patches that facilitate his diel shuttling behavior. Aerial surveys occur during daytime (C).

**Figure S5.**
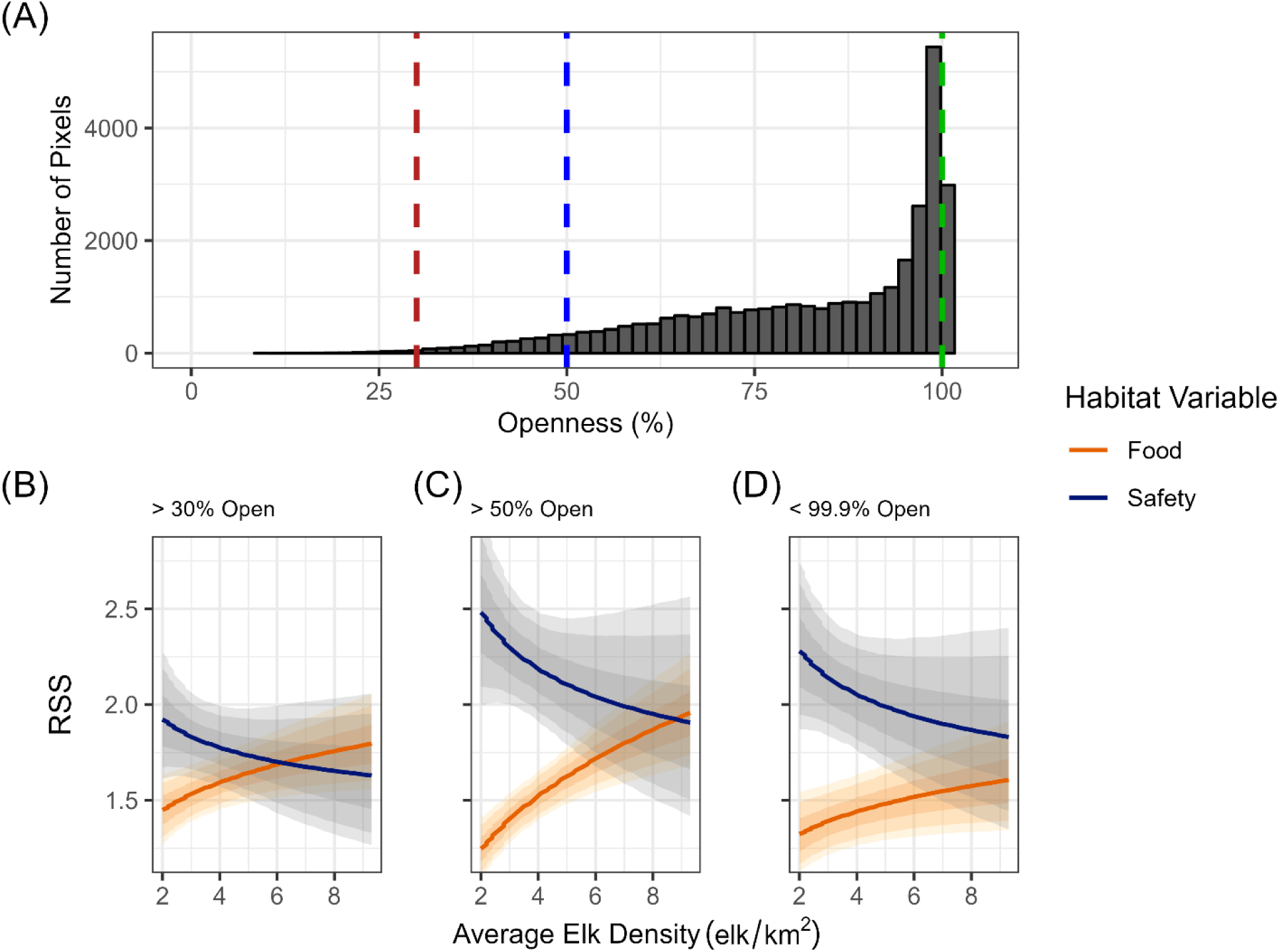
Openness sensitivity analysis. We conducted a sensitivity analysis to assess the potential impact of imperfect detection of elk due to low openness (high forest canopy cover) on our results. We discarded any pixels from the landscape that fell outside a particular openness cutoff in any year and refitted he model. (A) We defined three openness cutoffs, where the pixels retained for model fitting were >30% open (95% of original data retained; red dashed line), >50% open (70% of original data retained; blue dashed line), and <99.9% open (73% of original data retained; green dashed ine). (B) Results for the >30% cutoff, (C) >50% cutoff, and (D) <99.9% cutoff were qualitatively similar to the results presented in the main ext: we found positive DDHS for food, negative DDHS for safety, and a switch in their relative importance. DDHS for food equals the RSS for a 1-SD change in herbaceous biomass, and DDHS for safety equals a 0.5-SD change in openness and a 0.5-SD change in roughness. See main text for details. In panels (B) – (D), solid lines are posterior mean estimates and shaded envelopes are 50%, 80%, and 90% credible ntervals.

**Figure S6.**
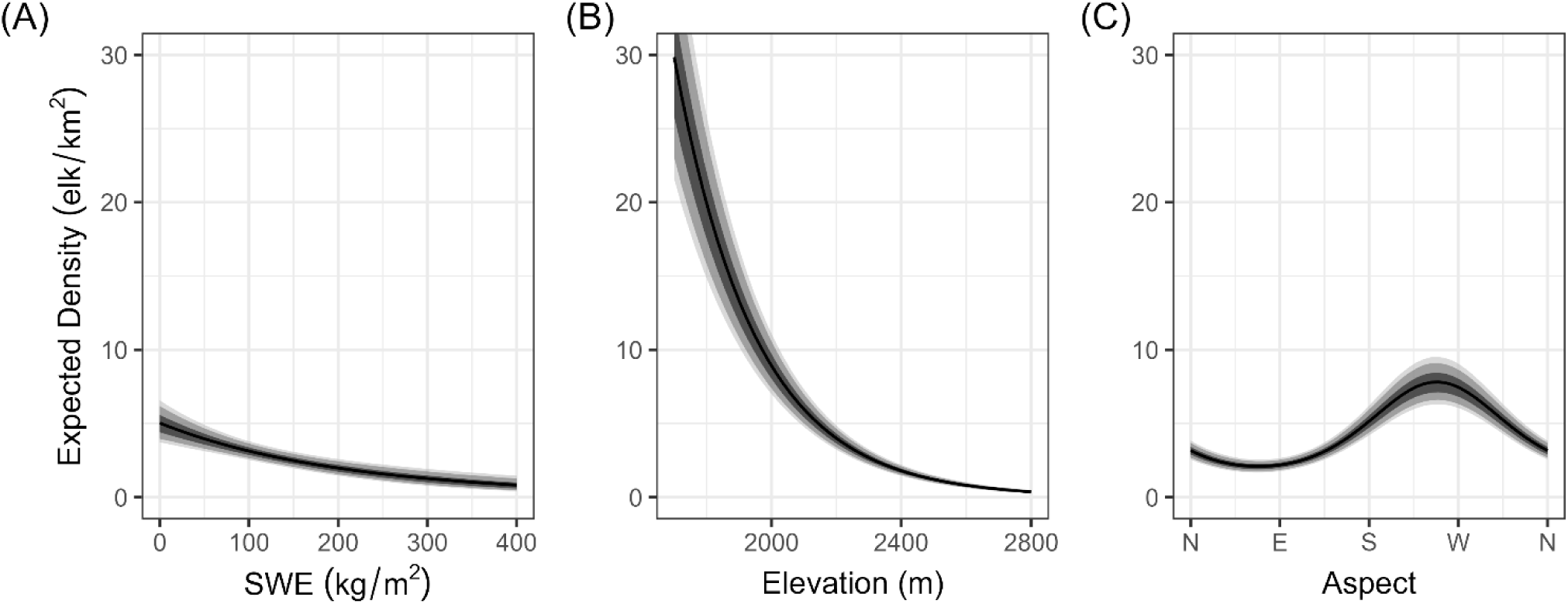
Mean effect of conditions. Mean effect of snow-water equivalent (SWE), elevation, and aspect on expected elk density during winter (December-February), with all other variables held at their mean. (A) Elk density was greatest in pixels with low snow, (B) low elevation, and (C) southwest aspects. Solid black lines show the mean effects, and the shaded gray envelopes show the 50%, 80%, and 90% credible intervals.

**Figure S7.**
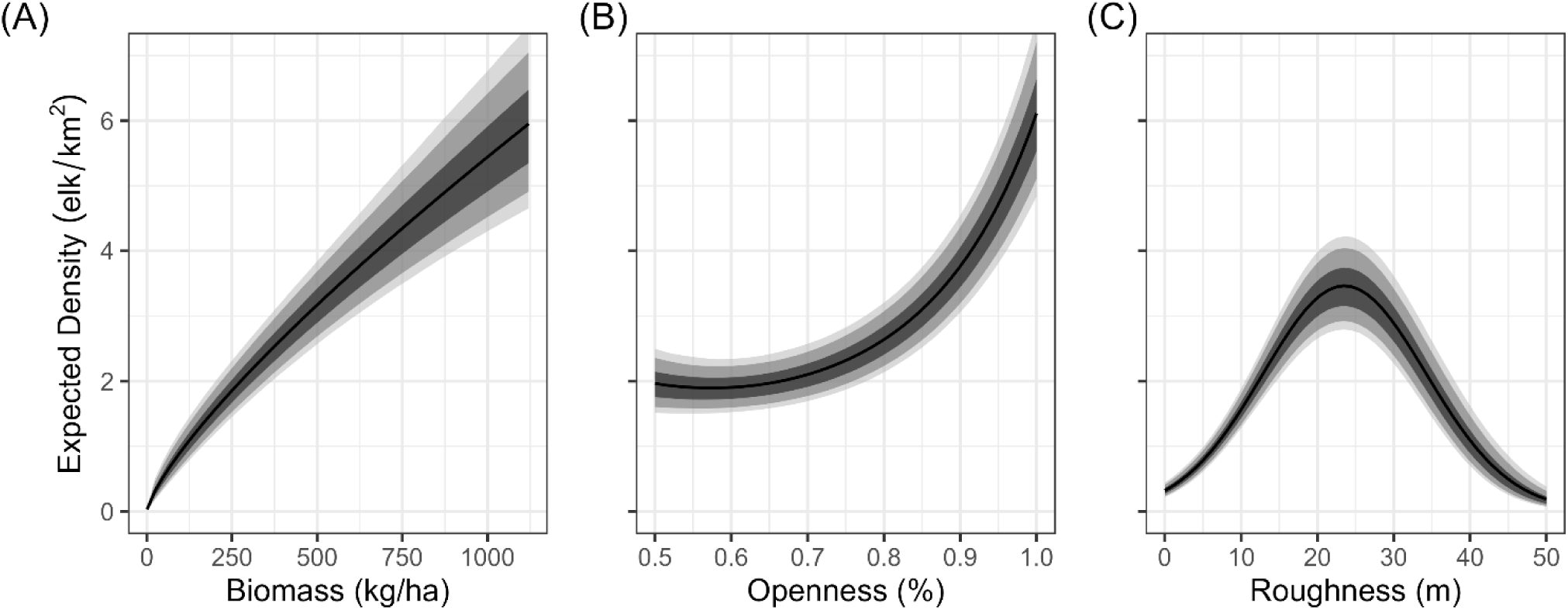
Mean effect of food and safety variables. Mean effects of food and safety variables on expected elk density, with all other variables held at their overall mean. (A) Expected elk density increased with herbaceous biomass, consistent with our assertion that it is an important food resource for elk. (B) Contrary to our prediction that elk would most prefer an intermediate openness, expected elk density was greatest at 100% openness, and the relationship between openness and elk density was largely monotonic for the observed range of openness. (C) Expected elk density was greatest for a roughness of 23.5 m (intermediate, as expected). Solid black lines show the mean effects, and the shaded gray envelopes show the 50%, 80%, and 90% credible intervals.

**Figure S8.**
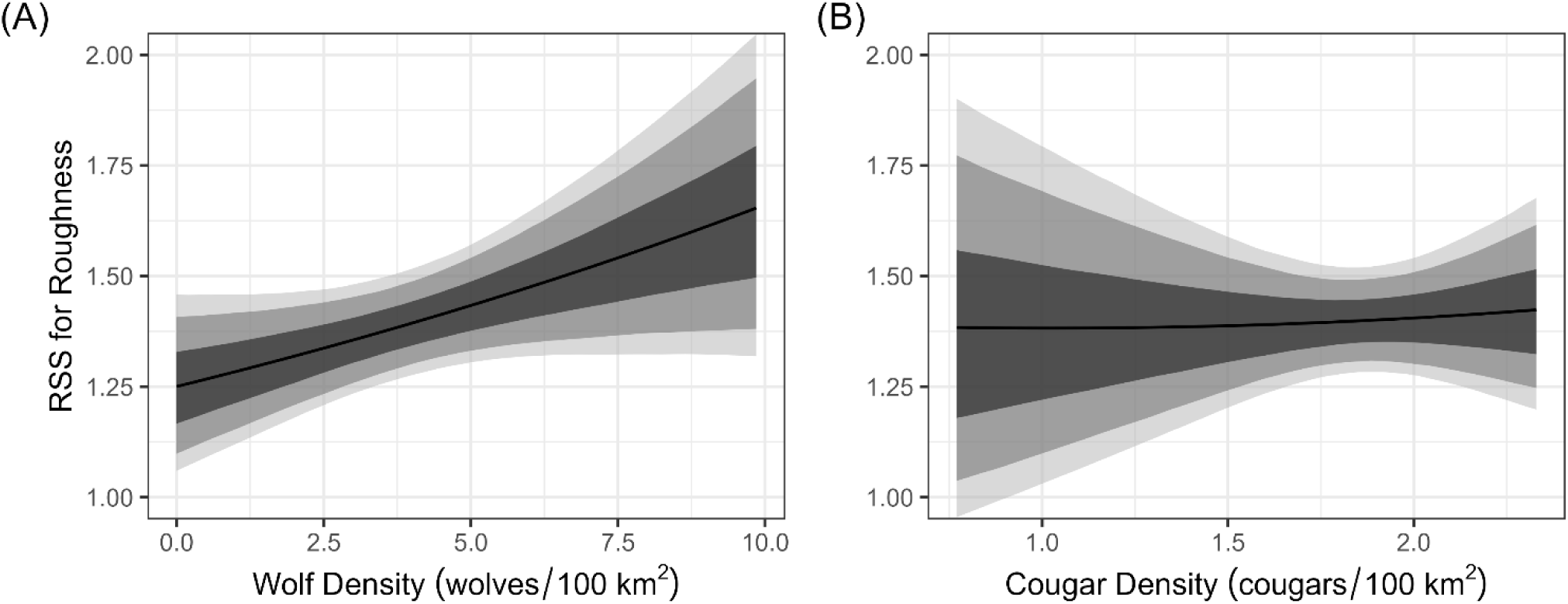
Effect of predator density on RSS for roughness. Mean effect of predator density on relative selection strength (RSS) for roughness, with all other variables held at their mean. (A) RSS for roughness increased with wolf density, in agreement with our prediction that elk would move away from the wolf habitat domain. (B) RSS for roughness did not change with cougar density, in agreement with our prediction that the wolf effect would be stronger than the cougar effect during the daytime aerial elk surveys. Solid black lines are mean effects, and shaded gray envelopes are 50%, 80%, and 90% credible intervals.

**Figure S9.**
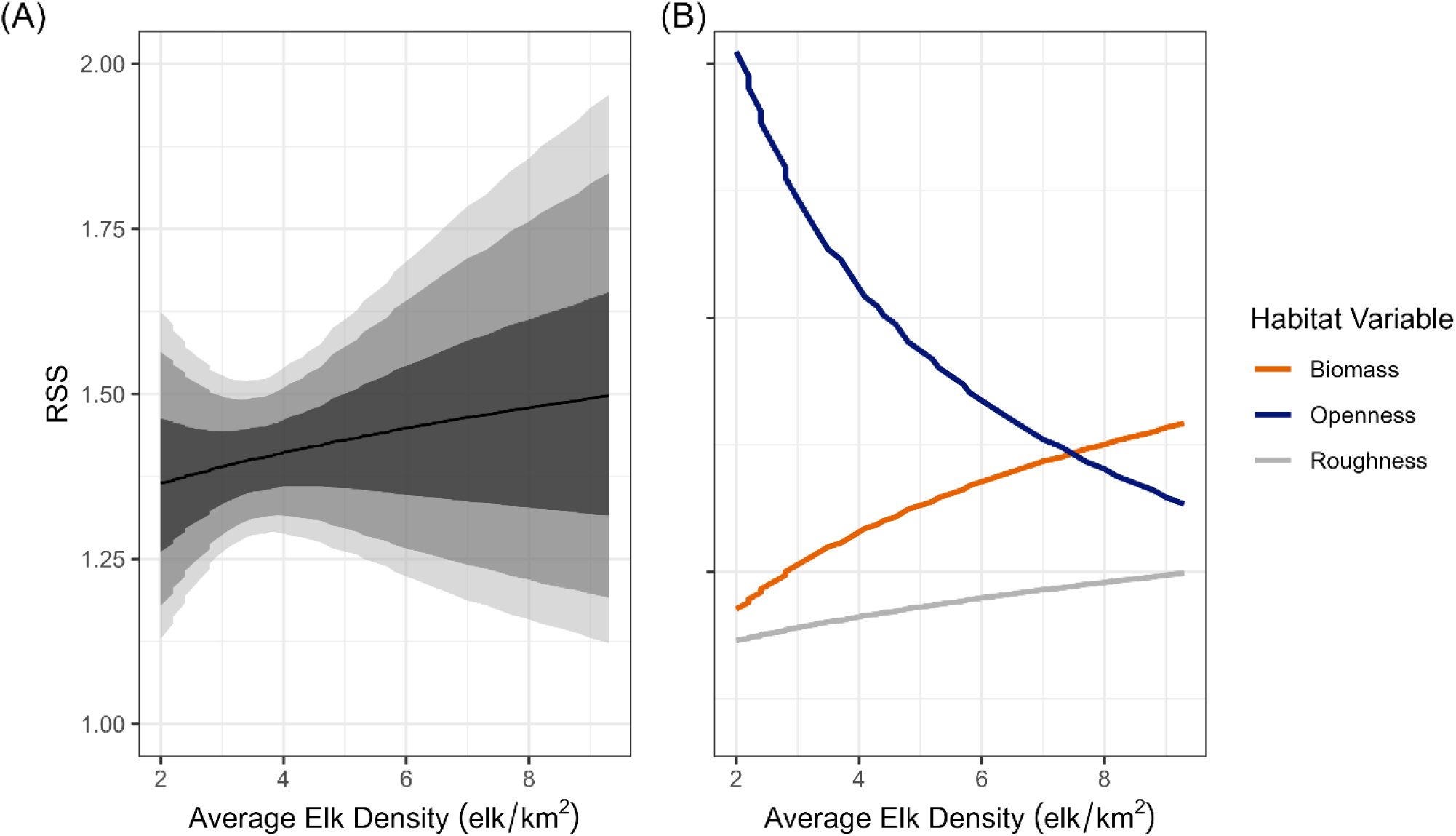
DDHS for roughness. Density-dependent habitat selection (DDHS) for food and safety variables decomposed. (A) Average elk density versus relative selection strength (RSS) for a 1-SD change in roughness. The mean effect is weakly positive, but the large uncertainty indicates no support for DDHS for roughness. RSS was calculated using samples from the entire posterior distribution. Solid black line is the mean effect, and he shaded gray envelopes are the 50%, 80%, and 90% credible intervals. (B) Comparison of RSS for a 1-SD change in all food and safety variables.

**Figure S10.**
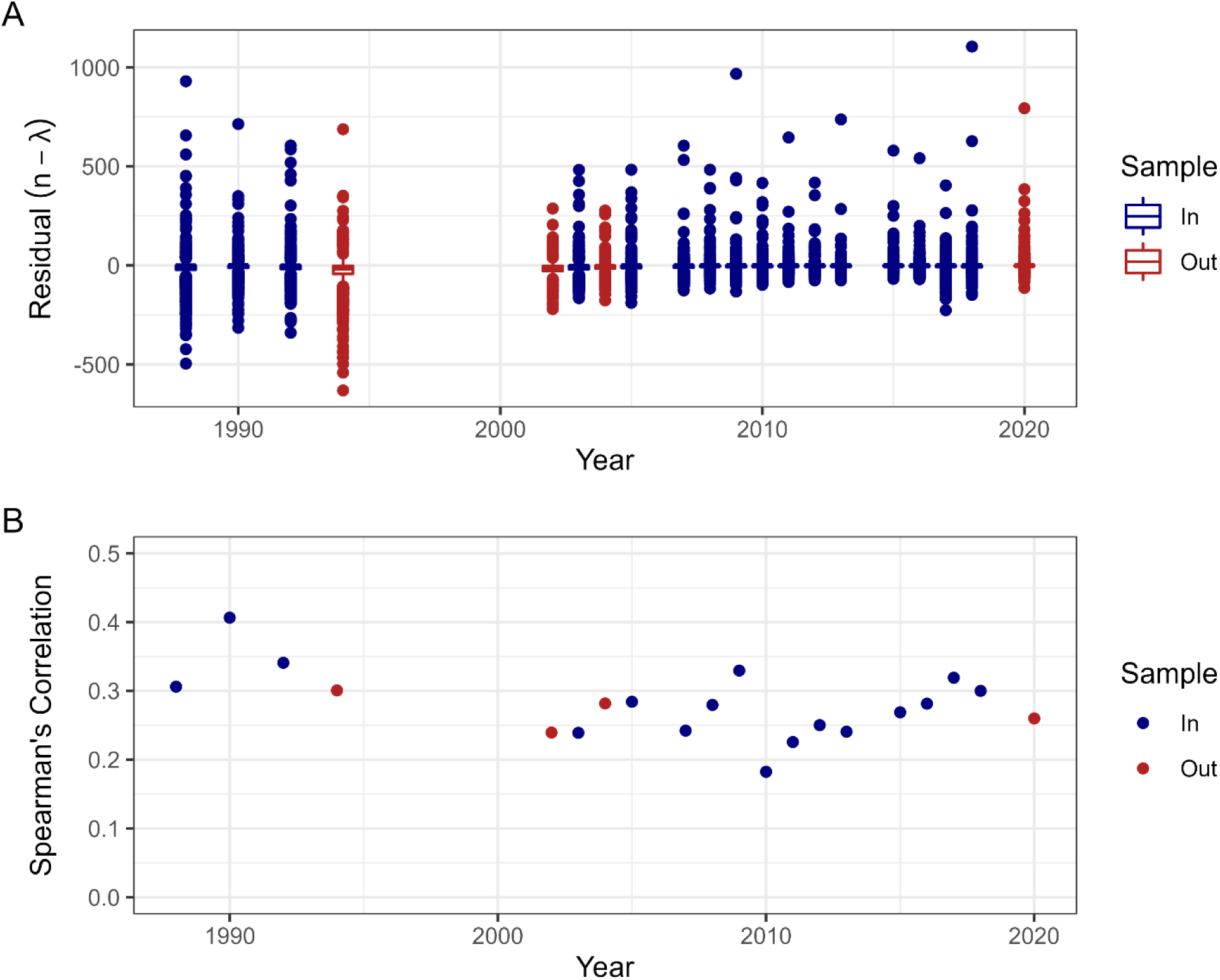
Model evaluation. Model evaluation. We compared (A) ordinary residuals and (B) Spearman’s correlation between expected and observed densities for both in-sample data (blue) and out-of-sample validation data (red). Residuals have mean near 0 in all years, showing good accuracy, but a wide spread, showing low precision. Spearman’s correlation is moderate in all years (mean in-sample = 0.28, mean out-of-sample = 0.27), which shows the model has moderate power to rank pixels by abundance in both training and testing data.

**Figure S11.**
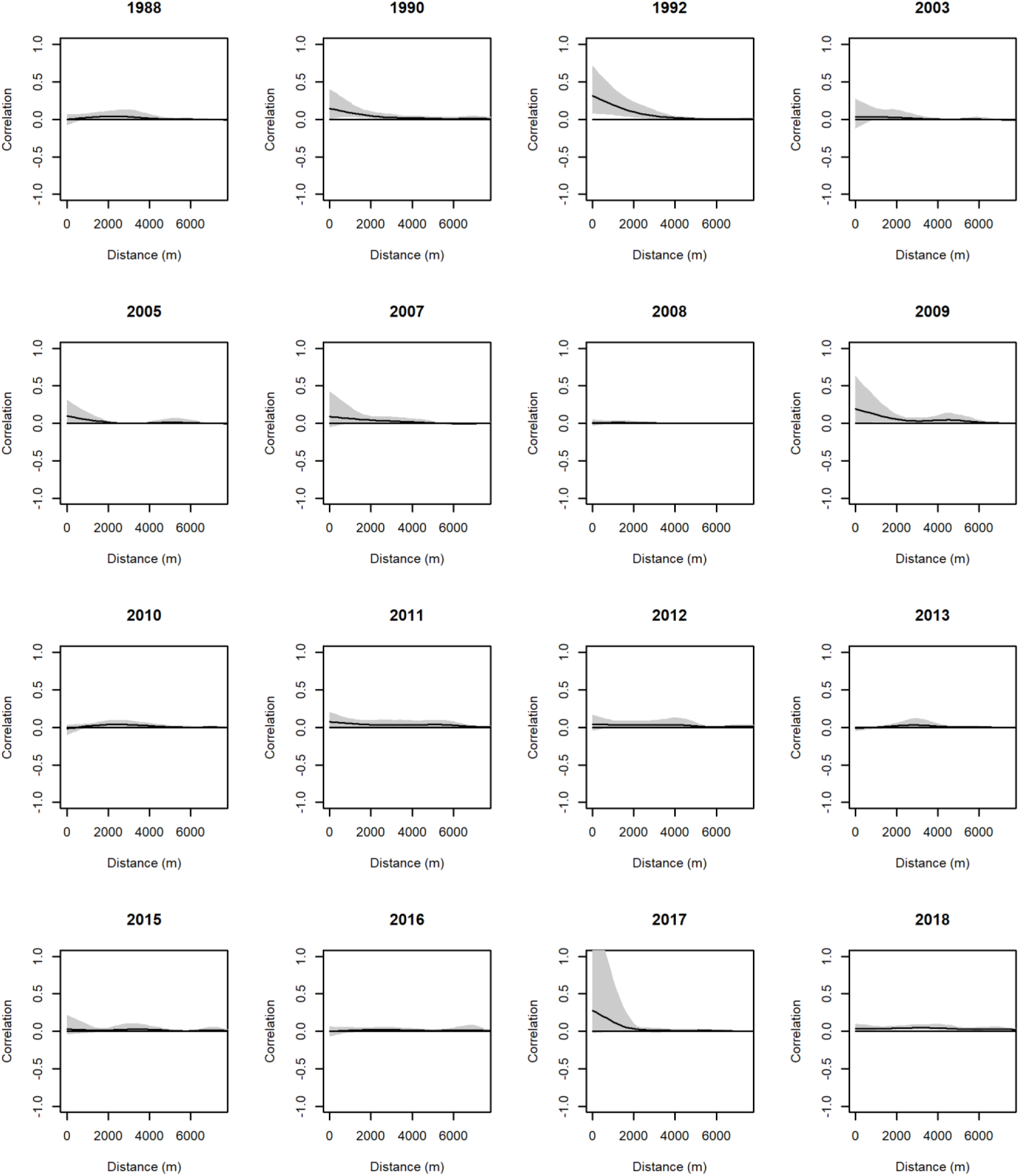
Residual spatial autocorrelation. Non-parametric spline correlograms used to assess residual spatial autocorrelation. We estimated correlograms using Pearson’s residuals for each pixel in each year. The scale of spatial autocorrelation is estimated as the distance where the confidence envelope first intersects the x-axis (correlation = 0). There was little to no residual spatial autocorrelation in all years, i.e., the confidence envelope crosses the x-axis near 0 m in nearly all years (1992 was the lone exception).

**Figure S12.**
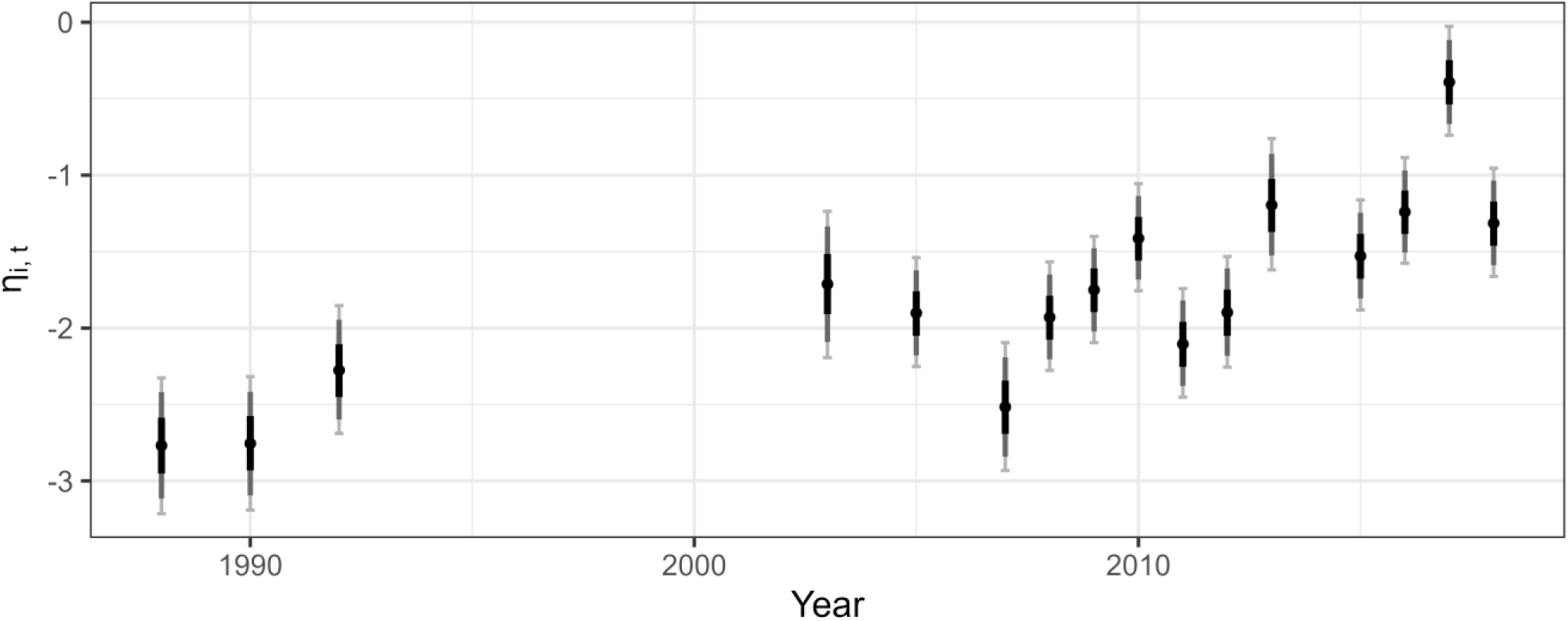
Temporal random effect posterior estimates. Temporal random effect (*η_i,t_*) over time. We included the temporal random effect to account for changing patterns of elk distribution inside versus outside Yellowstone National Park (YNP), along with a change in the late hunt outside of YNP after 2010. The emporal random effect affects pixels outside of YNP, so negative values indicate that pixels outside of YNP have lower expected elk density han would be predicted based on fixed effects alone. Points show posterior mean for each coefficient and bars show credible intervals. Black bars show 50% credible intervals, dark gray bars show 80% credible intervals, and light gray bars with end caps show 90% credible intervals.

**Figure S13.**
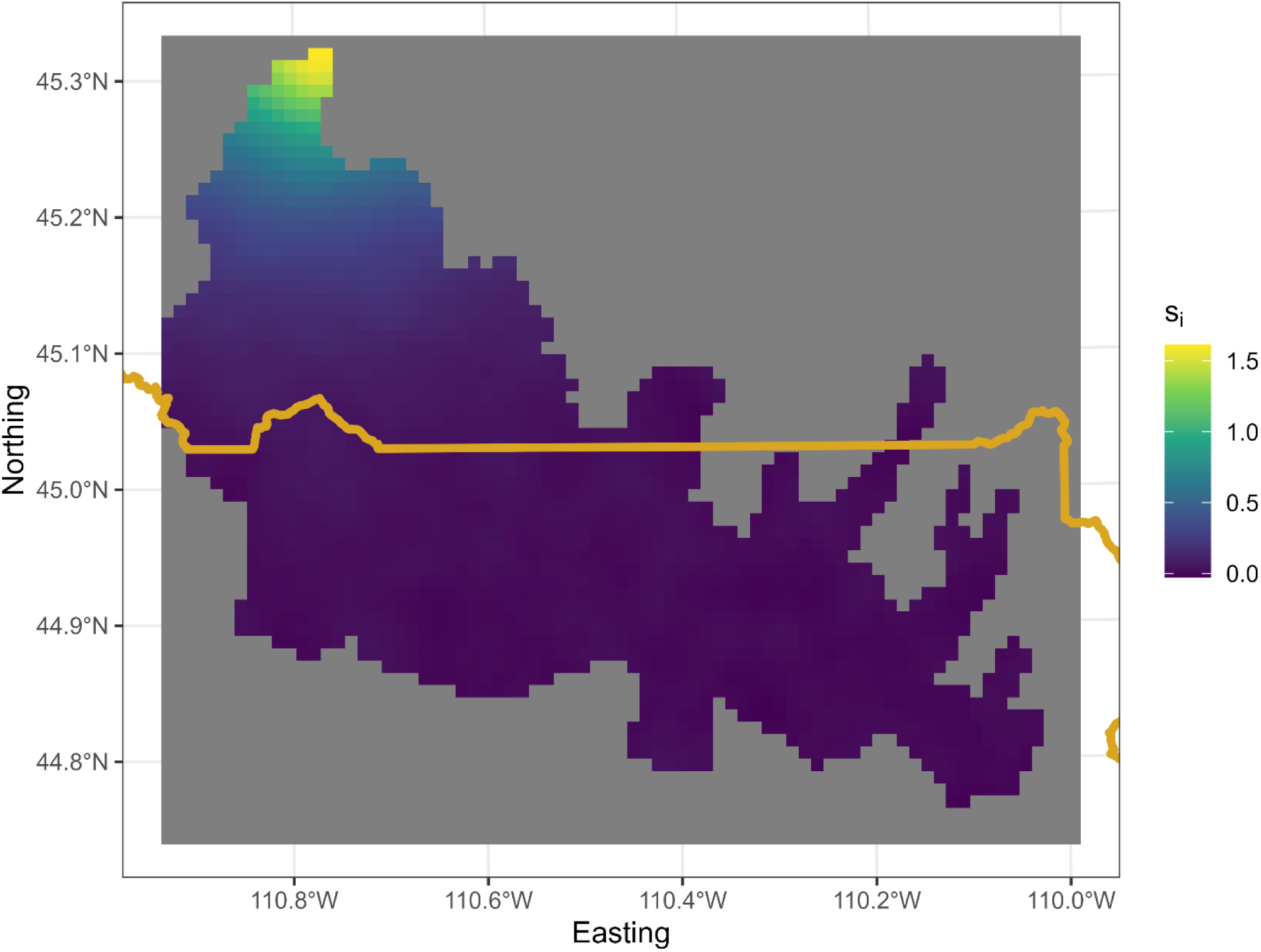
Spatial random effect posterior mean. Posterior mean of estimated spatial random effect, ***s***.

**Figure S14.**
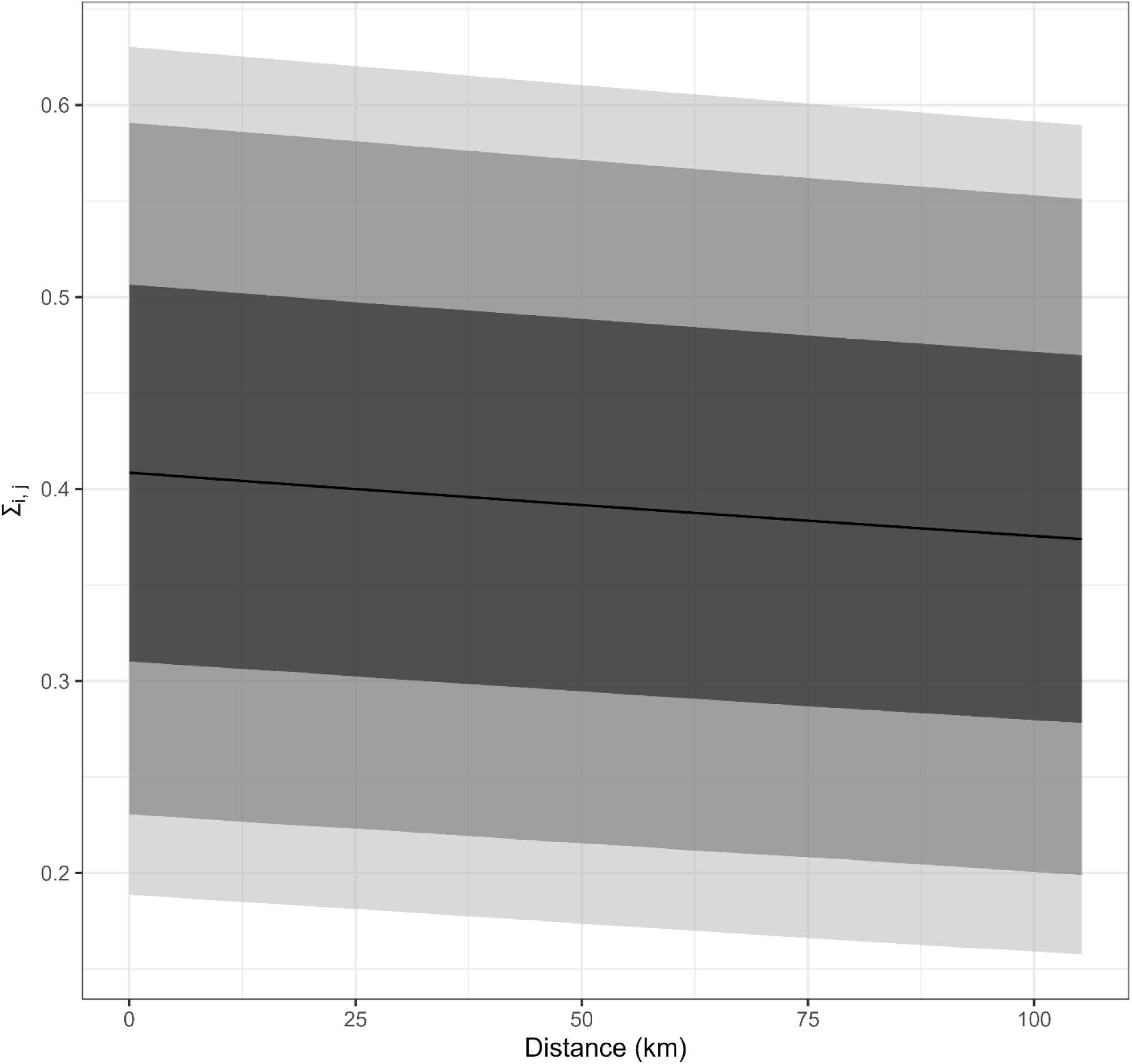
Fitted spatial covariance function. Fitted spatial covariance as a function of distance for the spatial random effect.

**Figure S15.**
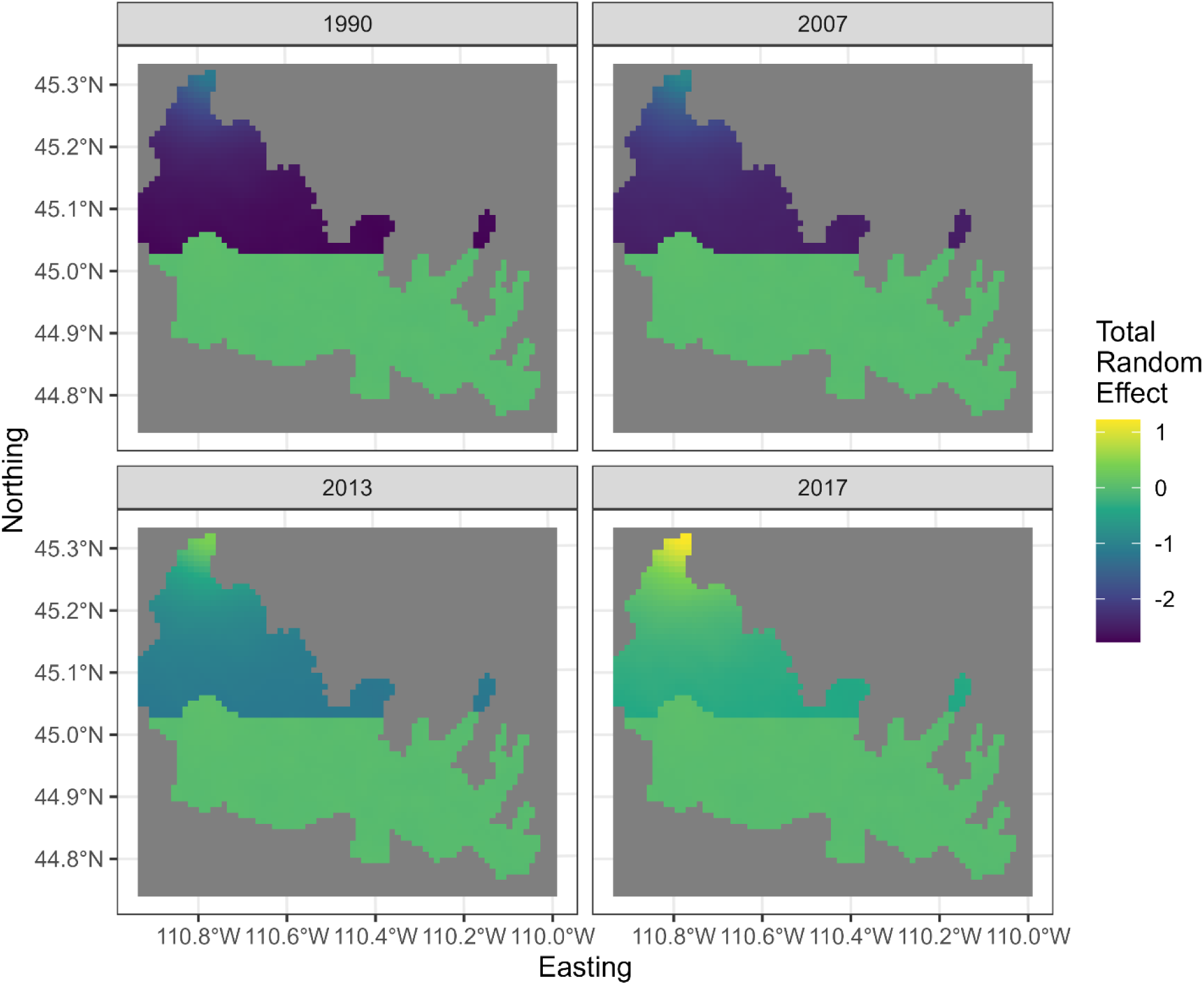
Combined temporal and spatial random effects. The temporal random effect was defined as operating outside Yellowstone National Park (YNP) and the estimated values were always negative. The spatial random effect could only take on positive values, but it could operate anywhere in the study area. The estimated values were largest outside of YNP. The negative effect of the temporal random effect and the positive effect of the spatial random effect hus acted in opposite directions and almost entirely outside YNP, with expected elk density inside YNP almost completely captured by fixed effects. Examples of the combined effect for 1990, 2007, 2013, and 2017. See Fig. S12 for temporal random effect alone in each year and Fig. S13 for spatial random effect alone.

**Figure S16.**
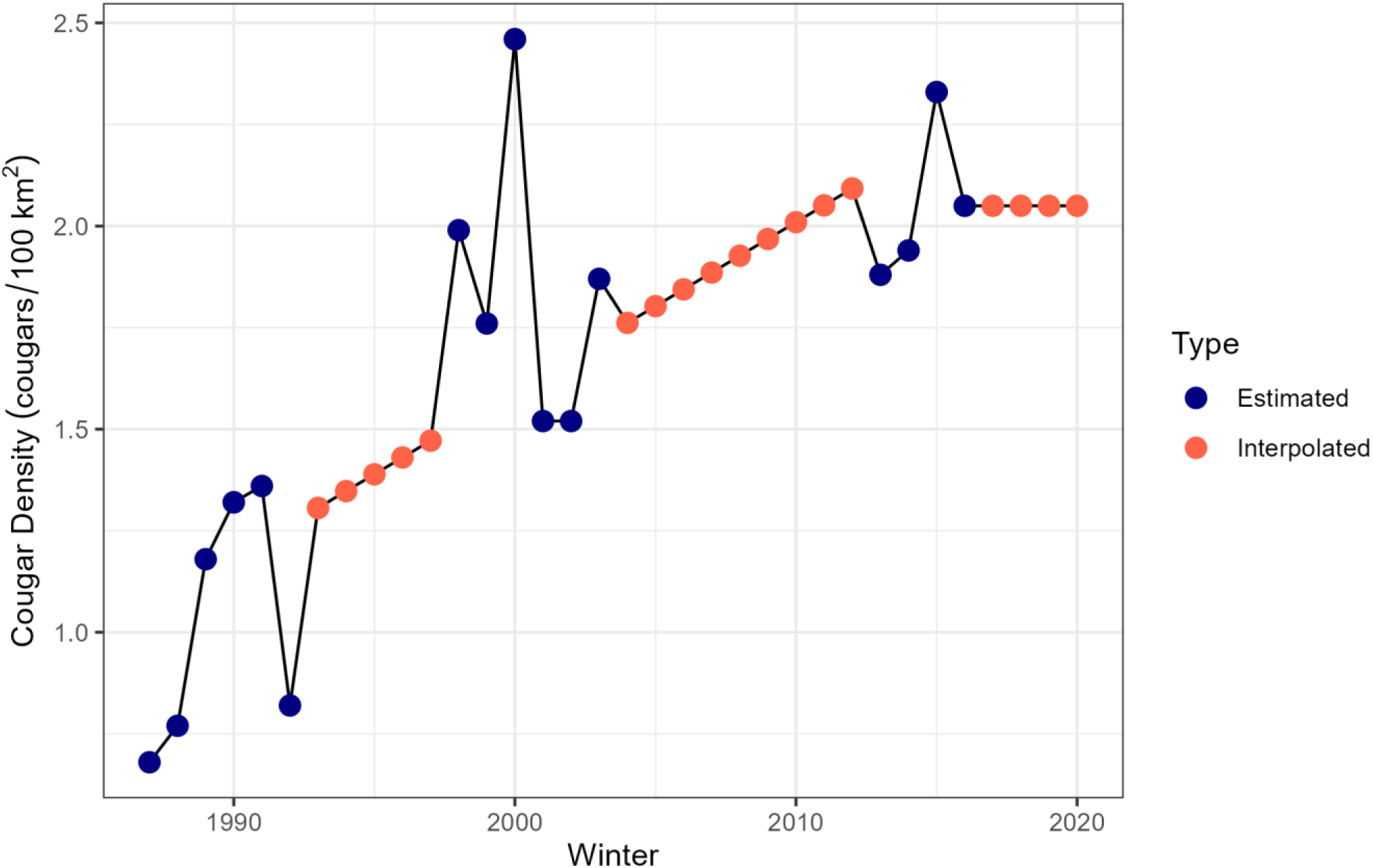
Estimated and interpolated cougar densities. Cougar densities were estimated from field surveys in three phases: Phase 1 (1987 – 1993; Ruth *et al*. 2019), Phase 2 (1998 – 2004; Ruth *et al*. 2019), and Phase 3 (2014 – 2017; Anton 2020). Values between Phases 1 and 2 and between Phases 2 and 3 were linearly interpolated from the entire dataset. Values after Phase 3 (2018 – 2020) were assigned the mean of Phase 3.

## Literature Cited

Abadi, F., Gimenez, O., Jakober, H., Stauber, W., Arlettaz, R. & Schaub, M. (2012). Estimating the strength of density dependence in the presence of observation errors using integrated population models. Ecol. Modell., 242, 1–9.

Avgar, T., Betini, G.S. & Fryxell, J.M. (2020). Habitat selection patterns are density dependent under the ideal free distribution. J. Anim. Ecol., 89, 2777–2787.

Avgar, T., Lele, S.R., Keim, J.L. & Boyce, M.S. (2017). Relative Selection Strength: Quantifying effect size in habitat- and step-selection inference. Ecol. Evol., 7, 5322–5330.

Bangs, E.E. & Fritts, S.H. (1996). Reintroducing the gray wolf to central Idaho and Yellowstone National Park. Wildl. Soc. Bull., 24, 402–413.

Bassing, S.B., Devivo, M., Ganz, T.R., Kertson, B.N., Prugh, L.R., Roussin, T., et al. (2022). Are we telling the same story? Comparing inferences made from camera trap and telemetry data for wildlife monitoring. J. Appl. Ecol., e2745.

van Beest, F.M., McLoughlin, P.D., Vander Wal, E. & Brook, R.K. (2014a). Density-dependent habitat selection and partitioning between two sympatric ungulates. Oecologia, 175, 1155–1165.

van Beest, F.M., Uzal, A., Vander Wal, E., Laforge, M.P., Contasti, A.L., Colville, D., et al. (2014b). Increasing density leads to generalization in both coarse-grained habitat selection and fine-grained resource selection in a large mammal. J. Anim. Ecol., 83, 147–156.

Berryman, A. & Turchin, P. (2001). Identifying the density-dependent structure underlying ecological time series. Oikos, 92, 265–270.

Bjørnstad, O.N. & Falck, W. (2001). Nonparametric spatial covariance functions: Estimation and testing. Environ. Ecol. Stat., 8, 53–70.

Le Bourlot, V., Tully, T. & Claessen, D. (2014). Interference versus Exploitative Competition in the Regulation of Size-Structured Populations. Am. Nat., 184, 609–623.

Boyce, M.S., Johnson, C.J., Merrill, E.H., Nielsen, S.E., Solberg, E.J. & van Moorter, B. (2016). Can habitat selection predict abundance? J. Anim. Ecol., 85, 11–20.

Boyce, M.S. & McDonald, L.L. (1999). Relating populations to habitats using resource selection functions. Trends Ecol. Evol., 14, 268–272.

Brennan, A., Cross, P.C. & Creel, S. (2015). Managing more than the mean: Using quantile regression to identify factors related to large elk groups. J. Appl. Ecol., 52, 1656–1664.

Brook, B.W. & Bradshaw, C.J.A. (2006). Strength of evidence for density dependence in abundance time series of 1198 species. Ecology, 87, 1445–1451.

Brown, J.S. & Kotler, B.P. (2004). Hazardous duty pay and the foraging cost of predation. Ecol. Lett., 7, 999–1014.

Dennis, B. & Taper, M.L. (1994). Density Dependence in Time Series Observations of Natural Populations: Estimation and Testing. Ecol. Monogr., 64, 205–224.

Fieberg, J., Signer, J., Smith, B. & Avgar, T. (2021). A “How to” guide for interpreting parameters in habitat-selection analyses. J. Anim. Ecol., 90, 1027–1043.

Fretwell, S.D. & Lucas, H.L. (1969). On territorial behavior and other factors influencing habitat distribution in birds. Acta Biotheor., 19, 16–36.

Gelman, A. & Rubin, D.B. (1992). lnference from Iterative Simulation Using Multiple Sequences. Stat. Sci., 7, 457–472.

Gerard, J.F., Bideau, E., Maublanc, M.L., Loisel, P. & Marchal, C. (2002). Herd size in large herbivores: Encoded in the individual or emergent? Biol. Bull., 202, 275–282.

Greene, C.M. & Stamps, J.A. (2001). Habitat selection at low population densities. Ecology, 82, 2091– 2100.

Guthery, F.S. & Shaw, J.H. (2013). Density dependence: Applications in wildlife management. J. Wildl. Manage., 77, 33–38.

Hooten, M.B. & Hobbs, N.T. (2015). A guide to Bayesian model selection for ecologists. Ecol. Monogr., 85, 3–28.

Houston, D.B. (1982). The northern Yellowstone elk: ecology and management. Macmillan Publishing Co., Inc., New York, NY.

Johnson, D.H. (1980). The Comparison of Usage and Availability Measurements for Evaluating Resource Preference. Ecology, 61, 65–71.

Jones, M.O., Robinson, N.P., Naugle, D.E., Maestas, J.D., Reeves, M.C., Lankston, R.W., et al. (2021). Annual and 16-Day Rangeland Production Estimates for the Western United States. Rangel. Ecol. Manag., 77, 112–117.

Kohl, M.T., Ruth, T.K., Metz, M.C., Stahler, D.R., Smith, D.W., White, P.J., et al. (2019). Do prey select for vacant hunting domains to minimize a multi-predator threat? Ecol. Lett., 22, 1724–1733.

Křivan, V., Cressman, R. & Schneider, C. (2008). The ideal free distribution: A review and synthesis of the game-theoretic perspective. Theor. Popul. Biol., 73, 403–425.

Lehtonen, J. & Jaatinen, K. (2016). Safety in numbers: the dilution effect and other drivers of group life in the face of danger. Behav. Ecol. Sociobiol., 70, 449–458.

Lemke, T.O. & Mack, J.A. (1998). Winter range expansion by the northern Yellowstone elk herd. Intermt. J. Sci., 4, 1–8.

Lima, S.L. & Dill, L.M. (1990). Behavioral decisions made under the risk of predation: a review and prospectus. Can. J. Zool., 68, 619–640.

Ma, Q., Johansson, A. & Sumpter, D.J.T. (2011). A first principles derivation of animal group size distributions. J. Theor. Biol., 283, 35–43.

MacArthur, R.H. & Pianka, E.R. (1966). On Optimal Use of a Patchy Environment. Am. Nat., 100, 603– 609.

MacNulty, D.R., Stahler, D.R., Wyman, C.T., Ruprecht, J.S., Smith, L.M., Kohl, M.T., et al. (2020). Population Dynamics of Northern Yellowstone Elk after Wolf Reintroduction. In: Yellowstone Wolves (eds. Smith, D.W., Stahler, D.R. & MacNulty, D.R.). The University of Chicago Press, Chicago, pp. 184–199.

Marcus, W.A., Meacham, J.E., Rodman, A.W., Steingisser, A.Y. & Menke, J.T. (2022). Atlas of Yellowstone. Second Edi. University of California Press, Oakland, CA.

Matthiopoulos, J., Fieberg, J.R., Aarts, G., Beyer, H.L., Morales, J.M. & Haydon, D.T. (2015). Establishing the link between habitat selection and animal population dynamics. Ecol. Monogr., 85, 413–436.

Matthiopoulos, J., Field, C. & MacLeod, R. (2019). Predicting population change from models based on habitat availability and utilization. Proc. R. Soc. B Biol. Sci., 286.

McLoughlin, P.D., Boyce, M.S., Coulson, T. & Clutton-Brock, T. (2006). Lifetime reproductive success and density-dependent, multi-variable resource selection. Proc. R. Soc. B Biol. Sci., 273, 1449–1454.

McLoughlin, P.D., Morris, D.W., Fortin, D., Vander Wal, E. & Contasti, A.L. (2010). Considering ecological dynamics in resource selection functions. J. Anim. Ecol., 79, 4–12.

Merrill, E., Killeen, J., Pettit, J., Trottier, M., Martin, H., Berg, J., et al. (2020). Density-Dependent Foraging Behaviors on Sympatric Winter Ranges in a Partially Migratory Elk Population. Front. Ecol. Evol., 8, 1–15.

Metz, M.C., Smith, D.W., Vucetich, J.A., Stahler, D.R. & Peterson, R.O. (2012). Seasonal patterns of predation for gray wolves in the multi-prey system of Yellowstone National Park. J. Anim. Ecol., 81, 553–563.

Mobæk, R., Mysterud, A., Egil Loe, L., Holand, Ø. & Austrheim, G. (2009). Density dependent and temporal variability in habitat selection by a large herbivore; an experimental approach. Oikos, 118, 209–218.

Moll, R.J., Redilla, K.M., Mudumba, T., Muneza, A.B., Gray, S.M., Abade, L., et al. (2017). The many faces of fear: a synthesis of the methodological variation in characterizing predation risk. J. Anim. Ecol., 86, 749–765.

Mooring, M.S., Fitzpatrick, T.A., Nishihira, T.T. & Reisig, D.D. (2004). Vigilance, predation risk, and the Allee effect in desert bighorn sheep. J. Wildl. Manage., 68, 519–532.

Morris, D.W. (1987). Tests of Density-Dependent Habitat Selection in a Patchy Environment. Ecol. Monogr., 57, 269–281.

Morris, D.W. (1988). Habitat-dependent population regulation and community structure. Evol. Ecol., 2, 253–269.

Morris, D.W. (1989). Density-dependent habitat selection: Testing the theory with fitness data. Evol. Ecol., 3, 80–94.

Morris, D.W. (2002). Measuring the Allee effect: Positive density dependence in small mammals. Ecology, 83, 14–20.

Morris, D.W. (2003). Toward an ecological synthesis: A case for habitat selection. Oecologia, 136, 1–13.

Morris, D.W. (2011). Adaptation and habitat selection in the eco-evolutionary process. Proc. R. Soc. B Biol. Sci., 278, 2401–2411.

Morris, D.W., Davidson, D.L. & Krebs, C.J. (2000). Measuring the ghost of competition: Insights from density-dependent habitat selection on the co-existence and dynamics of lemmings. Evol. Ecol. Res., 2, 41–67.

Mueller, T. & Fagan, W.F. (2008). Search and navigation in dynamic environments – from individual behaviors to population distributions. Oikos, 117, 654–664.

Nagelkerke, N.J.D. (1991). A note on a general definition of the coefficient of determination. Biometrika, 78, 691–692.

Northrup, J.M., Vander Wal, E., Bonar, M., Fieberg, J., Laforge, M.P., Leclerc, M., et al. (2022). Conceptual and methodological advances in habitat-selection modeling: guidelines for ecology and evolution. Ecol. Appl., 32, e02470.

Peacor, S.D. (2003). Phenotypic modifications to conspecific density arising from predation risk assessment. Oikos, 100, 409–415.

Peacor, S.D., Peckarsky, B.L., Trussell, G.C. & Vonesh, J.R. (2013). Costs of predator-induced phenotypic plasticity: A graphical model for predicting the contribution of nonconsumptive and consumptive effects of predators on prey. Oecologia, 171, 1–10.

Pérez-Barbería, F.J., Hooper, R.J. & Gordon, I.J. (2013). Long-term density-dependent changes in habitat selection in red deer (*Cervus elaphus*). Oecologia, 173, 837–847.

Phillips, E.M., Horne, J.K., Zamon, J.E., Felis, J.J. & Adams, J. (2019). Does perspective matter? A case study comparing Eulerian and Lagrangian estimates of common murre (*Uria aalge*) distributions. Ecol. Evol., 9, 4805–4819.

Proffitt, K.M., Gude, J.A., Shamhart, J. & King, F. (2012). Variations in elk aggregation patterns across a range of elk population sizes at Wall Creek, Montana. J. Wildl. Manage., 76, 847–856.

Robinson, N.P., Jones, M.O., Moreno, A., Erickson, T.A., Naugle, D.E. & Allred, B.W. (2019). Rangeland productivity partitioned to sub-pixel plant functional types. Remote Sens., 11, 1–9.

Rosenzweig, M.L. (1981). A Theory of Habitat Selection. Ecology, 62, 327–335.

Rosenzweig, M.L. (1991). Habitat Selection and Population Interactions: The Search for Mechanism. Am. Nat., 137, S5–S28.

Rosenzweig, M.L. & Abramsky, Z. (1985). Detecting Density-Dependent Habitat Selection. Am. Nat., 126, 405–417.

Rosenzweig, M.L. & Abramsky, Z. (1997). Two gerbils of the Negev: A long-term investigation of optimal habitat selection and its consequences. Evol. Ecol., 11, 733–756.

Ruth, T.K., Buotte, P.C. & Hornocker, M.G. (2019). Yellowstone cougars: ecology before and during wolf restoration. University Press of Colorado, Louisville, CO.

Sheriff, M.J., Peacor, S.D., Hawlena, D. & Thaker, M. (2020). Non-consumptive predator effects on prey population size: A dearth of evidence. J. Anim. Ecol., 89, 1302–1316.

Sinclair, A.R.E. & Arcese, P. (1995). Population Consequences of Predation-Sensitive Foraging. Ecology, 76, 882–891.

Smith, D.W., Cassidy, K.A., Stahler, D.R., MacNulty, D.R., Harrison, Q., Balmford, B., et al. (2020). Population Dynamics and Demography. In: Yellowstone Wolves. University of Chicago Press, pp. 77–92.

Stephens, P.A. & Sutherland, W.J. (1999). Consequences of the Allee effect for behaviour, ecology and conservation. Trends Ecol. Evol., 14, 401–405.

Tallian, A., Smith, D.W., Stahler, D.R., Metz, M.C., Wallen, R.L., Geremia, C., et al. (2017). Predator foraging response to a resurgent dangerous prey. Funct. Ecol., 31, 1418–1429.

Trussell, G.C., Ewanchuk, P.J. & Matassa, C.M. (2006). Habitat effects on the relative importance of trait- and density-mediated indirect interactions. Ecol. Lett., 9, 1245–1252.

Turchin, P. (1990). Rarity of density dependence or population regulation with lags? Nature, 344, 660– 663.

Turchin, P. (1998). Quantitative Analysis of Movement: Measuring and Modeling Population Redistribution in Animals and Plants. Sinauer Press, Sunderland, MA.

de Valpine, P., Paciorek, C., Turek, D., Michaud, N., Anderson-Bergman, C., Obermeyer, F., et al. (2021). NIMBLE: MCMC, Particle Filtering, and Programmable Hierarchical Modeling. R package version 0.11.1.

de Valpine, P., Turek, D., Paciorek, C., Anderson-Bergman, C., Temple Lang, D. & Bodik, R. (2017). Programming with models: writing statistical algorithms for general model structures with NIMBLE. J. Comput. Graph. Stat., 26, 403–413.

White, P.J., Proffitt, K.M. & Lemke, T.O. (2012). Changes in Elk Distribution and Group Sizes after Wolf Restoration. Am. Midl. Nat., 167, 174–187.

White, P.J., Proffitt, K.M., Mech, L.D., Evans, S.B., Cunningham, J.A. & Hamlin, K.L. (2010). Migration of northern Yellowstone elk: implications of spatial structuring. J. Mammal., 91, 827–837.

## Literature Cited

Anton, C.B. (2020). The demography and comparative ethology of top predators in a multi-carnivore system. PhD Dissertation. UC Santa Cruz.

van Beest, F.M., McLoughlin, P.D., Vander Wal, E. & Brook, R.K. (2014). Density-dependent habitat selection and partitioning between two sympatric ungulates. Oecologia, 175, 1155–1165.

Cusack, J.J., Kohl, M.T., Metz, M.C., Coulson, T., Stahler, D.R., Smith, D.W., et al. (2020). Weak spatiotemporal response of prey to predation risk in a freely interacting system. J. Anim. Ecol., 89, 120–131.

Fortin, D., Beyer, H.L., Boyce, M.S., Smith, D.W., Duchesne, T. & Mao, J.S. (2005). Wolves influence elk movements: Behavior shapes a trophic cascade in Yellowstone National Park. Ecology, 86, 1320– 1330.

Hijmans, R.J. (2022). raster: Geographic data analysis and modeling. R Packag. version 3.5-15.

Hooten, M.B., Johnson, D.S., McClintock, B.T. & Morales, J.M. (2017). Animal Movement. First. CRC Press, Boca Raton, FL.

Kohl, M.T., Stahler, D.R., Metz, M.C., Forester, J.D., Kauffman, M.J., Varley, N., et al. (2018). Diel predator activity drives a dynamic landscape of fear. Ecol. Monogr., 88, 638–652.

Manly, B.F.J., McDonald, L.L., Thomas, D.L., McDonald, T.L. & Erickson, W.P. (2002). Resource Selection by Animals. Springer Netherlands, Dordrecht, The Netherlands.

Mao, J.S., Boyce, M.S., Smith, D.W., Singer, F.J., Vales, D.J., Vore, J.M., et al. (2005). Habitat selection by elk before and after wolf reintroduction in Yellowstone National Park. J. Wildl. Manage., 69, 1691–1707.

Plummer, M., Best, N., Cowles, K. & Vines, K. (2006). CODA: Convergence Diagnosis and Output Analysis for MCMC. R News, 6, 7–11.

Samuel, M.D., Garton, E.O., Schlegel, M.W. & Carson, R.G. (1987). Visibility bias during aerial surveys of elk in northcentral Idaho. J. Wildl. Manage., 51, 622–630.

Singer, F.J. & Garton, E.O. (1994). Elk sightability model for the Super Cub. In: Aerial survey: user’s manual with practical tips for designing and conducting aerial big game surveys. pp. 47–48.

Thornton, M.M., Shrestha, R., Wei, Y., Thornton, P.E., Kao, S. & Wilson, B.E. (2020). Daymet: Daily Surface Weather Data on a 1-km Grid for North America, Version 4.

U.S. Geological Survey. (2020). 3D Elevation Program 1/3-arcsecond Digital Elevation Models (published 2020-11-13).

