## Appendix for "Density-dependent habitat selection alters drivers of population distribution in northern Yellowstone elk"

### Appendix S3 – Model Structure and Random Effects

As detailed in the main text, we modeled the expected count $\left(\lambda_{i,t}\right)$ in each pixel in each year as a function of 22 covariates (indexed by $k$; Table S1), a time-varying offset ($\alpha_t$), a temporal random effect ($\eta_{i,t}$), and a spatial random effect ($s_i$, eqn. 1). We described the fixed covariates in the main text, and we described the offset and random effects in detail here.

*Offset*

To account for changing total abundance across years, we included a time-varying offset, $\alpha_t$.The offset was the log of average range-wide density of elk across the study area in each year, i.e., the natural logarithm of the total count ($N_t$) divided by the number of pixels in the study area ($\Omega$), $\log\left(\frac{N_t}{\Omega}\right)$. If none of the covariates had any effect (i.e., for all $k$, $\beta_k = 0$), then the expected density in each pixel would simplify to $\lambda_{i,t} = \exp[\alpha_t] = \exp\left[\log\left(\frac{N_t}{\Omega}\right)\right] = N_t/\Omega$, which is simply the average range-wide density for that year. In that way, the offset acts like a time-varying intercept. Note that when we refer to "average range-wide density" or "log average range-wide density" throughout, we are referring to a predictor variable, such as this offset described here or the interaction terms described below. Contrast this with "expected density," which is the expectation of our response variable, $\lambda_{i,t}$.

*Temporal Random Effect*

We used the temporal random effect $\left(\eta_{i,t}\right)$ to account for temporal fluctuations in the effect of a raster pixel being outside YNP compared to inside YNP. The two main motivations for this were (1) an observed shift in the elk distribution from primarily inside YNP in the early years of our study to primarily outside the northern boundary of YNP in later years (Tallian et a. 2017; MacNulty et al. 2020), and (2) a late hunt managed by the state of Montana that occurred in January of each year prior to 2010 but ceased afterward (MacNulty et al. 2020). We estimated the effect in each year ($t$) as coming from a normal distribution with mean $\mu_\eta$ and variance $\sigma_\eta^2$. We multiplied the effect by a binary covariate indicating whether each pixel was outside of YNP, with outside = 1 and inside = 0.

The temporal random effect $\left(\eta_{i,t}\right)$ had a mean of -1.74 (90% credible interval = -2.05 – -1.42), and a variance of 0.56 (90% credible interval = 0.24 – 1.09). The posterior mean effect for each year ranged -2.77 – -0.39, and the trend was positive over time (Fig. S12). In this context, negative values indicate lower elk densities outside YNP than would be expected based on the fixed effects in the model and the positive trend indicates increasing densities outside YNP over time.

*Spatial Random Effect*

We used the spatial random effect ($s_i \in \boldsymbol{s}$) to account for residual spatial autocorrelation in elk density, i.e., consistent spatial pattern in elk density not captured by the other terms in the model. We modeled it as a Gaussian process (a.k.a., kriging), i.e., the vector $\boldsymbol{s}$ is a realization of a multivariate normal distribution with mean 0 and variance-covariance matrix, $\boldsymbol{\Sigma}$ ($\boldsymbol{s} \sim N(\boldsymbol{0}, \boldsymbol{\Sigma})$). Each element of the variance-covariance matrix was calculated as a decaying exponential function of the distance between the two pixels, $\left\|x_i - x_j\right\|$, and two free parameters, $\sigma$ and $\rho$, estimated by the model: $\Sigma_{i,j} = \sigma^2 \times \exp\left(-\frac{\|x_i - x_j\|}{\rho}\right)$. The parameter $\sigma$ controlled the variance within a pixel (i.e., the distance between a pixel and itself is 0, thus $\Sigma_{i,i} = \sigma^2$). The parameter $\rho$ controlled the rate of decay with distance, with smaller $\rho$ indicating faster decay. To improve model fitting, we scaled distances between pixels to range from 0 – 1 by dividing by the maximum distance.

### Appendix S8 – Supplemental Tables

#### *Table S1. Model covariates*

**Table S1.** Covariates included to model expected density $(\lambda_{i,t})$. Colons (:) in the "Name" column indicate interactions. "Rationale" describes the reason we included a parameter in the model. The first four variables (k = 1 – 4) measure fixed conditions that are known to affect elk density but were not the main target of inference. Variables including biomass (k = 5 – 6) are the food variables and variables including openness or roughness (k = 7 – 22) are the safety variables. DDHS = density-dependent habitat selection.

| Index $(k)$ | Name | Description | Rationale |
|---|---|---|---|
| 1 | SWE | Snow-water equivalent | Control for autecological conditions. |
| 2 | Elev | Elevation | |
| 3 | cos(Asp) | Cosine of aspect, N-S component | |
| 4 | sin(Asp) | Sine of aspect, E-W component | |
| 5 | Biomass | Above ground biomass of grasses and forbs from previous growing season (log-transformed) | Measure DDHS for food. |
| 6 | Biomass:Dens | Interaction between biomass and log(density) | |
| 7 | Open | Openness (1 - % tree) | Measure DDHS for safety. |
| 8 | Rough | Roughness | |
| 9 | $Open^2$ | Open-squared | |
| 10 | $Rough^2$ | Rough-squared | |
| 11 | Open:Dens | Interaction between safety and log(density) | |
| 12 | Rough:Dens | | |
| 13 | $Open^2$:Dens | | |
| 14 | $Rough^2$:Dens | | |
| 15 | Open:Wolf | Interaction between safety and wolf density | Measure change in perceived safety with predator density; corroborates safety variables as biologically meaningful measures of predation risk. |
| 16 | Rough:Wolf | | |
| 17 | $Open^2$:Wolf | | |
| 18 | $Rough^2$:Wolf | | |
| 19 | Open:Cougar | Interaction between safety and cougar density | |
| 20 | Rough:Cougar | | |
| 21 | $Open^2$:Cougar | | |
| 22 | $Rough^2$:Cougar | | |

*Table S2. ANOVA for residuals by month*

**Table S2.** ANOVA for model residuals by month. We examined whether the date of the survey affected model residuals. “Day-of-year Comparison” indicates the two compared dates, and “Residual Difference” indicates the difference in mean residuals between the two dates. The p-value was adjusted using Tukey’s Honest Significant Difference test. All p-values are > 0.23, showing that residuals do not differ between any pairs of dates. These same data are visualized in Fig. S3.

| Day-of-year Comparison | Residual Difference | p-value (adjusted) |
|---|---|---|
| 2-1 | -1.44 | 0.909 |
| 5-1 | -1.69 | 0.743 |
| 15-1 | -1.98 | 0.239 |
| 18-1 | -0.77 | 1.000 |
| 20-1 | -1.46 | 0.900 |
| 30-1 | -2.26 | 0.252 |
| 45-1 | -0.90 | 0.999 |
| 49-1 | -0.44 | 1.000 |
| 57-1 | -2.19 | 0.304 |
| 67-1 | -1.87 | 0.589 |
| 350-1 | -1.03 | 0.995 |
| 355-1 | -0.71 | 1.000 |
| 358-1 | -0.06 | 1.000 |
| 364-1 | -0.94 | 0.998 |
| 5-2 | -0.26 | 1.000 |
| 15-2 | -0.54 | 1.000 |
| 18-2 | 0.67 | 1.000 |
| 20-2 | -0.02 | 1.000 |
| 30-2 | -0.82 | 1.000 |
| 45-2 | 0.54 | 1.000 |
| 49-2 | 1.00 | 0.997 |
| 57-2 | -0.75 | 1.000 |
| 67-2 | -0.43 | 1.000 |
| 350-2 | 0.41 | 1.000 |
| 355-2 | 0.73 | 1.000 |
| 358-2 | 1.38 | 0.933 |
| 364-2 | 0.50 | 1.000 |
| 15-5 | -0.28 | 1.000 |
| 18-5 | 0.92 | 0.998 |
| 20-5 | 0.24 | 1.000 |
| 30-5 | -0.57 | 1.000 |
| 45-5 | 0.80 | 1.000 |
| 49-5 | 1.25 | 0.970 |
| 57-5 | -0.50 | 1.000 |
| 67-5 | -0.17 | 1.000 |
| 350-5 | 0.67 | 1.000 |
| 355-5 | 0.99 | 0.997 |
| 358-5 | 1.64 | 0.788 |
| 364-5 | 0.75 | 1.000 |
| 18-15 | 1.21 | 0.928 |
| 20-15 | 0.52 | 1.000 |
| 30-15 | -0.29 | 1.000 |
| 45-15 | 1.08 | 0.971 |
| 49-15 | 1.53 | 0.677 |
| 57-15 | -0.21 | 1.000 |
| 67-15 | 0.11 | 1.000 |
| 350-15 | 0.95 | 0.991 |
| 355-15 | 1.27 | 0.897 |
| 358-15 | 1.92 | 0.285 |
| 364-15 | 1.04 | 0.980 |
| 20-18 | -0.69 | 1.000 |
| 30-18 | -1.49 | 0.881 |
| 45-18 | -0.13 | 1.000 |
| 49-18 | 0.33 | 1.000 |
| 57-18 | -1.42 | 0.917 |
| 67-18 | -1.10 | 0.991 |
| 350-18 | -0.26 | 1.000 |
| 355-18 | 0.06 | 1.000 |

| | | |
|---|---|---|
| 358-18 | 0.71 | 1.000 |
| 364-18 | -0.17 | 1.000 |
| 30-20 | -0.81 | 1.000 |
| 45-20 | 0.56 | 1.000 |
| 49-20 | 1.01 | 0.996 |
| 57-20 | -0.73 | 1.000 |
| 67-20 | -0.41 | 1.000 |
| 350-20 | 0.43 | 1.000 |
| 355-20 | 0.75 | 1.000 |
| 358-20 | 1.40 | 0.926 |
| 364-20 | 0.52 | 1.000 |
| 45-30 | 1.37 | 0.938 |
| 49-30 | 1.82 | 0.632 |
| 57-30 | 0.07 | 1.000 |
| 67-30 | 0.40 | 1.000 |
| 350-30 | 1.24 | 0.973 |
| 355-30 | 1.55 | 0.845 |
| 358-30 | 2.21 | 0.292 |
| 364-30 | 1.32 | 0.952 |
| 49-45 | 0.46 | 1.000 |
| 57-45 | -1.29 | 0.960 |
| 67-45 | -0.97 | 0.997 |
| 350-45 | -0.13 | 1.000 |
| 355-45 | 0.19 | 1.000 |

| | | |
|---|---|---|
| 358-45 | 0.84 | 0.999 |
| 364-45 | -0.04 | 1.000 |
| 57-49 | -1.75 | 0.697 |
| 67-49 | -1.42 | 0.915 |
| 350-49 | -0.59 | 1.000 |
| 355-49 | -0.27 | 1.000 |
| 358-49 | 0.39 | 1.000 |
| 364-49 | -0.50 | 1.000 |
| 67-57 | 0.32 | 1.000 |
| 350-57 | 1.16 | 0.984 |
| 355-57 | 1.48 | 0.887 |
| 358-57 | 2.13 | 0.349 |
| 364-57 | 1.25 | 0.970 |
| 350-67 | 0.84 | 0.999 |
| 355-67 | 1.16 | 0.985 |
| 358-67 | 1.81 | 0.642 |
| 364-67 | 0.93 | 0.998 |
| 355-350 | 0.32 | 1.000 |
| 358-350 | 0.97 | 0.997 |
| 364-350 | 0.09 | 1.000 |
| 358-355 | 0.65 | 1.000 |
| 364-355 | -0.23 | 1.000 |
| 364-358 | -0.88 | 0.999 |

*Table S3. Gelman-Rubin Statistics*

**Table S3.** Gelman-Rubin statistics for MCMC convergence diagnosis. Values for potential scale reduction factors (PSRFs) close to 1 indicate model convergence. All PSRFs for top-level parameters were estimated to be < 1.02, indicating good convergence. Parameter names match the NIMBLE code, which is available on GitHub and archived on Zenodo (doi:10.5281/zenodo.6687905). The column "Point est." gives the estimated PSRF, and the column "Upper C.I." gives the upper bound on the PSRF.

| Parameter | Point est. | Upper C.I. |
|---|---|---|
| beta[1] | 1.00 | 1.00 |
| beta[2] | 1.01 | 1.02 |
| beta[3] | 1.00 | 1.01 |
| beta[4] | 1.00 | 1.02 |
| beta[5] | 1.00 | 1.00 |
| beta[6] | 1.00 | 1.00 |
| beta[7] | 1.00 | 1.00 |
| beta[8] | 1.00 | 1.00 |
| beta[9] | 1.00 | 1.00 |
| beta[10] | 1.00 | 1.00 |
| beta[11] | 1.00 | 1.00 |
| beta[12] | 1.00 | 1.00 |
| beta[13] | 1.00 | 1.00 |
| beta[14] | 1.00 | 1.00 |
| beta[15] | 1.00 | 1.00 |
| beta[16] | 1.00 | 1.00 |
| beta[17] | 1.00 | 1.00 |
| beta[18] | 1.00 | 1.00 |
| beta[19] | 1.00 | 1.01 |
| beta[20] | 1.00 | 1.01 |
| beta[21] | 1.00 | 1.00 |
| beta[22] | 1.00 | 1.00 |
| eta_out[1] | 1.00 | 1.00 |
| eta_out[2] | 1.00 | 1.00 |
| eta_out[3] | 1.00 | 1.00 |
| eta_out[4] | 1.00 | 1.00 |
| eta_out[5] | 1.00 | 1.01 |
| eta_out[6] | 1.01 | 1.02 |
| eta_out[7] | 1.00 | 1.01 |
| eta_out[8] | 1.00 | 1.00 |
| eta_out[9] | 1.00 | 1.00 |
| eta_out[10] | 1.00 | 1.00 |
| eta_out[11] | 1.00 | 1.00 |
| eta_out[12] | 1.00 | 1.00 |
| eta_out[13] | 1.00 | 1.01 |
| eta_out[14] | 1.00 | 1.00 |
| eta_out[15] | 1.00 | 1.01 |
| eta_out[16] | 1.00 | 1.00 |
| mu_out | 1.00 | 1.01 |
| nb_r | 1.00 | 1.00 |
| rho | 1.00 | 1.01 |
| sigma | 1.02 | 1.06 |
| sigma_out | 1.00 | 1.00 |

#### Appendix S9 – Supplemental Figures

*Figure S1. Study area map*

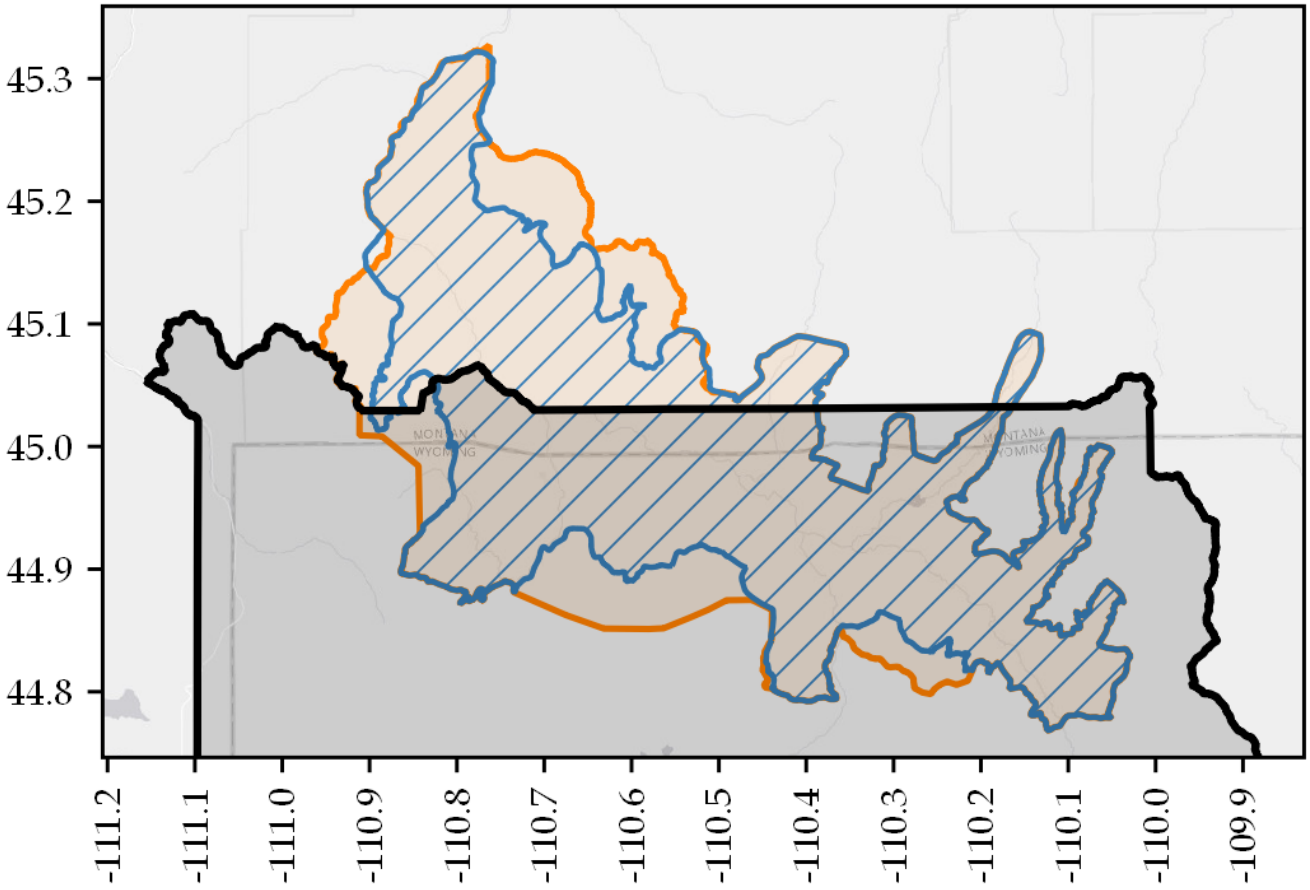


**Figure S1.** Study area showing historic northern Yellowstone elk winter range polygon (blue; 1,520 km$^2$), and our adjusted winter range polygon (orange; 1,900 km$^2$), and the boundary of Yellowstone National Park (black).

*Figure S2. Survey dates and times*

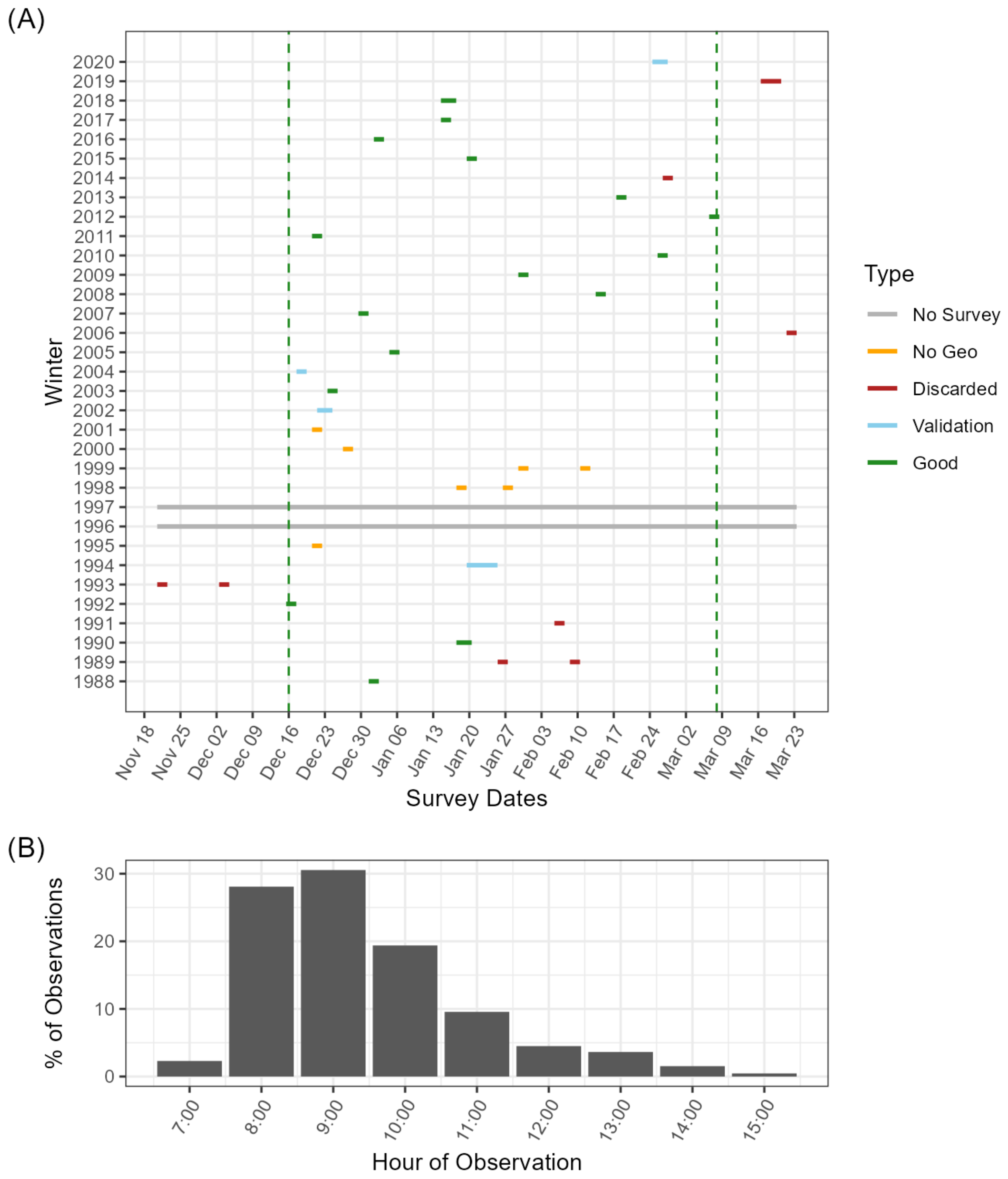


**Figure S2.** We fit our model using data collected from 1988 to 2020. Locations of elk groups (≥ 1 elk) and their sizes were recorded during aerial surveys from fixed-wing aircraft. (A) Surveys used in the model were conducted between December and March in each year. In 1996 and 1997, no survey occurred (gray horizontal lines). Counts for 1989, 1991, 2006, and 2014 are considered unreliable as a census of the population and so were discarded (red lines). The count for 2019 was conducted via helicopter late in the season and was also discarded (red line). Georeferenced group data are unavailable between 1995 and 2001 and could not be included in the model (orange). In 1994, 2002, and 2004, georeferenced group data are only available inside of Yellowstone National Park (data from the State of Montana were not georeferenced); these data were not included for model fitting but were included for model validation (blue lines). A full census was available for 2020, but we chose to withhold it as an additional validation year with complete georeferencing (blue line). All other years were used for model fitting (green lines). All surveys used in modeling fall between December 16 and March 7 (vertical green dashed lines). (B) Surveys were conducted primarily in the morning hours, and 92% of group observations were recorded between 08:00 and 12:00.

*Figure S3. Residuals by survey month*

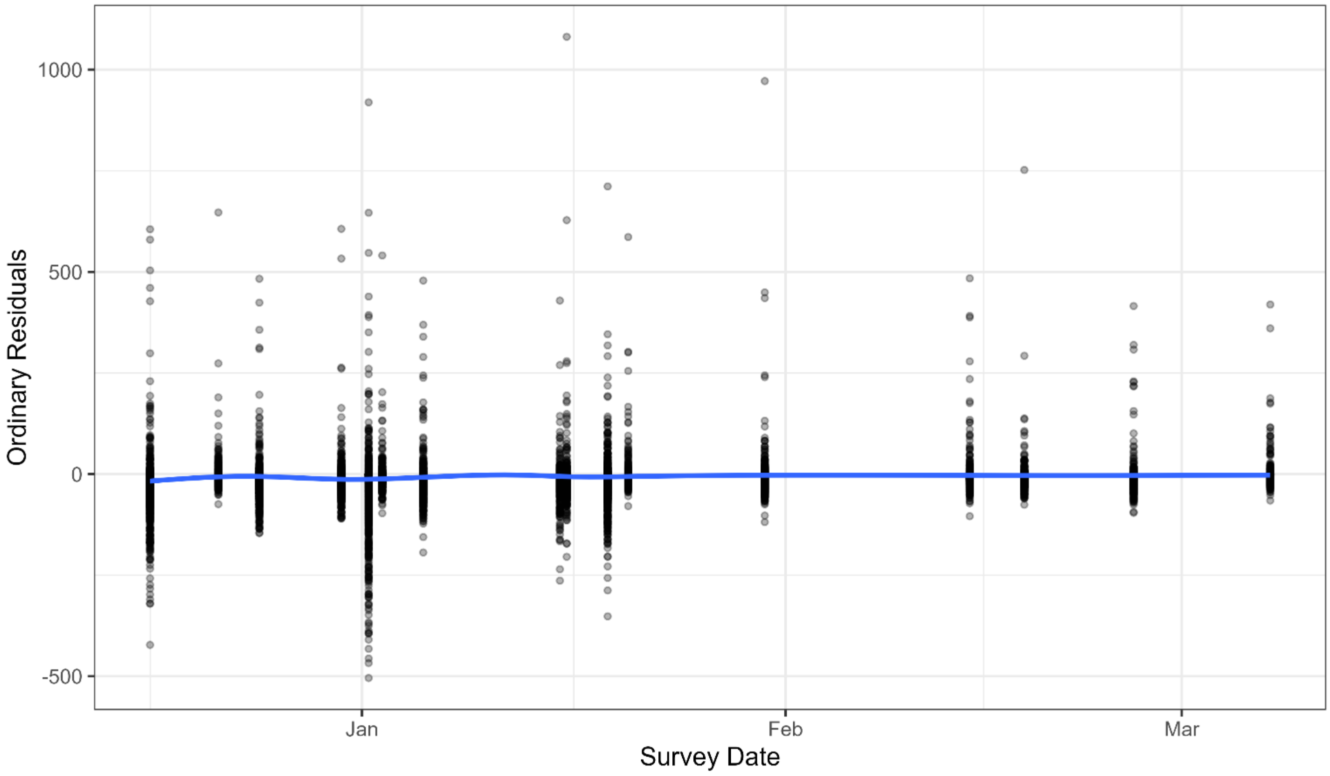


**Figure S3.** Ordinary model residuals by survey month. We checked that variable survey dates (Fig. S2A) did not introduce a bias in our model. We plotted the model's ordinary residuals as a function of survey date, and visually checked for any pattern by fitting a very flexible GAM smoothing line, which could have picked up subtle non-linear differences by date (blue line). The fitted line shows no obvious pattern, which was confirmed by an ANOVA and Tukey's HSD test (Table S2).

*Figure S4. Scale of analysis*

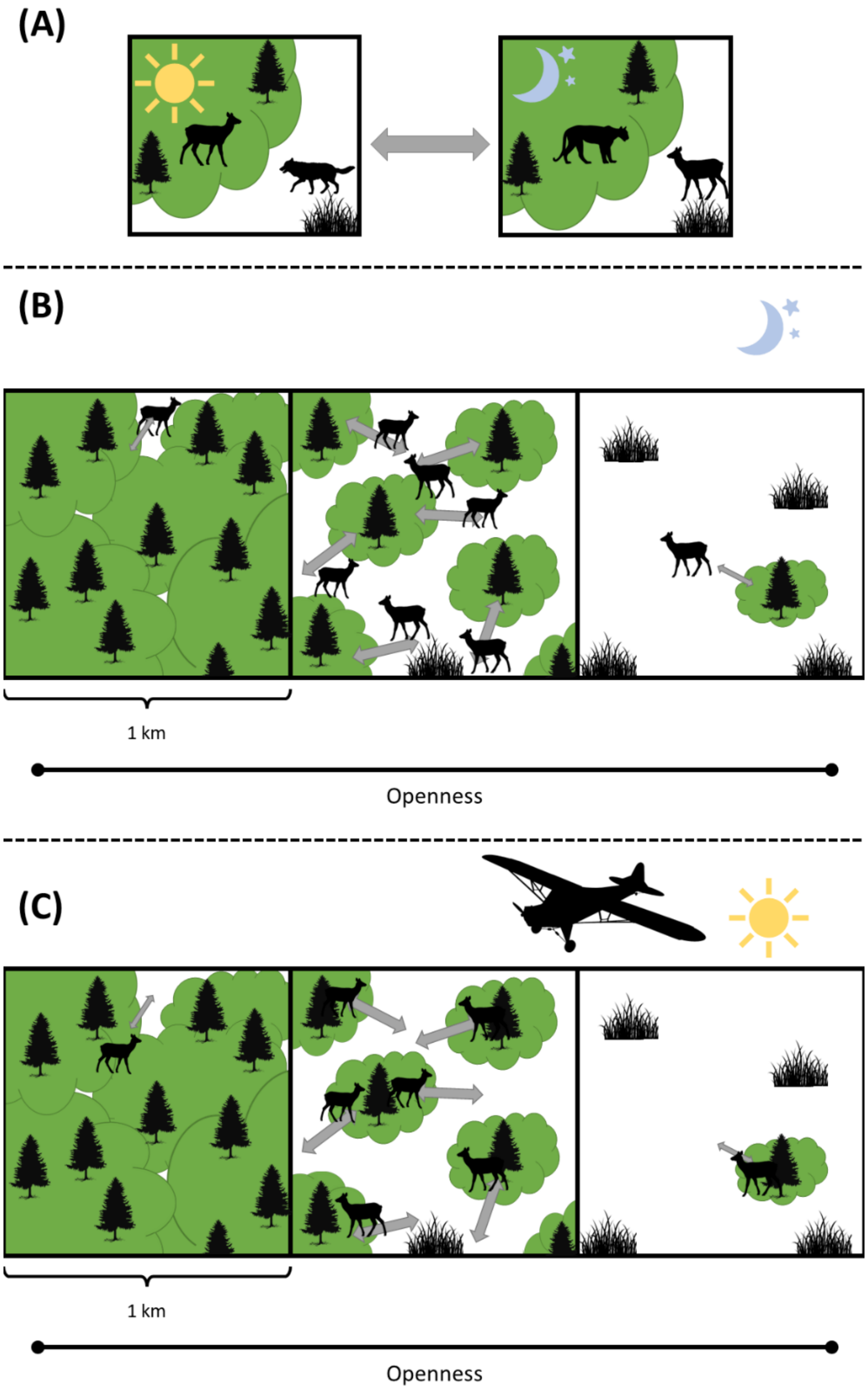


**Figure S4**. Spatial scale of analysis. We rasterized all counts and covariates on the same 1 km x 1 km raster grid. We chose this coarse spatial scale to average over daily elk movements and capture the population-level distribution. Elk are known to move between habitats characterized by openness (depicted here) and roughness (not depicted) at different times of day to manage their risk from both wolves and cougars. In terms of openness, risk from wolves is greatest in habitats with little or no tree cover, whereas risk from cougars is greatest in forested habitats. (A) Elk spend daytime hours, when wolves are more active, in forested habitats (left panel), whereas they spend nighttime hours, when cougars are more active, in open habitats (right panel). (B and C) In one-km pixels across a gradient of varying openness, elk are limited by the heterogeneity of the pixel, with gray bidirectional arrows representing diel shuttling. The gray arrows are identical between (B) and (C), but (B) shows the nighttime position of elk and (C) shows the daytime position of elk. At the 1-km scale, we hypothesized that elk density would be greatest in habitats with intermediate openness (middle panel of B and C), which provide multiple patches that facilitate this diel shuttling behavior. Aerial surveys occur during daytime (C).

*Figure S5. Openness sensitivity analysis*

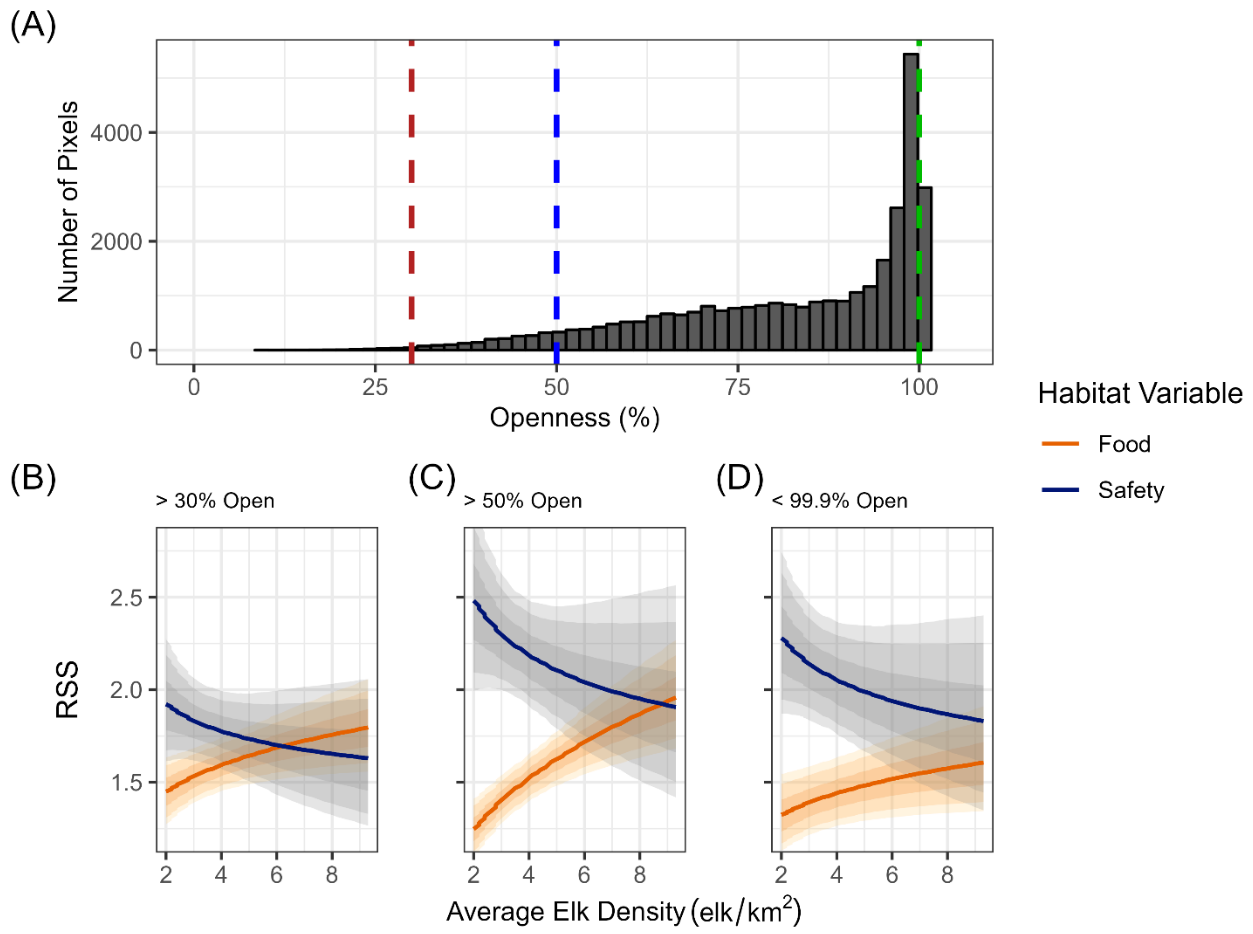


**Figure S5.** We conducted a sensitivity analysis to assess the potential impact of imperfect detection of elk due to low openness (high forest canopy cover) on our results. We discarded any pixels from the landscape that fell outside a particular openness cutoff in any year and refitted the model. (A) We defined three openness cutoffs, where the pixels retained for model fitting were >30% open (95% of original data retained; red dashed line), >50% open (70% of original data retained; blue dashed line), and <99.9% open (73% of original data retained; green dashed line). (B) Results for the >30% cutoff, (C) >50% cutoff, and (D) <99.9% cutoff were qualitatively similar to the results presented in the main text: we found positive DDHS for food, negative DDHS for safety, and a switch in their relative importance. DDHS for food equals the RSS for a 1-SD change in herbaceous biomass, and DDHS for safety equals a 0.5-SD change in openness and a 0.5-SD change in roughness. See main text for details. In panels (B) – (D), solid lines are posterior mean estimates and shaded envelopes are 50%, 80%, and 90% credible intervals.

*Figure S6. Mean effect of conditions*

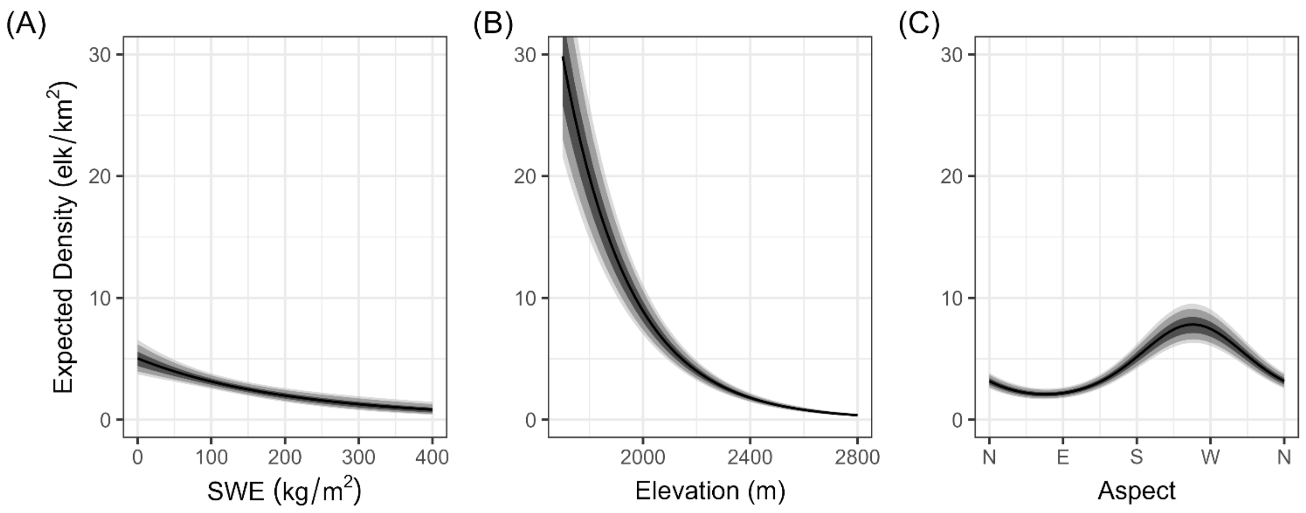


**Figure S6.** Mean effect of snow-water equivalent (SWE), elevation, and aspect on expected elk density during winter (December-February), with all other variables held at their mean. (A) Elk density was greatest in pixels with low snow, (B) low elevation, and (C) southwest aspects. Solid black lines show the mean effects, and the shaded gray envelopes show the 50%, 80%, and 90% credible intervals.

*Figure S7. Mean effect of food and safety variables*

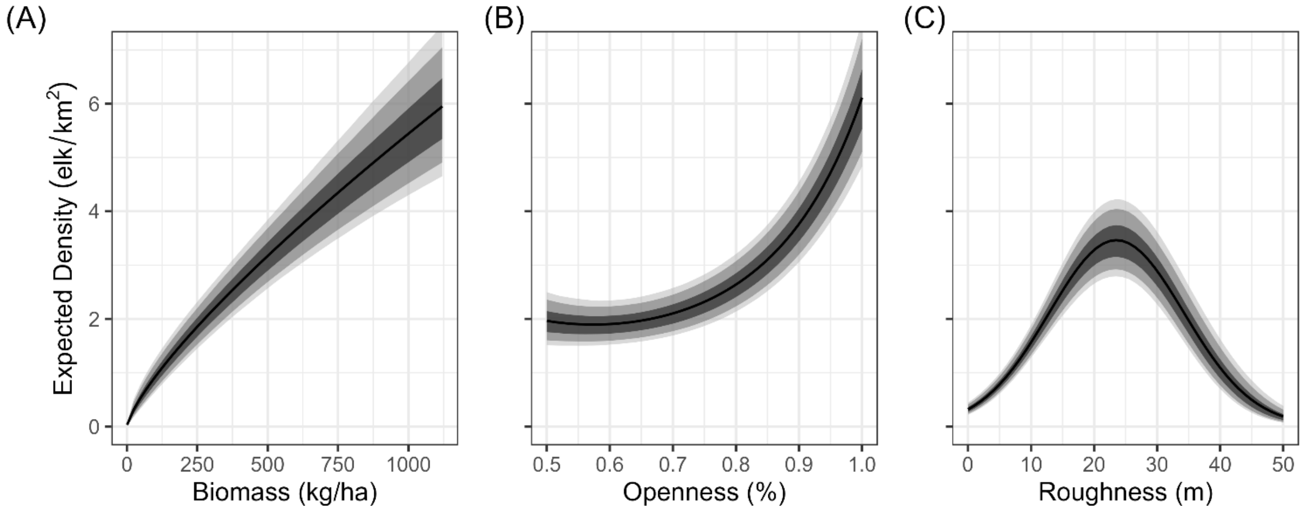


**Figure S7.** Mean effects of food and safety variables on expected elk density, with all other variables held at their overall mean. (A) Expected elk density increased with herbaceous biomass, consistent with our assertion that it is an important food resource for elk. (B) Contrary to our prediction that elk would most prefer an intermediate openness, expected elk density was greatest at 100% openness, and the relationship between openness and elk density was largely monotonic for the observed range of openness. (C) Expected elk density was greatest for a roughness of 23.5 m (intermediate, as expected). Solid black lines show the mean effects, and the shaded gray envelopes show the 50%, 80%, and 90% credible intervals.

*Figure S8. Effect of predator density on RSS for roughness*

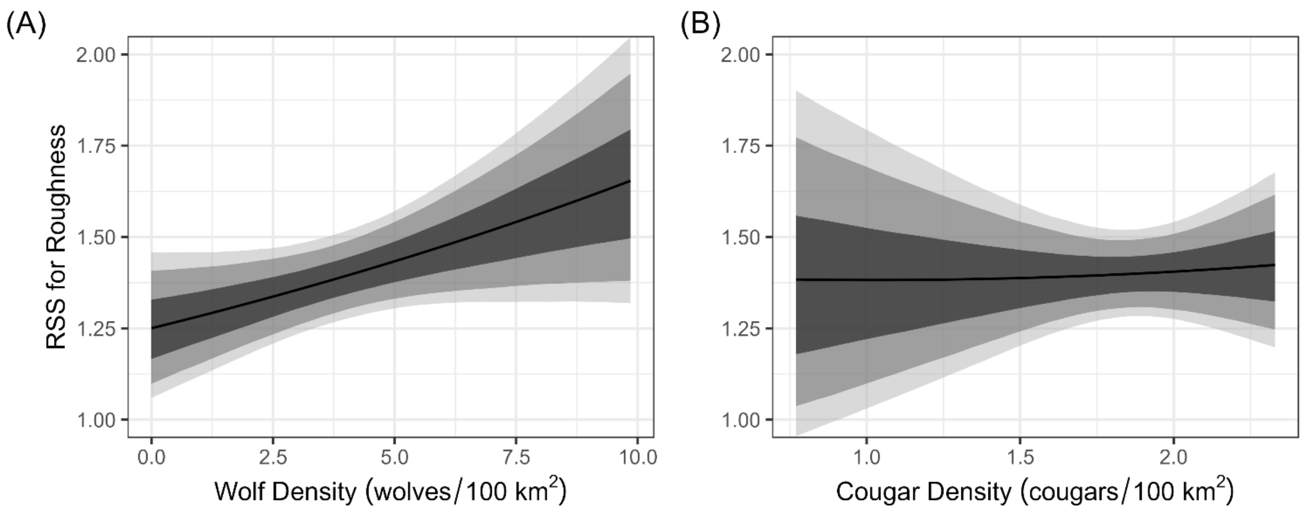


**Figure S8.** Mean effect of predator density on relative selection strength (RSS) for roughness, with all other variables held at their mean. (A) RSS for roughness increased with wolf density, in agreement with our prediction that elk would move away from the wolf habitat domain. (B) RSS for roughness did not change with cougar density, in agreement with our prediction that the wolf effect would be stronger than the cougar effect during the daytime aerial elk surveys. Solid black lines are mean effects, and shaded gray envelopes are 50%, 80%, and 90% credible intervals.

*Figure S9. DDHS for roughness*

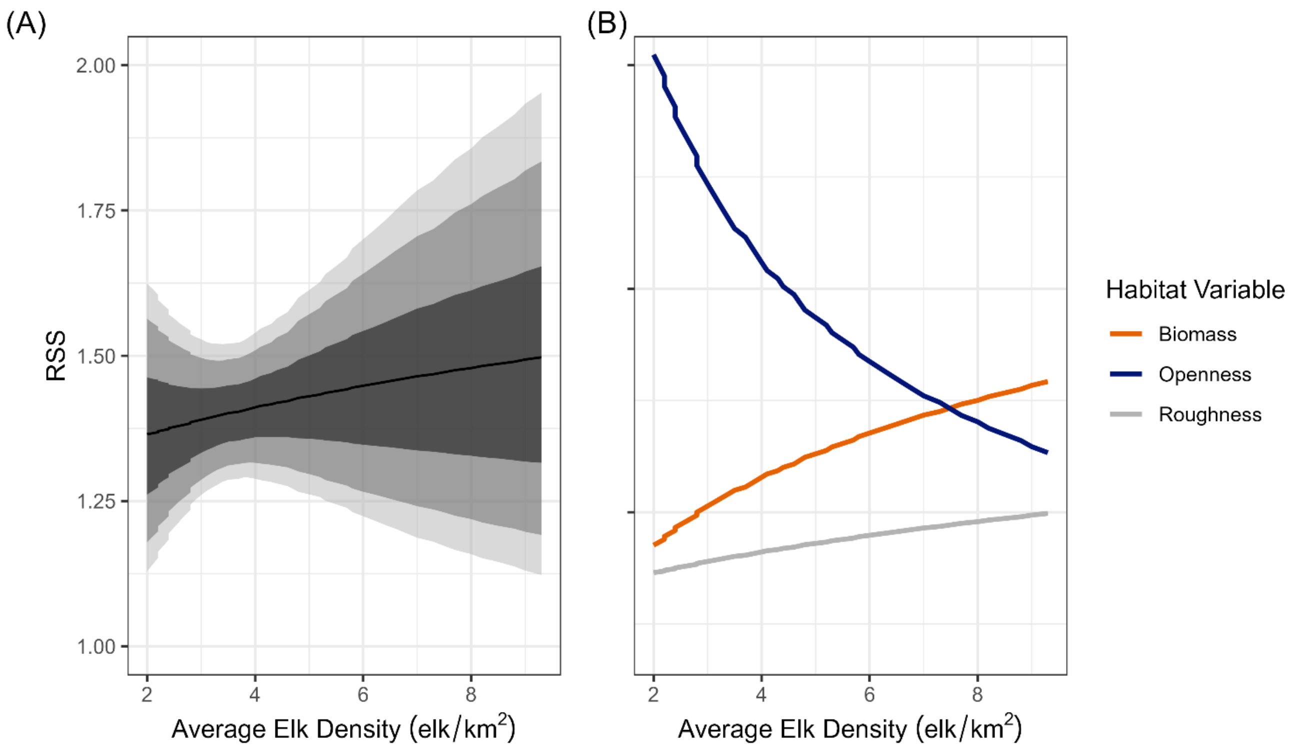


**Figure S9.** Density-dependent habitat selection (DDHS) for food and safety variables decomposed. (A) Average elk density versus relative selection strength (RSS) for a 1-SD change in roughness. The mean effect is weakly positive, but the large uncertainty indicates no support for DDHS for roughness. RSS was calculated using samples from the entire posterior distribution. Solid black line is the mean effect, and the shaded gray envelopes are the 50%, 80%, and 90% credible intervals. (B) Comparison of RSS for a 1-SD change in all food and safety variables.

*Figure S10. Model evaluation*

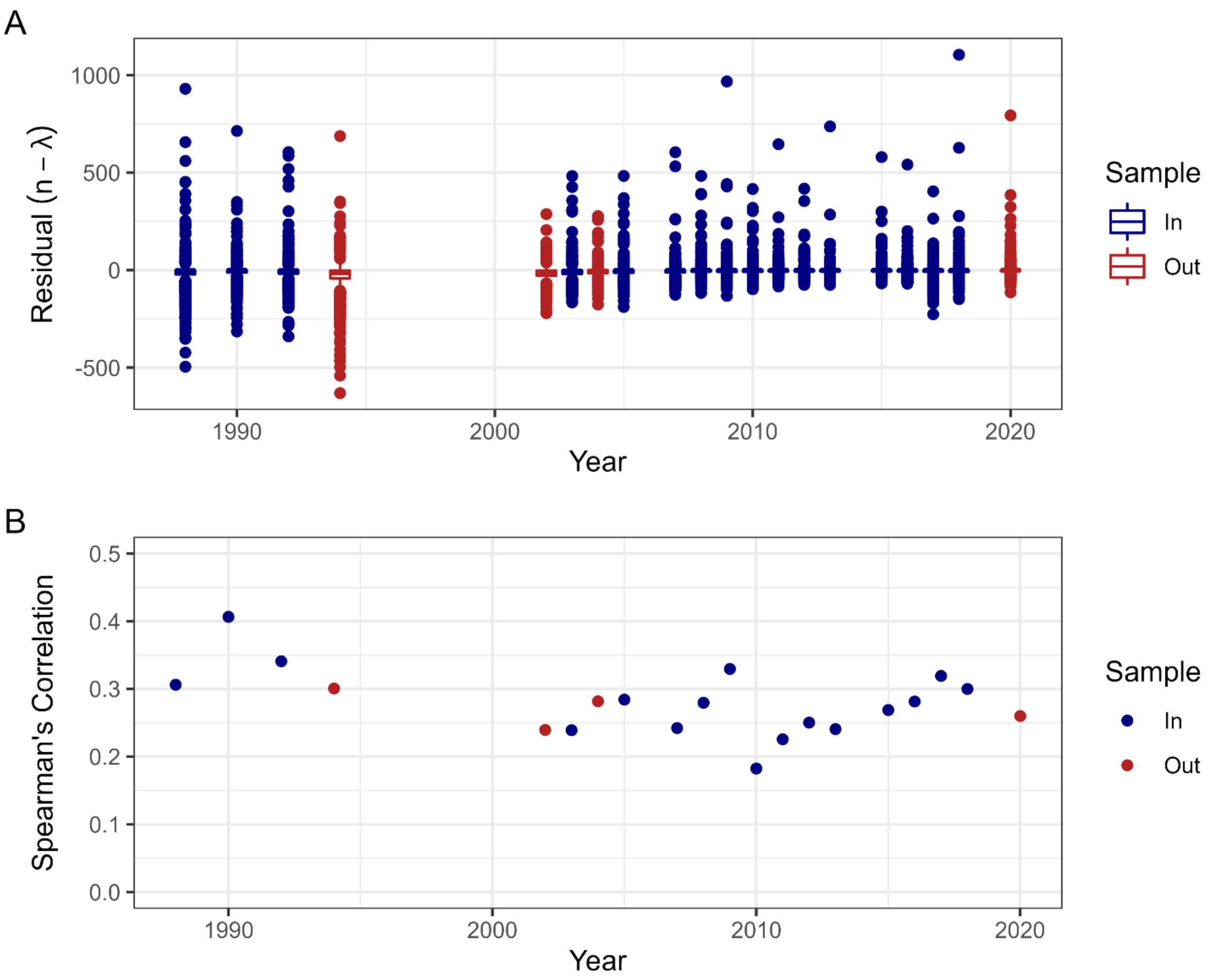


**Figure S10.** Model evaluation. We compared (A) ordinary residuals and (B) Spearman's correlation between expected and observed densities for both in-sample data (blue) and out-of-sample validation data (red). Residuals have mean near 0 in all years, showing good accuracy, but a wide spread, showing low precision. Spearman's correlation is moderate in all years (mean in-sample = 0.28, mean out-of-sample = 0.27), which shows the model has moderate power to rank pixels by abundance in both training and testing data.

*Figure S11. Residual spatial autocorrelation*

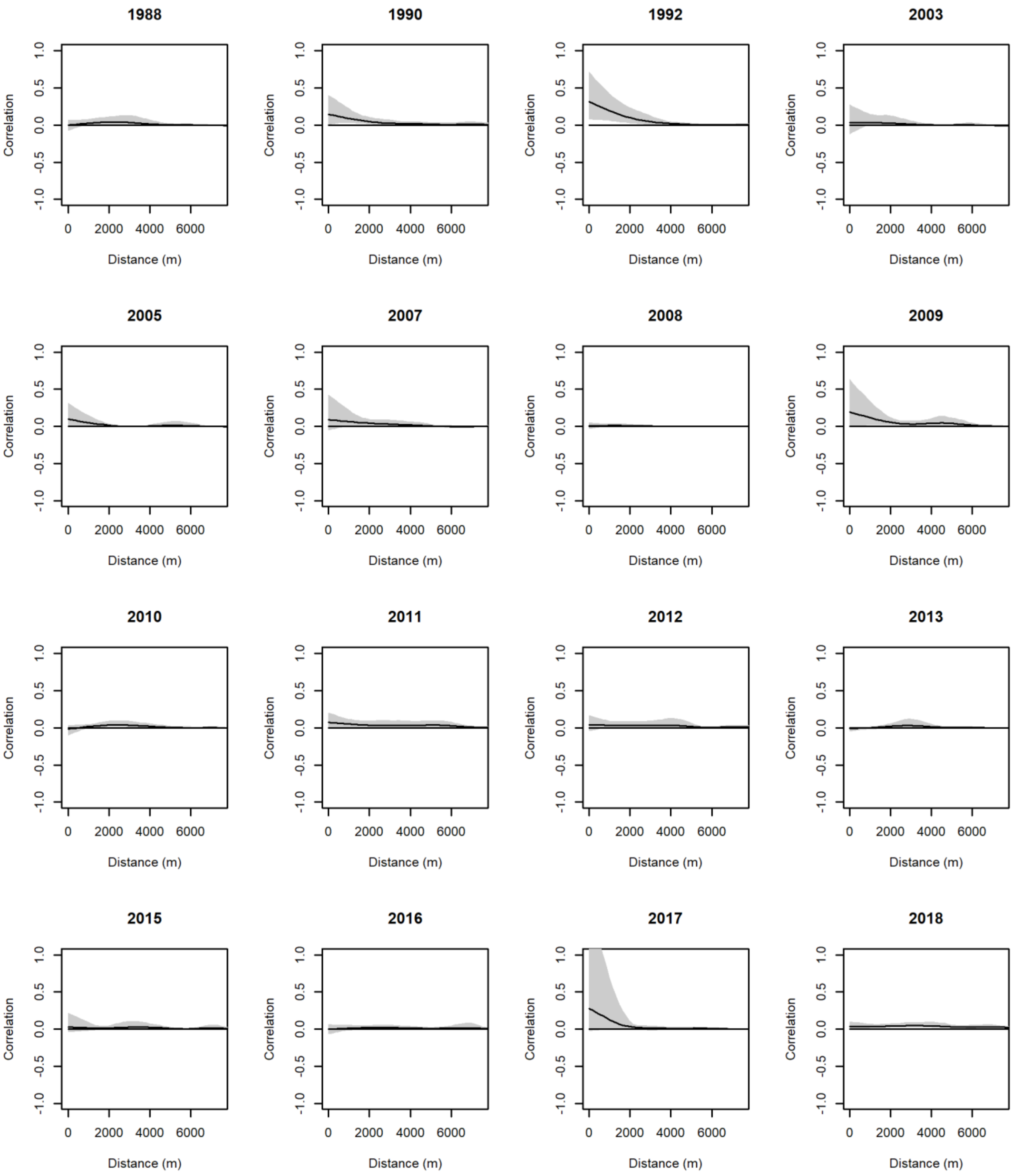


**Figure S11.** Non-parametric spline correlograms used to assess residual spatial autocorrelation. We estimated correlograms using Pearson's residuals for each pixel in each year. The scale of spatial autocorrelation is estimated as the distance where the confidence envelope first intersects the x-axis (correlation = 0). There was little to no residual spatial autocorrelation in all years, i.e., the confidence envelope crosses the x-axis near 0 m in nearly all years (1992 was the lone exception).

*Figure S12. Temporal random effect posterior estimates*

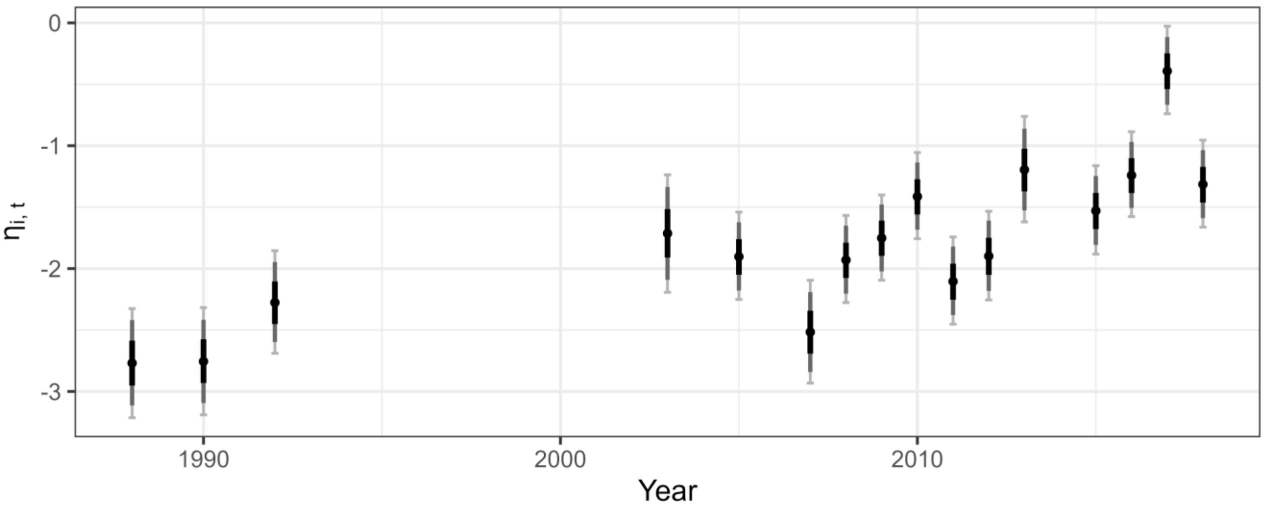


**Figure S12.** Temporal random effect $(\eta_{i,t})$ over time. We included the temporal random effect to account for changing patterns of elk distribution inside versus outside Yellowstone National Park (YNP), along with a change in the late hunt outside of YNP after 2010. The temporal random effect affects pixels outside of YNP, so negative values indicate that pixels outside of YNP have lower expected elk density than would be predicted based on fixed effects alone. Points show posterior mean for each coefficient and bars show credible intervals. Black bars show 50% credible intervals, dark gray bars show 80% credible intervals, and light gray bars with end caps show 90% credible intervals.

*Figure S13. Spatial random effect posterior mean*

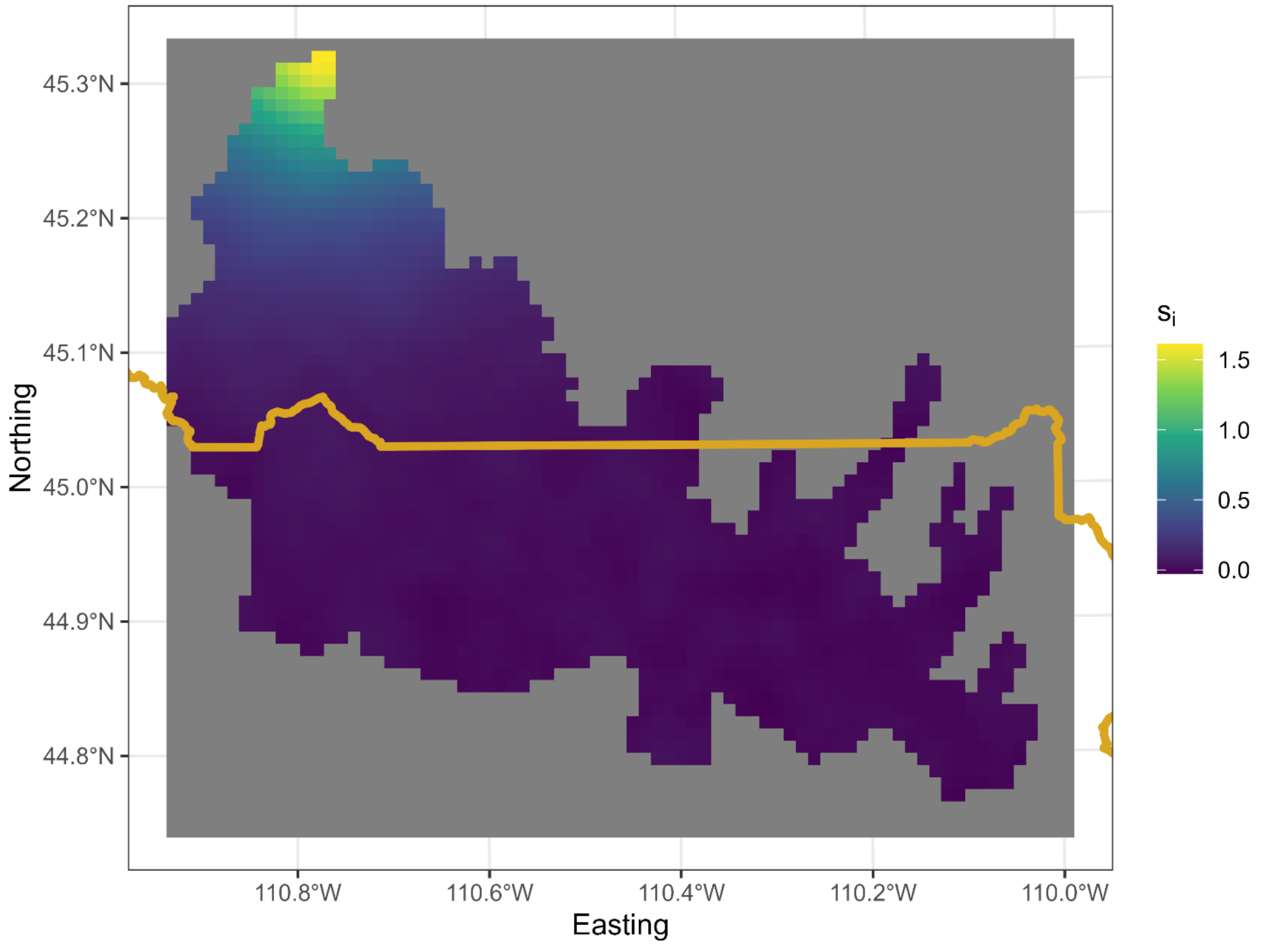


**Figure S13.** Posterior mean of estimated spatial random effect, ***s***.

*Figure S14. Fitted spatial covariance function*

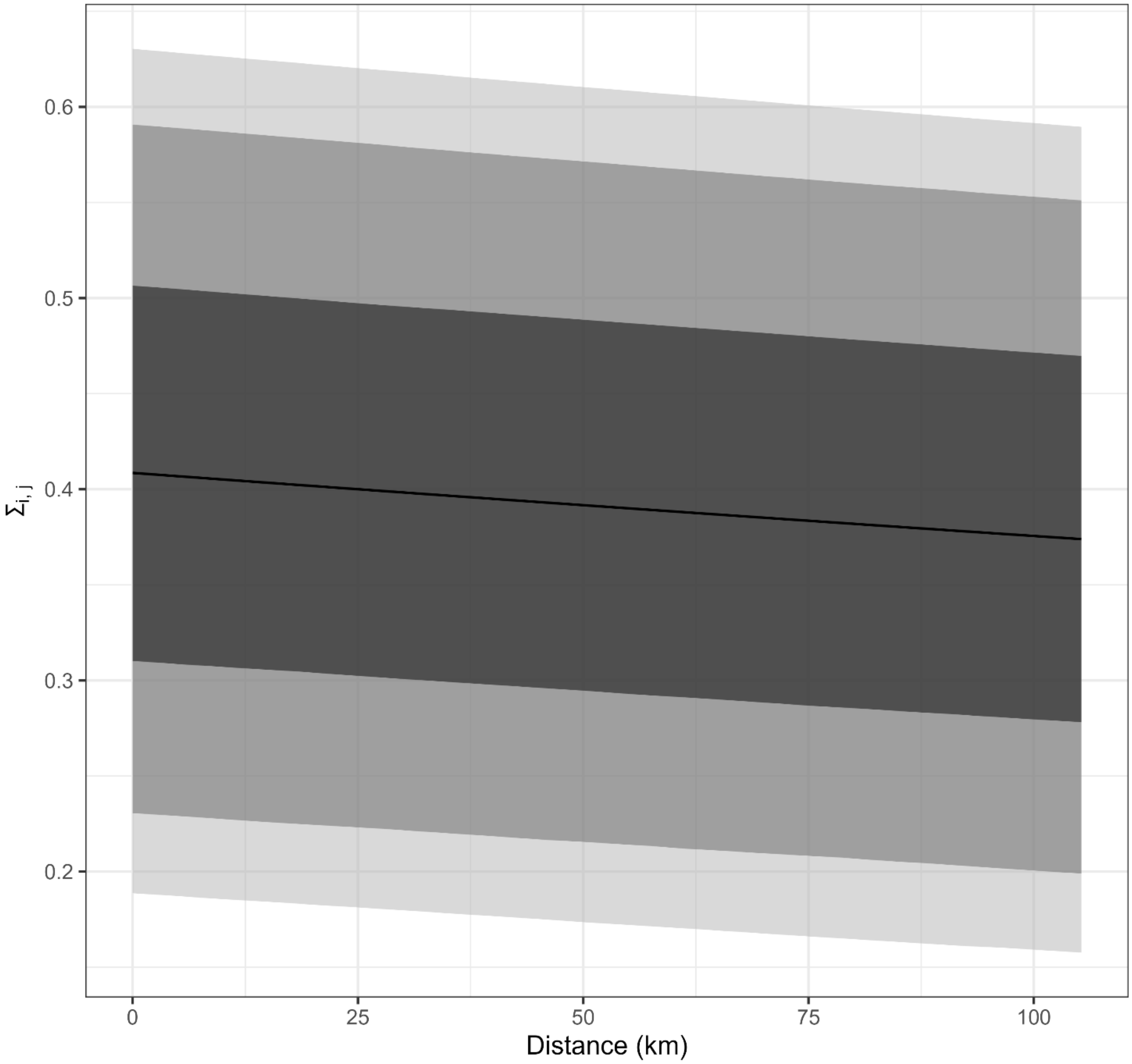


**Figure S14.** Fitted spatial covariance as a function of distance for the spatial random effect.

*Figure S15. Combined temporal and spatial random effects.*

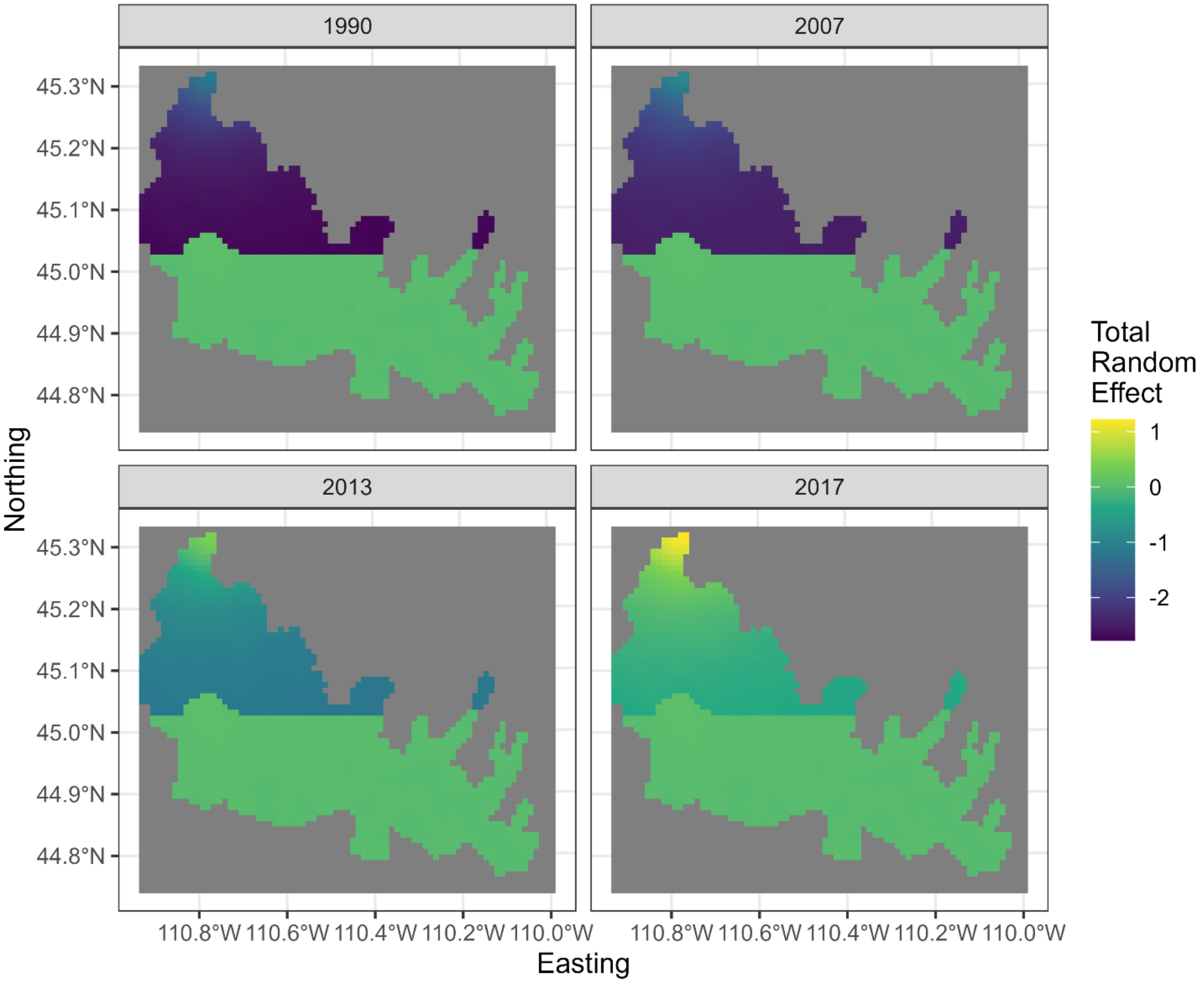


**Figure S15**. The temporal random effect was defined as operating outside Yellowstone National Park (YNP) and the estimated values were always negative. The spatial random effect could only take on positive values, but it could operate anywhere in the study area. The estimated values were largest outside of YNP. The negative effect of the temporal random effect and the positive effect of the spatial random effect thus acted in opposite directions and almost entirely outside YNP, with expected elk density inside YNP almost completely captured by fixed effects. Examples of the combined effect for 1990, 2007, 2013, and 2017. See Fig. S12 for temporal random effect alone in each year and Fig. S13 for spatial random effect alone.

*Figure S16. Estimated and interpolated cougar densities.*

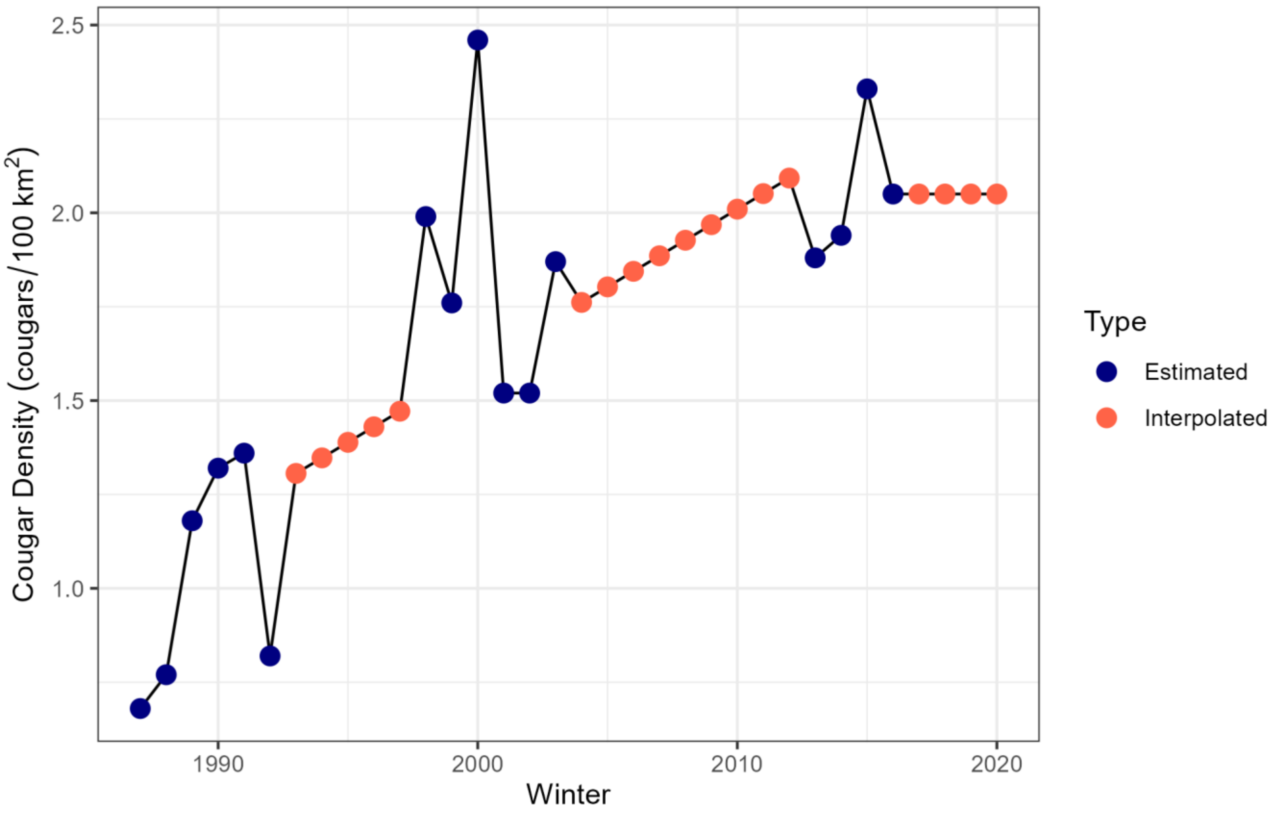


**Figure S16.** Cougar densities were estimated from field surveys in three phases: Phase 1 (1987 – 1993; Ruth *et al.* 2019), Phase 2 (1998 – 2004; Ruth *et al.* 2019), and Phase 3 (2014 – 2017; Anton 2020). Values between Phases 1 and 2 and between Phases 2 and 3 were linearly interpolated from the entire dataset. Values after Phase 3 (2018 – 2020) were assigned the mean of Phase 3.
